# Mechanism of membrane perforation in rotavirus cell entry

**DOI:** 10.64898/2026.01.21.700916

**Authors:** Marilina de Sautu, Conny Leistner, Tomas Kirchhausen, Simon Jenni, Stephen C. Harrison

## Abstract

Cell entry of non-enveloped, animal viruses requires translocation of a macromolecular assembly across a cellular membrane. Double-stranded RNA viruses introduce into the target cell an inner capsid particle that does not uncoat further. Instead, it extrudes capped viral mRNA by virtue of polymerase and capping activities within it. As described here, we used cryogenic electron tomography to visualize the full course of rhesus rotavirus entry, from cell attachment and virion uptake to release of the subviral particle. The cryo-tomograms and subtomogram averaging of classified subparticles link high-resolution structures of the virion and its components with time series from live-cell fluorescence microscopy. We outlined the mechanism of each step in the entry process, including the membrane perforation step that transfers a subviral particle into the cytosol.

---

Rotaviruses, so named for their wheel-like appearance in the electron microscope, are a group of non-enveloped viruses with segmented, double-stranded RNA (dsRNA) genomes (*1*). The outer layer of an infectious rotavirus particle (“triple-layer particle”: TLP) comprises two proteins – a spike-like asymmetric trimer (viral protein (VP) 4, magenta, red, orange and tan, in Fig. 1, A and B) and a Ca^2+^-stabilized trimer (VP7, yellow, in Fig. 1A) (*2-4*). The role of the outer layer is to deliver the double-layer particle (DLP) it surrounds into the cell to be infected. A maturation proteolytic cleavage processes VP4 into VP8* (magenta) and VP5* (red, orange, tan), with excision of a few residues at the junction and loss of all but the N-terminal 26 residues of VP8* from one of the three subunits (Fig. 1B). Within the DLP, VP1, the RNA-dependent, RNA polymerase (RdRp), associated 1:1 with each dsRNA segment, generates messages, capped by the activity of VP3, a guanylyl- and methyl-transferase, and extrudes them co-transcriptionally without further uncoating (*1, 5-9*). VP4, cleaved to VP8* and VP5*, mediates both attachment and uptake of a TLP. VP8* binds the receptor – sialic acid on glycosphingolipids, in the case of the rhesus rotavirus (RRV) used in the studies reported here, and blood-group antigens for certain human rotaviruses (*10, 11*). VP5* then engages the membrane through hydrophobic loops at the tip of its prominent β-barrel domain and undergoes a conformational change that inserts its C-terminal, “foot” domain into the target membrane (Fig. 1B) (*4*). These interactions drive membrane invagination, which for RRV entering BSC-1 cells is a largely self-driven, dynamin-independent, endocytic process (*12-14*).

**Fig. 1.**
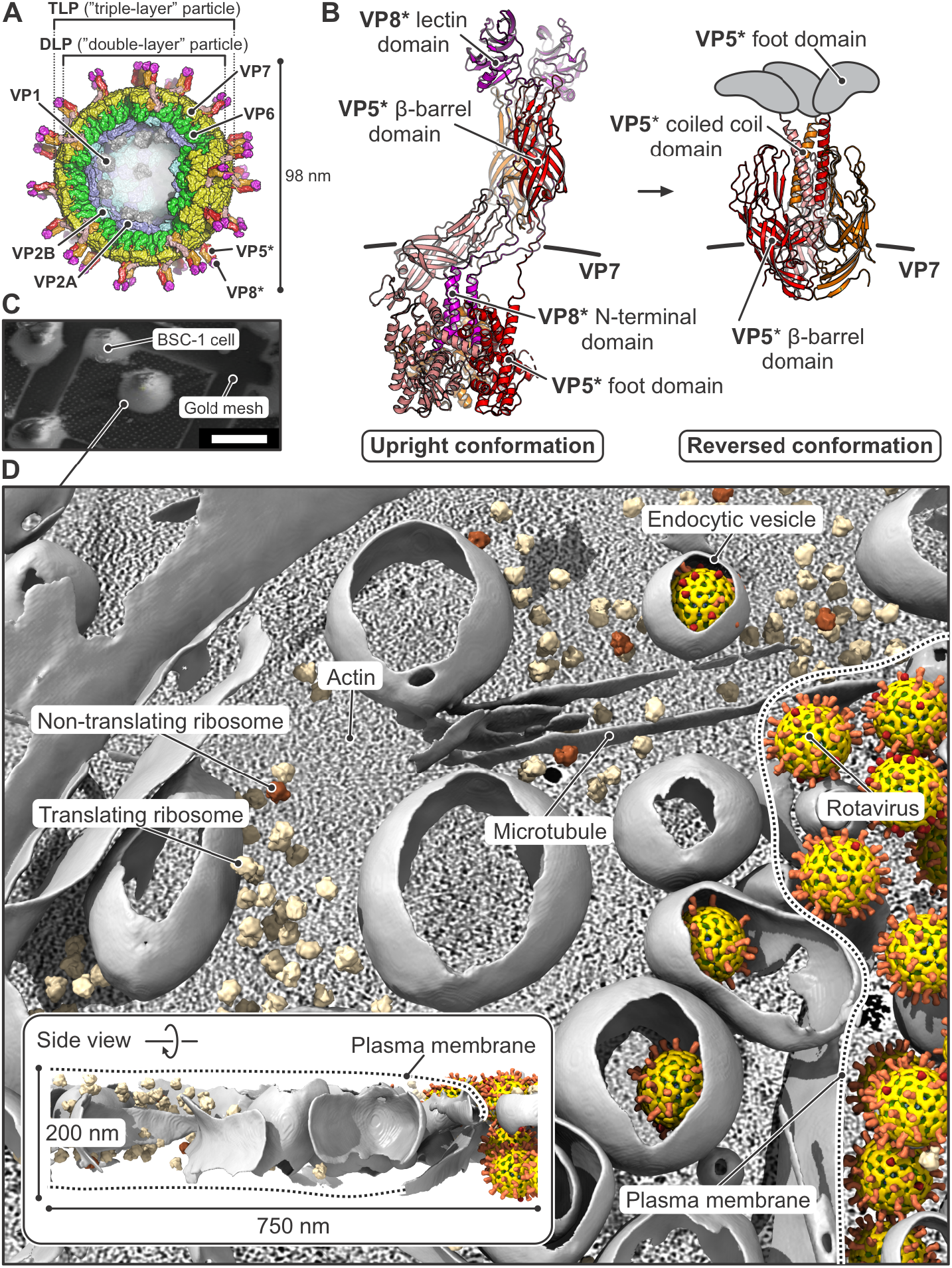
In situ visualization of rotavirus cell entry by cryo-ET. (**A**) Structure of a rotavirus particle, partially cut open to expose internal structures. Viral subunits are shown in surface representation. Viral protein (VP) 2 (VP2A and VP2B, in blue and cyan, respectively) and VP6 (green) form the shell of the “double-layer” particle (DLP), packaging the viral RNA, VP1 (gray, the capsid-bound RNA-dependent RNA polymerases [RdRp]), and VP3 (not shown, the RNA cap-generating enzyme). The outer layer of the “triple-layer” particle (TLP) contains VP7 (yellow) and VP4, proteolytically cleaved to VP8* (magenta) and VP5* (red, orange, and salmon). VP8* and VP5* asterisks indicate that these virion components are the results of proteolytic cleavage (of VP4) rather than primary gene products. (**B**) Upright (PDB-ID pdb_00009c1h) and reversed (PDB-ID pdb_00009c1j) conformations of VP4, after cleavage to VP8* (magenta) and VP5* (red, orange, salmon). Foot regions of reversed conformation shown schematically in gray. (**C**) Cryogenic scanning electron microscopy (cryo-SEM) image of BSC-1 cells deposited on the carbon film of a gold mesh and vitrified for tomography. (**D**) Molecular interpretation (segmentation) of a representative tomogram of a BSC-1 cell incubated with rotavirus. The plasma membrane is indicated by a dashed line. The inset shows a side view of the tomogram with its approximate dimensions. Anisotropy of the reconstructed tomogram limits membrane segmentation, giving rise to the “open” appearance of closed vesicles. We designated particles as fully engulfed in small vesicles only when we could detect known cytosolic components (actin filaments, microtubules, or ribosomes) in sections of the tomogram both above and below the vesicle in question. See Material and Methods for description of segmentation procedure. See movie S1 for the reconstructed tomogram and segmentation.

In experiments in which each of the components (VP4, VP7, DLP) was marked with a spectrally distinct fluorophore, we traced a trajectory from attachment to DLP delivery into the cytosol over a time course of no more than 10–15 min (*12, 15, 16*). An initial step after complete engulfment of a particle (i.e., pinching off of a virion-containing vesicle) is loss of Ca^2+^ from the particle and from the vesicle containing it (*15*). Permeabilization of the membrane to Ca^2+^ is a consequence of VP5* foot insertion, so that Ca^2+^ equilibrates with the cytosol once pinching off is complete (*13*). VP7, a Ca^2+^ stabilized trimer, then loses Ca^2+^ and dissociates from the particle (*3, 17*).

We describe here how VP7 dissociation leads to DLP delivery. We visualized the full entry pathway by cryogenic electron tomography (cryo-ET); the tomograms showed that VP7 dissociation correlated with opening of a pore in the vesicular membrane, allowing the DLP to escape into the cytosol. We confirmed this VP7 activity by experiments with liposomes in vitro. Our model for the activity – projection of a VP7 amphipathic helix shown previously to perforate liposomes as a peptide (*18, 19*) – resolves a longstanding puzzle and suggests a mechanism with potential significance for other non-enveloped viruses.

## Cryo-ET visualization of rotavirus entry

We recorded tilt series of rotavirus entering from the periphery of BSC-1 cells, which spread enough not to require further thinning by FIB-milling (Fig. 1C, fig. S1, and Methods). Our analysis included 838 tomograms from 7 separate experiments (datasets 1–7, table S1, and figs. S2 to S10; representative tomogram and segmentation in Fig. 1D and movie S1). We could describe the entry pathway in six steps (Fig. 2, A to F). To find entering particles in the small part of a cell seen in any single tomogram, we added a greater excess of TLPs than in our published live-cell imaging experiments (*12*). Nonetheless, we rarely found more than one or two particles in the same endocytic vesicle, so that the conclusions here correspond well to live-cell analyses, in which all traces had initial intensities corresponding to calibrated labeling for single virions (*12*).

**Fig. 2.**
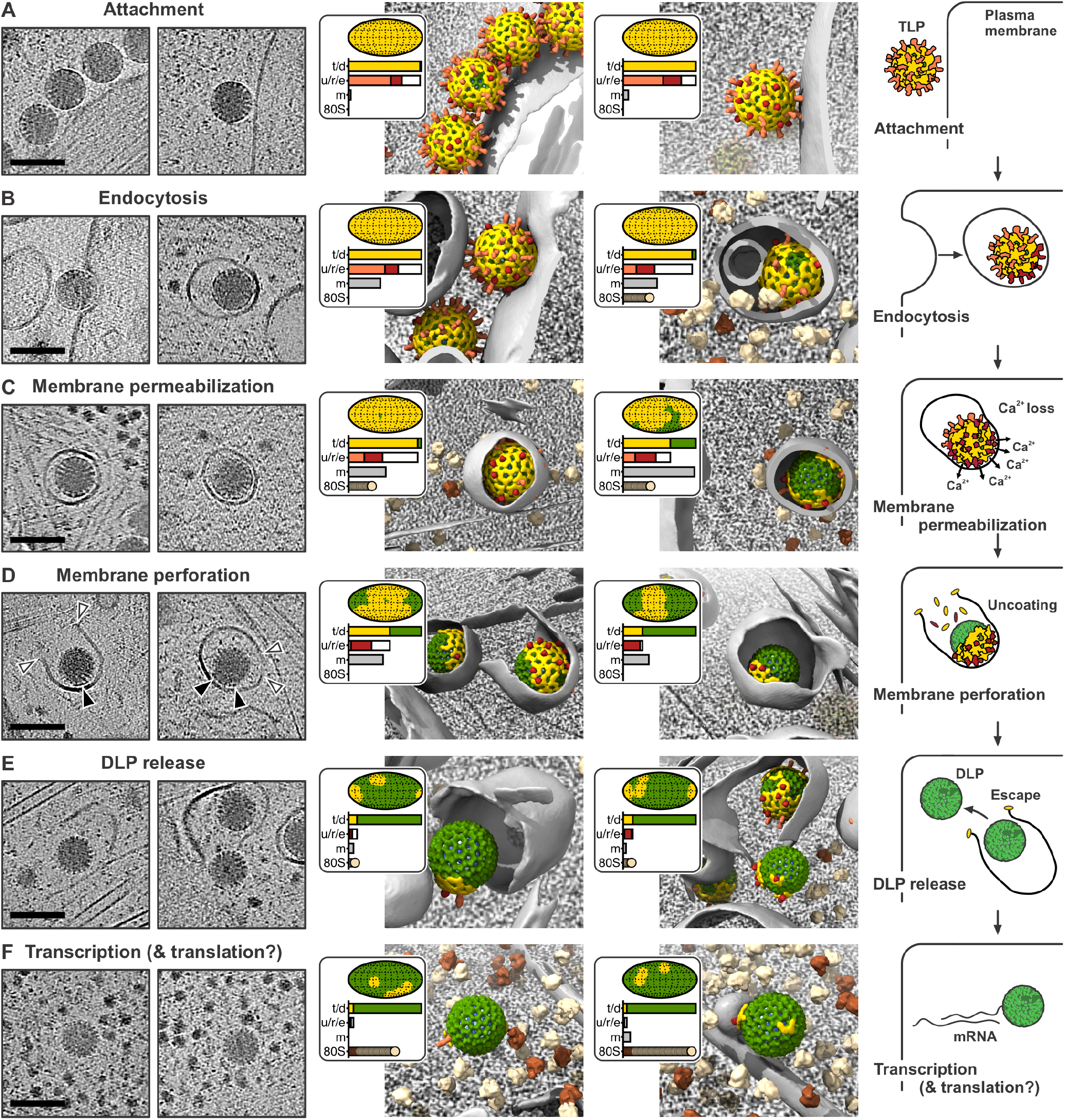
Rotavirus cell entry steps. Tomogram sections and interpretations based on subtomogram averaging, local classification of virus structure and conformations, segmenting cellular membranes, and identifying relevant cellular components. (**A** to **F**) ***Left-hand column***: two representative 10 nm-thick tomogram sections centered on particles of interest. Scale bars: 100 nm. ***Central column***: molecular interpretation of the two tomogram sections shown on the left. Particles are positioned and oriented as determined from the subtomogram analysis, with VP6 (green), VP7 (yellow), VP5*/VP8* in upright conformation (light red), and VP5* in reversed conformation (dark red). Segmented membranes are shown in gray, translating and non-translating ribosomes in beige and brown, respectively. For each particle, an inset shows the extent of VP7 uncoating on a viral surface projection and by a horizonal bar plot (t/d = TLP / DLP); the number and conformation of spikes in upright (light red) and reversed (dark red) conformation (u/r/e = upright / reversed / empty); the total volume of segmented membrane within a 16-nm sphere drawn at each spike position (m = membrane); and the number of ribosomes in the vicinity (radius of 150 nm from the center) of the particle (80S = ribosomes). See also movie S2 for membrane segmentation shown as a transparent surface. ***Right-hand column***: interpretative sketch. (A) Attachment of TLP to host cell. (B) Invagination of cell membrane and particle uptake. (C) Membrane permeabilization induced by VP5* bilayer interaction and conformational change from upright (light red) to reversed (dark red), resulting in loss of Ca^2+^ from the endocytic vesicle. (D) Membrane perforation by dissociated VP7. (E) DLP release into the cytosol. (F) Likely synthesis and extrusion of viral mRNA and potential translation of emerging transcripts.

RRV particles interact with cell membrane through their VP4 spikes in two distinct contact modes (*4, 12*). In the “loose” contact (Fig. 2A), the spike engages the membrane through the VP8* lectin domains.

In the “close” contact (Fig. 2B), which reflects the spike conformational transition from “upright” to “reversed” (Fig. 1B) (*4*), the VP5* hydrophobic loops contact the outer leaflet of the membrane bilayer, into which the foot domain inserts. Wrapping of membrane around the particle appeared in the tomograms to require the strong close contacts (Fig. 2B), and fully internalized particles usually had a large part of their surface in close contact with vesicular membrane (Fig. 2, B and C, and fig. S7F). At these contacts with reversed VP5*, Ca^2+^ will cross the membrane (*13*). We did not find clathrin or other coat-like structures around the wrapped membrane.

Once fully engulfed, the endocytic vesicle and TLP will lose Ca^2+^ as the ions pass from the lumen into the low-Ca^2+^ environment of the cytosol, causing VP7-bound Ca^2+^ ions to dissociate and the outer layer to disassemble. The tomograms showed that VP7 dissociation (“uncoating”) was on the side of the particle distal to the membrane-contacting surface; that is, uncoating initiated away from the site of attachment (Fig. 2, C and D). Even after extensive VP7 dissociation and membrane perforation, residual VP7 sometimes remained on one side of the largely uncoated particle (Fig. 2D, black arrows). Thus, despite generating a Ca^2+^ leak, insertion of reversed VP5* into the membrane at the close contact evidently stabilized the outer layer.

We detected various examples of membrane rupture with escaping DLPs (Fig. 2, D and E). From live-cell imaging experiments (*12, 15, 16*), we know that escaping particles diffuse rapidly away from the vesicle they leave, consistent with the relatively few examples of the exit step captured in our dataset. A single, opened, large aperture was observed in the vesicular membrane, rather than a general, multi-point dissolution of the bilayer. Density around the pore rim (e.g., white arrows in Fig. 2D) suggested that the scission event required stabilization of the exposed edges of the bilayer by the dissociated VP7.

DLPs that had escaped into the cytosol were often surrounded by clusters of ribosomes (Fig. 2F). Transcription from DLPs can initiate promptly after loss of the outer layer (*16*); the local ribosome clusters therefore suggest that polysomes might assemble on the emerging transcripts. A supervised classification procedure (Materials and Methods, fig. S10) indicated that over 80% of the ribosomes in DLP-adjacent clusters were in configurations associated with elongation (movie S3) (*20*), and we found a significantly greater number of ribosomes in the vicinity of free DLPs than in the neighborhoods of vesicle-enclosed rotavirus particles elsewhere in the tomograms (table S2). These observations are consistent with co-transcriptional translation, but do not yet establish it.

For each of the steps shown in Fig. 2, we determined the extent of VP7 uncoating, the ratio of upright to reversed spikes, their membrane proximity, and the results of classifying all ribosomes within 1500 Å of the particles assigned to that step (table S2). The statistics assembled in table s2, together with the subtomogram averaged density maps in fig. S6 derived from the selected particles assigned to each entry step, are in accord with the sequence of entry stages described above.

### VP5* membrane interaction

We generated a subtomogram average of membrane-bound VP5* in reversed-conformation spikes as outlined in Supplementary Text and diagrammed in fig. S2. The 3D subtomogram averaged map reconstructed in M (fig. S9) showed the VP5* β-barrel trimer, characteristic of the reversed conformation, with a central three-chain coiled-coil domain that extended through the membrane (Fig. 3A). The reversed-spike structure (PDB-ID pdb_00009c1j) could be docked confidently into the map.

**Fig. 3.**
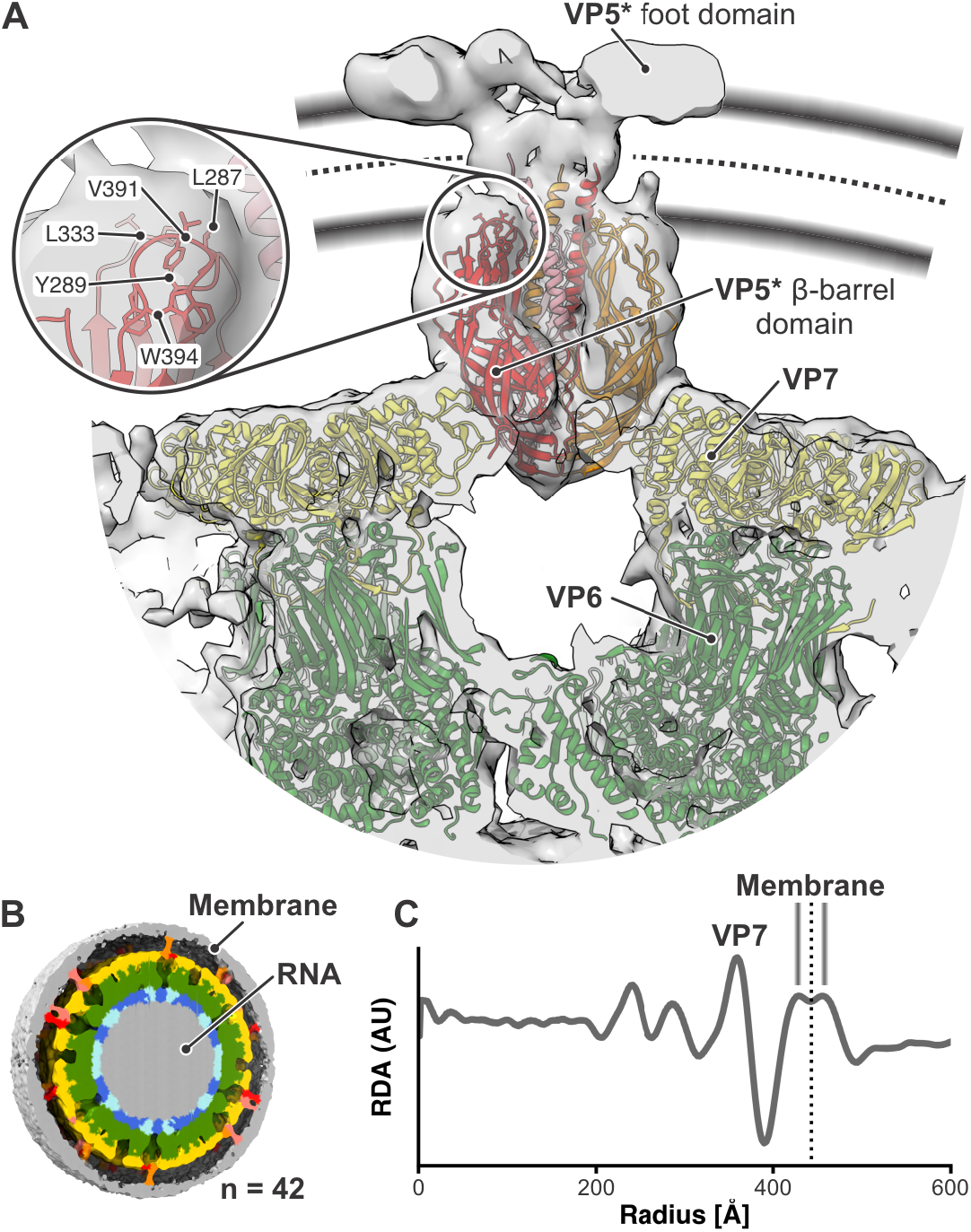
Membrane interaction of VP5*. (**A**) Structure of membrane-bound, reversed VP5*. Subtomogram average density of 3,255 reversed spikes obtained after classification (C_3_ symmetry imposed) in gray. The rotavirus reversed VP5* model (PDB-ID pdb_00009c1j), placed into the density, is colored as in Fig. 1. VP5* foot domain density is labeled. The middle of the membrane (dotted line), as determined from the radial density profile shown in (C). The thickness of the membrane is shown by the approximate position of the lipid head groups (gray bands). The close-up shows the hydrophobic loops at the tip of the β-barrel domains with amino acid residues labeled. (**B**) Icosahedrally averaged density from 42 rotavirus particle subtomograms for which most spikes were in close contact with membrane. RNA, gray; VP2A, VP2B, blue and cyan, respectively; VP6, green; VP7 yellow; VP5*, red, orange, and salmon; membrane, gray. (**C**) Radial density profile (RDA) in arbitrary units (AU), calculated from 42 rotavirus subtomograms averaged to generate the figure in (B) (see Materials and Methods for details). The peak corresponding to the VP7 layer is labeled. Membrane appears as a double peak. The middle of the lipid bilayer is 443 Å from the center of the virus and marked with a dotted line.

To determine the membrane position surrounding the virion, we calculated a rotational density average from 42 viruses with high membrane overlap and close contact with vesicular membrane over most of their surface (Fig. 3, B and C). We estimated from this average the membrane boundaries shown in Fig. 3A. The tips of the VP5* hydrophobic loops insert into the proximal leaflet of the bilayer, and the three-chain coiled-coil extends across the bilayer’s hydrophobic core (Fig. 3A). High-resolution structures show that the distal part of the coiled-coil in the reversed conformation includes most of the initial α helix of the foot domain in the upright conformation (*2, 4, 21*). Three globular features that likely represent the rest of the foot splay symmetrically away from the coiled-coil at a level corresponding to the cytosolic-side polar headgroups of the membrane bilayer (Fig. 3A).

Uniform close contacts between entering TLPs and the invaginating plasma membrane show that the coiled-coil and foot of VP5* insert into the membrane as inward budding proceeds, thus generating the work needed to distort the bilayer and wrap it tightly around the particle (Fig. 2B). The same membrane insertion also results in Ca^2+^ permeability (*13*), and we therefore expect Ca^2+^ to enter the cell from the medium even before engulfment is complete. Imaging of cells transfected with GCaMP6s-CAAX, a plasma membrane targeted Ca^2+^ sensor (*22*), indeed showed Ca^2+^ concentration transients during short incubation with rotavirus particles (fig. S11).

In the vesicle pinched off from the wrapped membrane, close contacts are present over at least one side of the TLP but the distal surface generally faces a small volume of enclosed medium (Fig. 2, B and C). This property of the budded vesicles suggests that pinching off requires some cellular mechanical activity – most likely local mobilization of cortical actin, as indicated by reduced efficiency of infection in the presence of latrunculin A (fig. S12A). One potential functional consequence of Ca^2+^ permeation as the vesicle forms is reorganization of actin close to the entering TLP, by Arp2/3 activation as a downstream consequence of Ca^2+^ signaling. Actin at sites of entry forms branched networks surrounding the viral invaginations (fig. S12, B to H), with the geometry expected of Arp-2,3-mediated nucleation (fig. S12I).

### Liposome disruption by VP7 monomer

We detected VP7 dissociation by loss of the thin, wagon-wheel rim of the particle (Fig. 2D). Extensive uncoating correlated with membrane rupture and particle escape into the cytosol (Fig. 2D). These observations suggested that monomeric VP7, dissociated from virus particles because of Ca^2+^ depletion within the vesicle, had perforated the membrane and facilitated particle escape. We tested whether VP7 monomers or trimers, without contribution of any cellular factor, could disrupt a liposome membrane and induce release of an enclosed carboxyfluorescein (CF) dye (Fig. 4A and Methods). The lipid composition was similar to that of mammalian plasma membranes. Time-lapse images showed that in the presence of VP7 monomer in EDTA buffer, the CF signal often disappeared abruptly while the lipid signal persisted, indicating membrane perforation that had allowed dye to leak from the liposomes (Fig. 4B). At 30 min after adding VP7 monomer, about a third of the liposomes had lost more than 50% of their initial CF intensity; neither VP7 trimer in Ca^2+^ buffer nor EDTA buffer alone yielded any significant CF signal loss (Fig. 4C and fig. S13). Incomplete loss of CF indicated either a multilamellar liposome or two unresolved, adjacent liposomes. Cryo-EM images of the same liposome preparations showed localized perforations (Fig. 4D and fig. S14) that recapitulated the site-specific vesicle disruption patterns observed in infected cells by cryo-ET (Fig. 2, D and E, and Fig. 4E). Tilt-series acquisition and tomographic reconstruction of liposomes exposed to monomeric (i.e., Ca^2+^ free) VP7 likewise showed large, well-defined, single pores in the bilayer, with associated densities around the rim of the opening consistent with bound protein (movie S4).

**Fig. 4.**
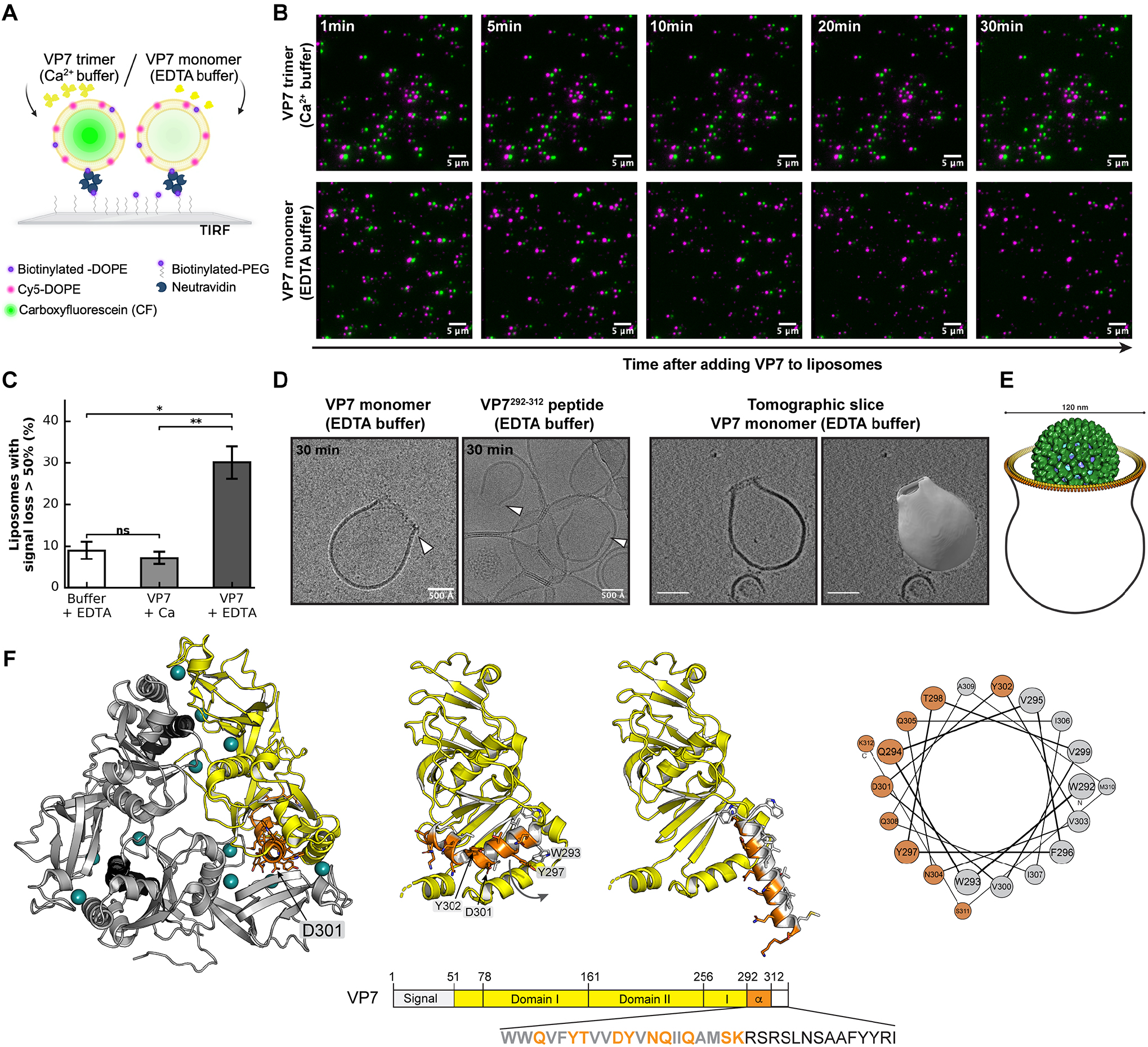
Membrane perforation by VP7 monomer. (**A**) Schematic of the total internal reflection fluorescence (TIRF) microscopy experimental set-up: carboxyfluorescein (CF, green)-containing liposomes with biotinylated-DOPE and Cy5-DOPE (magenta) are tethered to a biotinylated-PEG coated coverslip by neutravidin. (**B**) Time-lapse TIRF microscopy imaging of liposomes incubated with VP7 trimer (Ca^2+^ buffer, top row) or VP7 monomer (EDTA buffer, bottom row) and imaged at indicated time points. CF-channel is shifted 7 pixels to the right from lipid-channel for better colocalization visualization. (**C**) Quantification of liposome permeabilization showing the percentage of liposomes with signal loss of more than 50% after 30 min incubation with buffer only (white), VP7 trimer (light grey) or VP7 monomer (dark grey). Error bars represent the standard deviation of the mean (SD, n = 3 independent experiments); t-test statistical analysis was used between samples. *, p < 0.05; **, p < 0.01; ns, not significant. (**D**) Cryo-EM images (left) of perforated liposomes after 30 min incubation with VP7 monomer or VP7^292–312^ peptide and slice through a cryo-ET reconstruction together with segmentation (right) of a VP7-perforated liposome. (**E**) Schematic diagram, based on Fig. 2, D and E, representing perforations seen in viral uptake vesicles, for comparison with the images in (D) of liposomes incubated with VP7 monomer. (**F**) VP7 trimer (PDB-ID pdb_00009c1g (*3*)) highlighting in orange the C-terminal amphipathic helix (residues 292–312). The VP7 monomer (yellow, center) shows domains I and II, with the C-terminal helix extending from the bipartite globular body of the protein. Helical wheel projection of residues 292–312, showing segregation of hydrophobic (gray) and hydrophilic (orange) residues. The sequence below shows the amphipathic region (orange) and adjacent residues at the C-terminal end of the polypeptide chain. [Panel (A) created with BioRender.com]

Early, unpublished experiments, using a bulk assay to monitor liposome disruption by VP7, showed activity from a peptide close to the C terminus of the VP7 polypeptide chain, W292–K312 (Fig. 4F), released from the protein by adventitious proteolytic cleavage (*18*). That assay did not, however, detect disruption by intact, uncleaved VP7 monomer. Another group later described liposome disruption by a similar, 21-residue, synthetic peptide (Q280–M310) (*19*). As a specific precaution to avoid proteolysis, we added trypsin inhibitor to the single liposome experiments described above.

In the VP7 trimer, residues W292–K312 (orange, in Fig. 4F) form a kinked helix at the Ca^2+^-dependent interface, with one of the Ca^2+^ ligands (D310) near its C terminus. If straightened, W292–K312, would form a perfectly amphipathic, ∼30 Å-long α helix. We suggest that the kinked helix can unfold and straighten when not constrained by its Ca^2+^-stabilized contacts in the trimer (Fig. 4F). The proteolytic sensitivity of the R291-W292 peptide bond, when the trimer dissociates, exposes the interface, and liberates the helix in question, is evidence for flexibility near R291 (Fig. 4F). In cell entry, the high local concentration of monomeric VP7 in the vesicle will favor interaction of this amphipathic helix with the luminal membrane leaflet of the vesicle. The synthetic VP7 peptide (VP7^292-312^) produced localized perforations in single liposomes very similar to those observed by intact VP7 monomer (Fig. 4D), consistent with a mechanism in which the corresponding segment projects from the folded VP7 globular structure.

Rotavirus infection in vertebrate hosts occurs in the trypsin-rich lumen of the proximal gut. We considered whether inclusion of traces of trypsin within the endocytic vesicle generated by the entering virus might release the membrane-disrupting, C-terminal peptide from monomeric VP7 dissociated from the Ca^2+^-depleted particle. We carried out an infectivity assay in the presence of either small amounts of trypsin or of trypsin inhibitor and found no significant infectivity difference (fig. S15). Thus, release by tryptic cleavage of the amphipathic VP7 peptide is not required for rotavirus infection to proceed.

## Discussion

Our results lead to the following inferences, consistent with the kinetics established by live-cell imaging (*12, 15, 16*). (1) Binding of RRV TLPs at the surface of BSC-1 cells relies on the upright conformation of VP4 (“loose” contacts with the cell surface), but invagination and final engulfment rely on the reversed conformation (“close” contacts). Insertion of the C-terminal part of VP5* (coiled-coil and foot) into the plasma-membrane bilayer, generating the close contacts, wraps membrane around the particle. We detect no features that might be ascribed to a protein receptor. (2) VP5* insertion allows Ca^2+^ ions to enter the cell (*13*), leading to downstream Ca^2+^ signaling and likely to local reorganization of the actin cytoskeleton, facilitating closure of the vesicle that surrounds the entering TLP. (3) Vesicle closure allows depletion of Ca^2+^ within it, as there is no longer replenishment from the medium, initiating VP7 dissociation. (4) Dissociation is distal to the close membrane contacts, which appear to stabilize the VP7 layer, even though the Ca^2+^ flux is through the reversed VP5* spikes within the close contact. (5) Dissociated VP7 monomers are therefore the presumptive agents of membrane perforation. (6) In vitro, monomeric VP7 disrupts liposomes and forms single, large pores that closely resemble the DLP exit pores captured by cryo-ET. A peptide spanning residues 292–312 of VP7 does the same. These observations show that VP7 is sufficient to generate the observed openings. Although from our cellular cryo-ET alone we cannot rule out some participation by VP5*, released along with VP7, in vesicle perforation, the in vitro results make such participation unlikely. (7) Ribosomes often surround the DLPs released into the cytosol, suggesting translation of nascent viral mRNA as it exits the transcriptionally active DLP.

The sequence corresponding to VP7 residues 292–312 in RRV is conserved among rotavirus A strains; in the other groups, it retains both amphipathicity and a potential aspartate Ca^2+^ ligand, despite somewhat greater variability. We note that rotavirus entry through clathrin-mediated endocytosis might in principle be an infectious pathway in other cell types or for other rotavirus strains. Clathrin dissociates within a minute of pinching off (*23*), and the resulting uncoated vesicle would be nearly indistinguishable from the small vesicles from which RRV DLPs penetrate in BSC-1 cells. VP5* insertion could occur either during coated pit assembly or after clathrin uncoating. Ca^2+^ loss, and VP7 dissociation would then follow as seen here.

The entry mechanism just described includes two membrane-insertion events. The first accompanies the upright-to-reversed conformational transition of VP5*; the second is perforation by VP7. VP5* insertion requires the VP8* lectin domains to dissociate from the hydrophobic tips of the VP5* β barrels (*24*). The dissociation can be spontaneous, as the interface between VP8* and VP5* is small, but VP8* cannot fully dissociate because of the tether that connects the lectin domain to the VP8* N terminus, anchored in the foot (*4*). That is, the lectin domain is probably always in an “on-off” equilibrium, and when not attached to membrane, it will reassociate promptly. When bound to a membrane-glycolipid headgroup, however, its dissociation can allow the hydrophobic tips of the VP5* β barrels to engage the membrane, preventing reassociation and initiating the upright-to-reversed conformational change. Zippering up of the central coiled-coil will extract the unfolding foot from beneath the VP7 layer and project it through the bilayer. The model for membrane-inserted VP5* we obtained by sub-tomogram averaging represents the result of this transition. Although of limited resolution, the model suggests a potential axial channel through which Ca^2+^ might pass.

The mechanism of DLP release visualized by cryo-ET is consistent with experiments showing that crosslinking of VP7 subunits within a trimer, either by divalent antibodies or by mutationally introduced disulfides, inhibits infection and prevents concomitant entry of the toxin, α-sarcin (*25*). Because DLP transcriptional activity requires loss of VP7 (*26*), those observations did not by themselves imply that VP7 has perforating activity. It is now clear that loss of Ca^2+^, which causes VP7 to dissociate from the particle, leads both to activation of the DLP as a transcriptase and to generation of the VP7 monomer as a pore-former.

The VP7-generated pores in our cryo-tomograms are strikingly similar to those VP7 generates on liposomes *in vitro*. There is always a single pore per vesicle/liposome, of variable size *in vitro* and probably in cells; the strong density at the edges of the pores shows that many copies of VP7 line the rim; the membrane bilayer curves outward at the rim of the pore, so that the plane of the bilayer edge is roughly normal to the pore axis. A single pore per vesicle/liposome indicates cooperativity; the variable size suggests an initiation-growth mechanism (see Supplementary text). To cover the rim of a pore 100– 120 nm in diameter with close contacts between helices – roughly the size of the perforations we see in the tomograms – would require 300–350 helices – i.e., participation of less than half of the 780 total VP7 monomers on a rotavirus particle. Thus, the available VP7 can readily form the observed openings.

Release of a pore-forming activity from the virion, triggered by change in a specific environmental variable (e.g., Ca^2+^ concentration or pH), may also describe the entry mechanism of other non-enveloped viruses that deliver a modified virion or subviral particle into the cytosol. For example, an amphipathic helix at the N terminus of adenovirus protein VI is essential for translocation of an infecting adenovirus particle from an early endosome into the cytosol (*27*). Loss of pentons and peripentonal hexons, induced by endosomal low pH, allows release of pVI from the modified virion (*28-30*). For any pore-forming mechanism, the number of released, pore-forming molecules (e.g., VP7 or, hypothetically, pVI) must be sufficient to form an opening of diameter large enough to release the subviral particle. Design of non-enveloped delivery vehicles for macromolecular cargo might be able to take advantage of a similar mechanism.

## Supporting information

data file (membrane overlap)

## ACKNOWLEDGMENTS

We thank Shaun Rawson, Isobel Garrett and the staff at the Harvard Medical School Center for Cryo-Electron Microscopy in Structural Biology, for expert advice and assistance, and Eric Marino, Gustavo Scanavachi and members of the Kirchhausen laboratory (Boston Children’s Hospital), for assistance with fluorescence microscopy.

## Author contributions

Conceptualization: all authors; Methodology: all authors; Investigation: M.d.S., C.L., S.J.; Visualization: M.d.S., S.J.; Writing – original draft: M.d.S., S.J., S.C.H.; Writing – review and editing: all authors.

## Data, code, and materials availability

The cryo-ET maps from subtomogram averaging are deposited in the Electron Microscopy Data Bank with accession identifiers EMD-75185 (icosahedral virus reconstruction from dataset 2, fig. S3) and EMD-75186 (membrane-bound, reversed VP5* trimer, classification 3 with a tight mask, class 1, Fig. 3A and fig. S9). Atomic coordinates are deposited in the Protein Data Bank with accession identifiers PDB-ID pdb_000010ic (icosahedral virus) and PDB-ID pdb_000010id (reversed VP5* trimer). Data analysis C shell, bash, Python, and MATLAB scripts used for this study are available the Zenodo data repository (*31*). All other data are available in the manuscript or the supplementary material. This study did not generate new materials.

## SUPPLEMENTARY MATERIALS

Materials and Methods

Supplementary Text

Figs. S1 to S15

Tables S1 to S2

References (*32*–*65*)

Movies S1 to S4

Data S1

## REFERENCES AND NOTES

1. S. Crawford, S. Ding, H. B. Greenberg, M. K. Estes, in Fields Virology, 7th ed., P. M. Howley, D. M. Knipe, Eds. (Wolters Kluyver, Philadelphia, PA, 2023), vol. 3, pp. 362–413.

2. E. C. Settembre, J. Z. Chen, P. R. Dormitzer, N. Grigorieff, S. C. Harrison, Atomic model of an infectious rotavirus particle. EMBO J. 30, 408–416 (2011).

3. S. T. Aoki, E. C. Settembre, S. D. Trask, H. B. Greenberg, S. C. Harrison, P. R. Dormitzer, Structure of rotavirus outer-layer protein VP7 bound with a neutralizing Fab. Science 324, 1444–1447 (2009).

4. T. Herrmann et al., Functional refolding of the penetration protein on a non-enveloped virus. Nature 590, 666–670 (2021).

5. X. Lu et al., Mechanism for coordinated RNA packaging and genome replication by rotavirus polymerase VP1. Structure 16, 1678–1688 (2008).

6. N. M. Bartlett, S. C. Gillies, S. Bullivant, A. R. Bellamy, Electron microscopy study of reovirus reaction cores. J. Virol. 14, 315–326 (1974).

7. J. A. Lawton, M. K. Estes, B. V. Prasad, Three-dimensional visualization of mRNA release from actively transcribing rotavirus particles. Nat. Struct. Biol. 4, 118–121 (1997).

8. S. Jenni et al., In situ Structure of Rotavirus VP1 RNA-Dependent RNA Polymerase. J. Mol. Biol. 431, 3124–3138 (2019).

9. K. Ding et al., In situ structures of rotavirus polymerase in action and mechanism of mRNA transcription and release. Nat. Commun. 10, 2216 (2019).

10. P. R. Dormitzer, Z. Y. Sun, O. Blixt, J. C. Paulson, G. Wagner, S. C. Harrison, Specificity and affinity of sialic acid binding by the rhesus rotavirus VP8* core. J. Virol. 76, 10512–10517 (2002).

11. L. Hu et al., Cell attachment protein VP8* of a human rotavirus specifically interacts with A-type histo-blood group antigen. Nature 485, 256–259 (2012).

12. A. H. Abdelhakim et al., Structural correlates of rotavirus cell entry. PLoS Pathog. 10, e1004355 (2014).

13. M. de Sautu, T. Herrmann, G. Scanavachi, S. Jenni, S. C. Harrison, The rotavirus VP5*/VP8* conformational transition permeabilizes membranes to Ca2+. PLoS Pathog. 20, e1011750 (2024).

14. K. T. Kaljot, R. D. Shaw, D. H. Rubin, H. B. Greenberg, Infectious rotavirus enters cells by direct cell membrane penetration, not by endocytosis. J. Virol. 62, 1136–1144 (1988).

15. E. N. Salgado, B. Garcia Rodriguez, N. Narayanaswamy, Y. Krishnan, S. C. Harrison, Visualization of Calcium Ion Loss from Rotavirus during Cell Entry. J. Virol. 92, e01327–01318 (2018).

16. E. N. Salgado, S. Upadhyayula, S. C. Harrison, Single-particle detection of transcription following rotavirus entry. J. Virol. 91, e00651–00617 (2017).

17. P. R. Dormitzer, H. B. Greenberg, S. C. Harrison, Purified recombinant rotavirus VP7 forms soluble, calcium-dependent trimers. Virology 277, 420–428 (2000).

18. S. D. Trask, The Mechanism of Membrane Penetration by Rotavirus. Ph.D. thesis, Harvard University, Boston (2008).

19. S. Elaid et al., A peptide derived from the rotavirus outer capsid protein VP7 permeabilizes artificial membranes. Biochim. Biophys. Acta 1838, 2026–2035 (2014).

20. W. Zheng et al., Visualizing the translation landscape in human cells at high resolution. Nat. Commun. 16, 10757 (2025).

21. P. R. Dormitzer, E. B. Nason, B. V. Prasad, S. C. Harrison, Structural rearrangements in the membrane penetration protein of a non-enveloped virus. Nature 430, 1053–1058 (2004).

22. F. C. Tsai et al., A polarized Ca2+, diacylglycerol and STIM1 signalling system regulates directed cell migration. Nat. Cell Biol. 16, 133–144 (2014).

23. R. H. Massol, W. Boll, A. M. Griffin, T. Kirchhausen, A burst of auxilin recruitment determines the onset of clathrin-coated vesicle uncoating. Proc. Natl. Acad. Sci. U. S. A. 103, 10265–10270 (2006).

24. S. Jenni et al., Rotavirus VP4 Epitope of a Broadly Neutralizing Human Antibody Defined by Its Structure Bound with an Attenuated-Strain Virion. J. Virol. 96, e0062722 (2022).

25. S. T. Aoki, S. D. Trask, B. S. Coulson, H. B. Greenberg, P. R. Dormitzer, S. C. Harrison, Cross-linking of rotavirus outer capsid protein VP7 by antibodies or disulfides inhibits viral entry. J. Virol. 85, 10509–10517 (2011).

26. J. Cohen, J. Laporte, A. Charpilienne, R. Scherrer, Activation of rotavirus RNA polymerase by calcium chelation. Arch. Virol. 60, 177–186 (1979).

27. C. M. Wiethoff, G. R. Nemerow, Adenovirus membrane penetration: Tickling the tail of a sleeping dragon. Virology 479-480, 591–599 (2015).

28. U. F. Greber, M. Willetts, P. Webster, A. Helenius, Stepwise dismantling of adenovirus 2 during entry into cells. Cell 75, 477–486 (1993).

29. O. Maier, D. L. Galan, H. Wodrich, C. M. Wiethoff, An N-terminal domain of adenovirus protein VI fragments membranes by inducing positive membrane curvature. Virology 402, 11–19 (2010).

30. C. L. Moyer, C. M. Wiethoff, O. Maier, J. G. Smith, G. R. Nemerow, Functional genetic and biophysical analyses of membrane disruption by human adenovirus. J. Virol. 85, 2631–2641 (2011).

31. M. de Sautu, C. Leistner, T. Kirchhausen, S. Jenni, S. C. Harrison, Data for: Mechanism of membrane perforation in rotavirus cell entry, Zenodo (2026); 10.5281/zenodo.21242795.

32. H. B. Greenberg et al., Production and preliminary characterization of monoclonal antibodies directed at two surface proteins of rhesus rotavirus. J. Virol. 47, 267–275 (1983).

33. W. J. H. Hagen, W. Wan, J. A. G. Briggs, Implementation of a cryo-electron tomography tilt-scheme optimized for high resolution subtomogram averaging. J. Struct. Biol. 197, 191–198 (2017).

34. D. Tegunov, P. Cramer, Real-time cryo-electron microscopy data preprocessing with Warp. Nat. Methods 16, 1146–1152 (2019).

35. D. N. Mastronarde, S. R. Held, Automated tilt series alignment and tomographic reconstruction in IMOD. J. Struct. Biol. 197, 102–113 (2017).

36. S. Zheng et al., AreTomo: An integrated software package for automated marker-free, motion-corrected cryo-electron tomographic alignment and reconstruction. J. Struct. Biol. X 6, 100068 (2022).

37. M. Chen, J. M. Bell, X. Shi, S. Y. Sun, Z. Wang, S. J. Ludtke, A complete data processing workflow for cryo-ET and subtomogram averaging. Nat. Methods 16, 1161–1168 (2019).

38. S. H. Scheres, RELION: implementation of a Bayesian approach to cryo-EM structure determination. J. Struct. Biol. 180, 519–530 (2012).

39. D. Tegunov, L. Xue, C. Dienemann, P. Cramer, J. Mahamid, Multi-particle cryo-EM refinement with M visualizes ribosome-antibiotic complex at 3.5 A in cells. Nat. Methods 18, 186–193 (2021).

40. D. Castano-Diez, M. Kudryashev, M. Arheit, H. Stahlberg, Dynamo: a flexible, user-friendly development tool for subtomogram averaging of cryo-EM data in high-performance computing environments. J. Struct. Biol. 178, 139–151 (2012).

41. J. Agirre et al., The CCP4 suite: integrative software for macromolecular crystallography. Acta Crystallogr. D Struct. Biol. 79, 449–461 (2023).

42. L. Lamm et al., MemBrain: A deep learning-aided pipeline for detection of membrane proteins in Cryo-electron tomograms. Comput. Methods Programs Biomed. 224, 106990 (2022).

43. E. F. Pettersen et al., UCSF ChimeraX: Structure visualization for researchers, educators, and developers. Protein Sci. 30, 70–82 (2021).

44. C. R. Harris et al., Array programming with NumPy. Nature 585, 357–362 (2020).

45. P. Virtanen et al., SciPy 1.0: fundamental algorithms for scientific computing in Python. Nat. Methods 17, 261–272 (2020).

46. M. G. F. Last, L. Abendstein, L. M. Voortman, T. H. Sharp, Ais: streamlining segmentation of cryo-electron tomography datasets. Elife 13, 1–32 (2024).

47. P. V. Afonine et al., Real-space refinement in PHENIX for cryo-EM and crystallography. Acta Crystallogr. D Struct. Biol. 74, 531–544 (2018).

48. J. M. Bell, M. Chen, P. R. Baldwin, S. J. Ludtke, High resolution single particle refinement in EMAN2.1. Methods 100, 25–34 (2016).

49. M. Kyoung, Y. Zhang, J. Diao, S. Chu, A. T. Brunger, Studying calcium-triggered vesicle fusion in a single vesicle-vesicle content and lipid-mixing system. Nat. Protoc. 8, 1–16 (2013).

50. E. Cocucci, F. Aguet, S. Boulant, T. Kirchhausen, The first five seconds in the life of a clathrin-coated pit. Cell 150, 495–507 (2012).

51. D. N. Mastronarde, SerialEM: A Program for Automated Tilt Series Acquisition on Tecnai Microscopes Using Prediction of Specimen Position. Microsc. Microanal. 9, 1182–1183 (2003).

52. S. D. Trask, P. R. Dormitzer, Assembly of highly infectious rotavirus particles recoated with recombinant outer capsid proteins. J. Virol. 80, 11293–11304 (2006).

53. J. Schindelin et al., Fiji: an open-source platform for biological-image analysis. Nat. Methods 9, 676–682 (2012).

54. J. D. Hunter, Matplotlib: A 2D graphics environment. Comput. Sci. Eng. 9, 90–95 (2007).

55. J. G. Galaz-Montoya, J. Flanagan, M. F. Schmid, S. J. Ludtke, Single particle tomography in EMAN2. J. Struct. Biol. 190, 279–290 (2015).

56. A. Burt et al., An image processing pipeline for electron cryo-tomography in RELION-5. FEBS Open Bio. 14, 1788–1804 (2024).

57. L. Lamm et al., MemBrain v2: an end-to-end tool for the analysis of membranes in cryo-electron tomography. bioRxiv, 2024.2001.2005.574336 (2025).

58. J. Rossjohn, G. Polekhina, S. C. Feil, C. J. Morton, R. K. Tweten, M. W. Parker, Structures of perfringolysin O suggest a pathway for activation of cholesterol-dependent cytolysins. J. Mol. Biol. 367, 1227–1236 (2007).

59. Y. Shai, Mechanism of the binding, insertion and destabilization of phospholipid bilayer membranes by alpha-helical antimicrobial and cell non-selective membrane-lytic peptides. Biochim. Biophys. Acta 1462, 55–70 (1999).

60. A. Niitsu et al., Rational Design Principles for De Novo alpha-Helical Peptide Barrels with Dynamic Conductive Channels. J. Am. Chem. Soc. 147, 11741–11753 (2025).

61. L. Yang, T. A. Harroun, T. M. Weiss, L. Ding, H. W. Huang, Barrel-stave model or toroidal model? A case study on melittin pores. Biophys. J. 81, 1475–1485 (2001).

62. L. A. Gillies, H. Du, B. Peters, C. M. Knudson, D. D. Newmeyer, T. Kuwana, Visual and functional demonstration of growing Bax-induced pores in mitochondrial outer membranes. Mol. Biol. Cell 26, 339–349 (2015).

63. T. Kuwana, N. H. Olson, W. B. Kiosses, B. Peters, D. D. Newmeyer, Pro-apoptotic Bax molecules densely populate the edges of membrane pores. Sci. Rep. 6, 27299 (2016).

64. Y. Zhang et al., Structural basis of BAX pore formation. Science 388, eadv4314 (2025).

65. P. E. Czabotar, A. J. Garcia-Saez, Mechanisms of BCL-2 family proteins in mitochondrial apoptosis. Nat. Rev. Mol. Cell Biol. 24, 732–748 (2023).

