## Supplemental text, Methods,figures and tables for "Mechanism of membrane perforation in rotavirus cell entry"

**The PDF file includes:**

Materials and Methods
Supplementary Text
Figs. S1 to S15
Tables S1 to S2
References (*32–65*)

**Other Supplementary Materials for this manuscript include the following:**

Movies S1 to S4
Data S1

#### Materials and Methods

##### Cells, plasmids and constructs

MA104 cells (ATCC) were grown in medium 199 Earle's salts (M199, Thermo Fisher Scientific) supplemented with 25 mM 4-(2-hydroxyethyl)-1-piperazineethanesulfonic acid (HEPES) and 10% Hi-FBS (Thermo Fisher Scientific). BSC-1 cells (ATTC) were grown in Dulbecco's Modified Eagle Medium (DMEM) with GlutaMAX (Thermo Fisher Scientific) supplemented with 10% Hi-FBS (Thermo Fisher Scientific). For VP7 expression, we cloned the full-length VP7 genomic sequence (G3 serotype, NCBI: txid444185) into a pFastbac expression vector, which we transformed into *E. coli* DH10α cells. For bacmid generation, VP7 vectors were transformed into DH10-Bac cells (Thermo Fisher Scientific) and plated onto LB-agar plates supplemented with 50 μg/ml kanamycin, 7 μg/ml gentamycin, 10 μg/ml tetracycline, 100 μg/ml blue-gal and 40 μg/ml β-D-1-thiogalactopyranoside (IPTG).

##### Reagent preparation

*Purification of rotavirus TLPs* – MA104 cells were grown to confluency in 850 $cm^2$ roller bottles in M199 supplemented with 10% FBS and 25 mM HEPES. Cells were infected with rhesus rotavirus (RRV, strain G3P5B[3], biosafety level 2, approved by the Committee on Microbiological Safety [COMS] at Harvard Medical School) at a multiplicity of infection (MOI) of 0.1 with 5 μg/ml of trypsin in the medium. After incubation at 37 °C for 24 h, the medium containing cells and debris was collected and frozen at -80 °C. After thawing, cells and debris were pelleted by centrifugation in a Beckman Coulter rotor JS 4.2 at 2,900 g for 30 min at 4 °C. Supernatant was then removed and the pellet resuspended in 1 ml ice cold TNC buffer (20 mM Tris, 100 mM NaCl, 1 mM $CaCl_2$, pH 8.0). Virus particles were concentrated from the supernatant by ultracentrifugation in a Beckman Coulter rotor Ti-45 rotor at 224,500 g. The supernatant was then discarded, and the pellet was resuspended in 1–3 ml ice cold TNC buffer and combined with the cell-debris fraction from the first centrifugation. The suspension was transferred to a 15 ml conical tube, and TNC was added to a final volume of 4 ml. The suspension was mixed with 4 ml of 1,1,2-trichloro-1,2,2-trifluoroethane (Freon 113) by tube inversion 5–10 times, sonicated for 15 seconds (0.5s on, 0.5s off) at 11% amplitude on a Branson Digital Sonifier in the cold room and centrifuged for 10 min in a Beckman Coulter rotor SX4750 at 200 g at 4 °C. The upper aqueous phase was removed and transferred to a new clean 15 ml conical tube, and the Freon extraction was repeated two more times. The resulting virus solution was divided in aliquots of 500 μl and layered on top of a 35–60.0% (w/v) CsCl gradient (1.26–1.45 g/ml) prepared in six SW60 tubes. CsCl solutions were prepared in TNC. Gradients were spun at 4 °C in a Beckman Coulter rotor SW 60 at 406,000 g for 2.5 h. Bands for TLPs and DLPs were harvested by side puncture. TLPs and DLPs were dialyzed against 2 L of TNC or TNE (20 mM Tris, 100 mM NaCl, 1 mM EDTA, pH 8.0) overnight. Virus particles were concentrated by pelleting at 4 °C in a Beckman Coulter rotor SW 60 at 406,000 g for 1 h. Supernatants were removed carefully, and pellets resuspended in 100–200 μl of remaining buffer. TLP concentration was measured by densitometry of VP6 bands on Coomassie stained SDS-PAGE gels against DLP standards ranging from 0.1 to 1.0 mg/ml.

*Purification of recombinant VP7* – Baculovirus vectors carrying the genetic information of full-length rhesus rotavirus VP7 (G3 serotype) were produced from bacmid transfected Sf9 cells (Thermo Fisher Scientific). The viruses were then inoculated into fresh Sf9 cells grown on T25 flasks and passaged three times with an incubation time of 72 h for each passage. Eight glass spinner flasks with 500 ml of Sf9 cells with approximately 2 million cells per ml were then infected with 13 ml of passaged virus stock solution and incubated for 72 h. Cells were harvested at 4 °C by centrifugation in Beckman Coulter rotor

J4.2 at 3,500 g for 30 min. Purification was performed as previously described (*3*), with minor modifications. The supernatant was collected and benzamidine and sodium azide were added to final concentrations of 1 mM and 0.01% (w/v), respectively. Supernatant was loaded into 50 ml of a concanavalin A Sepharose (ConA) resin. Resin was washed with five column volumes (CV) of TNC buffer. Protein was eluted with five CVs of TNC buffer containing 0.6 M α-methyl mannose. The eluate was then loaded onto 11 ml protein A resin with immobilized antibody m159 (*32*) (10 mg per ml of protein A resin), specific for trimeric VP7, and equilibrated in 20 mM Tris, 50 mM NaCl, 0.1 mM $CaCl_2$, pH 8.0. We washed bound VP7 protein with five CVs of buffer and eluted with five CVs of 20 mM Tris, 50 mM NaCl, 1 mM EDTA, pH 8.0. Buffer was exchanged by overnight dialysis in 0.1HNC (2 mM HEPES, 10 mM NaCl, 0.1 mM $CaCl_2$). Protein was frozen in liquid nitrogen and stored at -80 °C.

**Cryo-ET data collection and processing**

*Grid preparation* – BSC-1 cells were cultured overnight on 200-gold mesh Quantifoil grids (2/2) with $SiO_2$ previously incubated for 4 h with 1 mg/ml fibronectin (Corning). RRV, strain G3P5B[3], at 0.034 mg/ml was added and incubated for 10 min. The grids were then washed with medium and incubated for an additional 10–20 min. Samples were vitrified in liquid ethane using a Leica EM GP plunge freezer. Cell distribution was assessed by scanning electron microscopy (SEM) on a Thermo Fisher Scientific Aquilos 2 (fig. S1).

*Tomographic tilt series collection* – Tilt series, imaged at the thin cell periphery, were acquired on a Thermo Fisher Scientific Titan Krios G3i equipped with a Falcon 4i detector and a Selectris energy filter (10 eV slit width). Data were collected following a dose-symmetric bidirectional tilt scheme (*33*), with a nominal defocus range of -3 to -5 µm. Tilt angles spanned from +60° to -60° in 3° increments, yielding 41 tilts per series. Each tilt was acquired at a dose of 3.7–3.9 electrons per $Å^2$, for a total cumulative dose of ~150 electrons per $Å^2$. Four frames were recorded per tilt image. A total of 838 tilt series were acquired across seven datasets (table S1).

*Initial tomographic tilt series processing* – Tilt image movie frames were aligned and averaged with WarpTools version warp_2.0.0dev29 (fs_motion, grid 1×1×5) and used for contrast transfer function (CTF) parameter estimation (fs_ctf, grid 2×2×1) (*34*). To exclude bad tilt images from downstream processing, we calculated the pixel value distributions from the tilt image averages using IMOD (clip histogram -F 0.5,1) (*35*) and excluded bad tilt images from the corresponding mdoc files based on pathologic falloff of the density histograms. We made tilt series image stacks with WarpTools (ts_stack) and then used AreTomo2 version 1.1.2 (*36*) for initial tomographic alignment (VolZ 4400, AlignZ 1400). Alignments were imported with WarpTools (ts_import_alignments), and the handedness of the CTF model was checked (ts_defocus_hand) before fitting it for the entire tilt series (ts_ctf). We then reconstructed 4×- (~9.4 Å per pixel) and 8×-binned (~18.8 Å per pixel) tomograms with WarpTools (ts_reconstruct). The data processing workflow is summarized in fig. S2.

*Rotavirus particle picking (3D template matching)* – With e2spt_tempmatch.py from EMAN2 (*37*), we located rotavirus particles in the 4×-binned tomograms (~9.40 Å per pixel), using a previously determined icosahedrally averaged cryo-EM reconstruction of the rotavirus TLP (EMD-45118) scaled to the correct pixel size. We imposed $I_2$ symmetry in the computation and kept particles with a score above a defined threshold (vthr 8.0). We extracted 4×-binned 3D subtomograms with WarpTools (ts_export_particles, box 128) and calculated low pass filtered 2D projections with RELION (relion_project) (*38*) for visualization and manual exclusion of false picks. See table S1 for the number of particles from each dataset.

*Initial subtomogram alignment* – After extracting 2×-binned (~4.70 Å per pixel) 3D subtomograms and corresponding 3D CTF volumes with WarpTools (ts_export_particles, box 256), we used RELION (relion_refine) for reference-based subtomogram alignment. This step was redundant, as alignment parameters were in principle already obtained from the EMAN2 particle picking, but we included the step because e2spt_tempmatch.py did not output the alignment angles. The alignment-angle convention is the same in Warp/M and RELION. Alignment and reconstruction were done with $I_2$ symmetry imposed, and the reference was masked with a spherical mask, outer radius 512 Å.

*Tomographic tilt series refinement* – We used M (MCore, version warp_2.0.0dev29) for tomographic tilt series refinement (*39*). We created a population with a single species (as defined by Warp/M) based on the previously located and aligned rotavirus particles. The box was 512 cubic pixels (unbinned data) at ~2.35 Å per pixel (table S1). As mask, we used a low-resolution envelope of the three protein layers (VP2, VP6, VP7) and the VP4 spikes in upright conformation. Several iterations of M refinement were carried out with $I_2$ symmetry imposed and with a sequential increase in the complexity of the distortion model (image warp, particle poses, stage angles, volume warp, defocus, tilt movie refinement, magnification distortion, CTF refinement). We did the final refinement rounds after fitting weights per tilt series and fine-sampling the trajectories of the particle poses to 3. We determined the true pixel size for each dataset (table S1) by scaling the final reconstruction to a reference, a previously determined cryo-EM reconstruction of the virus at 2.36 Å resolution (EMD-45118) (fig. S3A). The Fourier shell correlation (FSC) curves calculated from the half maps for the icosahedral reconstructions of the 7 datasets are shown in fig. S3B. We also compared the reconstructions to a "perfect" model, the 2.36 Å cryo-EM reconstruction (EMD-45118), to assess the extent of fitted noise after tilt series refinement (fig. S3C). The variation in the observed resolution between reconstructions from different datasets (ranging from to 4.7 [Nyquist] to 8.8 Å, table S1) arose primarily from differences in the average thickness of the sample and stage stability during data collection. The appearance of the virus density is consistent with the reported resolution (fig. S3, D and E).

*Subtomograms of spike positions* – For the analysis of individual spike positions, we symmetry-expanded the virus particles from $I_2$ to $C_1$ with M (MTools, expand_symmetry). We then shifted the particle poses and corresponding reference maps and masks along x, y, and z by 132, 298, and 194 Å, respectively, thus placing the VP4 spike (in upright or reversed conformation) at the center of the new box. We then repeated the M tilt series refinement with the symmetry-expanded particles at the shifted positions (box of 512 cubic pixels). This step allows local spike reconstructions and spike subtomograms to be identified and retrieved with specified grid samplings and box sizes.

*Tomograms* – Tomograms for visualization and particle picking (3D template matching) were calculated with Warp (WarpTools, ts_reconstruct) before and after tomographic tilt series refinement at 10.0 and 20.0 Å per pixel.

*Merging of datasets* – Table S1 lists pixel sizes that we used for processing the individual datasets. A posteriori, we determined the accurate pixel size for each dataset by comparing their icosahedral full virus reconstructions with the 2.36 Å cryo-EM reconstruction (EMD-45118), for which the scale had been validated by a refined atomic model with full stereochemical restraints. The errors in the processing pixel sizes were small (table S1), and CTF correction or merging of subtomograms was not substantially affected. Because of the 100 nm-diameter rotavirus particle size, however, the errors led to small offsets of the spike subtomogram box centers among the datasets. We therefore corrected these offsets using Warp (WarpTools, shift_species) to adjust the relative box center offsets after determining shifts based on rigid body-fitted models into the spike consensus reconstruction of the individual datasets. We chose a consensus pixel size of 2.35 Å for processing of the merged subparticles from all datasets.

*Metadata handling and unique particle identifiers* – At each computational step, from particle picking to classification and selection of spike subparticles, we carried forward a star file with particle metadata including unique particle identifiers. After certain processing steps (e.g. after subtomogram extraction, refinement, or particle shifts), the particle poses changed, and the particle order in the default metadata files was not preserved (depending on the number of parallel jobs and order of submission during Warp/M computation). We used Python routines from Dynamo (*40*) to read and write star files. Python dictionaries, to lookup particle hashes, and the computation of quaternion distances (calculated from the alignment angles) helped with the non-trivial task of efficiently matching particles between Warp/M input and output files. Thus, every spike subtomogram had a unique particle identifier (dataset, tomogram, virus, spike subparticle), which allowed us to track it from the reconstructed tomogram to a final M reconstruction and 3D classification.

*Spike subtomogram classification* – For local classification of spike positions, we extracted 3D particles with corresponding 3D CTF volumes (132 cubic pixels, 3.2 Å per pixel) and applied a similar strategy as previously used for single particle data (*4, 13, 24*). Prior to running the 3D classification in RELION, we masked each individual spike subtomogram, using a mask encompassing the volume of a single VP6 and VP7 trimer, plus the volume occupied by VP5*/VP8* in upright or reversed conformation. Before applying the 3D mask to the subtomogram, the mask was transformed according to the particle alignment angles and shifts to match the subtomogram as it was extracted. The same 3D mask was then also used to mask the reference volumes during classification. Classification was carried out without alignment. We essentially obtained the same result when requesting 4, 8, or 12 classes. A first round of classification was done for each dataset (classification 1) and the result is summarized in fig. S4. All subparticles belonging to either upright, reversed, or empty classes were merged and re-classified (classification 2), after which we obtained the following assignment for all spike positions: upright, 299,405 particles (59%); reversed, 55,972 particles (11%), empty, 158,523 particles (31%) (fig. S5).

*Supervised classification of VP7* – We made DLP and TLP masks, respectively, from the consensus RRV model (PDB-ID pdb_00009c1g, symmetry expanded to generate the full virus). For this, we selected all VP2A/B, VP6, and VP7 (in case of the TLP mask only) chains within the spike position subparticle box and made masks using a 4 Å probe radius with CCP4 (*41*), which were then multiplied with a 192 Å-radius spherical mask. We then masked the M reconstruction from all subparticles to generate references for DLP and TLP, respectively. Supervised classification was done in RELION with the DLP/TLP references and the TLP mask as solvent mask, for one iteration and without alignment (--iter 1 --tau2_fudge 0.2 --K 2 --ini_high 10 --skip_align). The degree of VP7 uncoating shown in Fig. 2 and table S2 is per asymmetric unit of the icosahedral virus, where each of the 60 asymmetric units (containing 13 VP7 mononers each) was assigned either as VP7-occupied (TLP) or uncoated (DLP) based on the supervised classification result observed for the corresponding spike position.

*Membrane segmentation* – We segmented membrane in all tomograms (reconstructed at 10 Å per pixel) with MemBrain (*42*) using a pre-trained MemBrain segmentation model (MemBrain_seg_v10_beta) (fig. S7, A and B). For further analysis (see membrane overlap below) and display, the volumes containing the segmentation result were Fourier-sampled (for display) or mean-binned (for membrane overlap calculation [see below]) to a pixel size of 20 Å, masked (removing membrane inside viruses, false segmentation) and dedusted (surface level 0.2, size 600) using ChimeraX (*43*).

*Membrane overlap and distance measurements* – We calculated a membrane overlap value for each spike position from the M-aligned rotavirus particles and the membrane-segmented tomograms (see above, mean-binned to 20 Å pixel size, masked and dedusted), by counting the sum of density of segmented membrane within a 160 Å-radius mask centered at each spike position (fig. S7, A to D). We applied the 160 Å-radius mask first to the membrane-segmented tomogram, converting the map to a NumPy array (*44*), and then taking the sum of it. We also calculated a membrane distance value for each spike position, with both, the upright and reversed conformations, as reference points (fig. S8). To do so, we determined the intersection of segmented membrane with the local 2-fold or 3-fold axes, defined by the VP5* upright and reversed structures, respectively (placed at each spike position regardless of its configuration). At each spike position, we cropped a box from the membrane segmentation tomograms, aligned the virtual 3-fold along the z axis, applied a cylindrical mask (5-pixel radius), and projected the density on the z axis. We detected peaks in the 1D membrane density profile using SciPy (*45*) (peaks with a peak height of >50% of the maximum were kept), and fitted each peak with a Gaussian. Histograms of membrane distance values for the upright and reversed spike conformation are shown in fig. S8. For an accurate location of the membrane in case of the reversed VP5* trimer shown in Fig. 3C, we calculated the average rotational density average (RDA) from the central section of 42 viruses with extensive membrane overlap and relatively uniform close membrane contacts (Fig. 3, B and C).

*Classification of membrane-engaged, reversed-conformation VP5** – From the reversed 55,972 spike subparticles, we selected 5,539 that had been assigned as TLP in the supervised VP7 classification (not uncoated), had a membrane overlap of more than 50, and a latitude of less than |±45°|. To allow classification focusing on the reversed VP5* trimer with $C_3$ symmetry imposed, we changed the particle alignment in the star file such that all VP5* 3-fold axes aligned and passed through the center of the box when calculating a reconstruction. We 3D classified this re-oriented particle stack in RELION (classification 3, fig. S9).

*Ribosome supervised classification* – We picked ribosomes in the 4×-binned tomograms (10.00 Å per pixel) with easymode in standard ribosome-segmentation mode (*46*). This yielded a total of 36,256 ribosome coordinates from all tomograms. Initial 2×-binned subtomograms were extracted with WarpTools and aligned with RELION. Refinement in M, first with a 80S mask, then with a 60S mask, converged to a consensus reconstruction with overall Nyquist resolution of 4.7 Å (fig. S10, A to C). To determine the state of each ribosome, we used a supervised classification approach, based on references prepared from the PDB files of a previous publication, where the authors obtained high-resolution structures of 23 ribosomal states by cryo-EM from human cells after focused ion beam (FIB) milling thin lamella (*20*). We first rigid-body fitted the 60S submit of the state 1 structure (PDB-ID pdb_00009p72) into the consensus reconstruction with phenix.real_space_refine (*47*) and then superimposed all structures onto the 60S RNA (PDB-IDs pdb_00009p72, pdb_00009p6z, pdb_00009pkg, pdb_00009p8b, pdb_00009p7o, pdb_00009p76, pdb_00009p7k, pdb_00009p7n, pdb_00009p7l, pdb_00009p7y, pdb_00009p7x, pdb_00009p7w, pdb_00009p8h, pdb_00009p73, pdb_00009p9i, pdb_00009p9h, pdb_00009p78, pdb_00009p79, pdb_00009p8c, pdb_00009p7a, pdb_00009p7c, pdb_00009p7d, pdb_00009p7e). We calculated structure factors with phenix.fmodel and maps with fft from CCP4 (*41*). The radial structure factor for all 23 references was matched with EMAN2 (*48*). Supervised classification was done in RELION with the 23 ribosomal maps as references and a 80S mask, for one iteration and without alignment (--iter 1 --tau2_fudge 0.2 --K 23 --ini_high 8 --skip_align). The resulting class distribution is shown in fig. S10, D and E, and movie S3. 86% of the picked particles classified as translating ribosomes (e.g. belonging to an elongation cycle state) and 14% classified as non-translating ribosomes (e.g. belonging to a hibernating state). We calculated ribosome density maps (fig. S10F) by summing at each voxel of a tomogram (sampled at a pixel size of 50 Å) the density in the corresponding easymode ribosome segmentation map within a sphere of 1500 Å radius.

*Tomogram interpretation display* – For the figures shown in Figs. 1D and 2 (middle column), we populated individual maps for each component (e.g. VP2A, VP2B, VP6, VP7, VP4, ribosomes) with the corresponding reference structure at the location orientations as determined from the subtomogram analysis and classification.

**Liposome disruption experiment, TIRF and cryo-EM**

*Liposome preparation* – A 2 mg total lipid mixture in chloroform of cholesterol, phosphocholine egg extract (egg PC), sphingomyelin (SM), 1,2-dioleyl-sn-glycero-3-phopshoethenolamine (DOPE), biotinylated-DOPE, Cyanine5-DOPE (Cy5-DOPE), 1-palmitoyl-2-oleyo-sn-glycerol-3-phosphoetanolamine (POPE) and 1,2-dioleyl-sn-glycero-3-phospho-L-serine (DOPS) in molar ratio (40:22.5:10:7.25:0.5:0.25:8.5:8) was dried in a round bottom glass tube under argon gas flow. Remaining solvent was removed by incubating the lipids under high vacuum overnight. Dried lipids were resuspended in 250 μl of EDTA buffer (20 mM Tris pH 8.0, 100 mM NaCl, 1 mM EDTA) containing 2 mM carboxyfluorescein (CF) 5 and 6 dye by vortexing for two minutes, forming an 8 mg/ml lipid suspension, which was subjected to two freeze-thaw cycles with liquid nitrogen. Liposomes were then formed by extrusion with 41 passages through a 200 nm pore filter using a mini extruder (Avanti Research). The liposome suspension was loaded onto a G25 PD10 column equilibrated with TNE to separate liposomes from non-incorporated fluorophore. Fractions of 500 μl were collected.

*TIRF measurements* – 25 mm circular glass coverslips were cleaned and coated with a 10% (w/v) solution of biotinylated-polyethylene glycol (bio-PEG) and PEG in a ratio 1:99 (Laysan Bio, cat. no. mPEG-SCM-5000) (*49*) and pre-incubated with 0.5 mg/ml neutrAvidin (Thermo Scientific, cat. no. LF144746) for 15 min at room temperature (RT) before use. Liposome solutions were diluted 100 times in EDTA buffer or $Ca^{2+}$ buffer (20 mM Tris pH 8.0, 100 mM NaCl, 1 mM EDTA) and incubated over the coverslip for 10 min at RT. Unbound liposomes were removed before imaging by several washes with EDTA or $Ca^{2+}$ buffer. Upon addition of either EDTA buffer alone, VP7 trimer (calcium buffer) at 1.5 mg/ml, or VP7 monomer (EDTA buffer) at 1.5 mg/ml. Defined Trypsin Inhibitor (Gibco) at 1 μg/ml was added to prevent proteolysis. Movies were started on a Axio Observer Zeiss microscope equipped with Vector3 illumination hardware and a Photometrics Prime 95B sCMOS Camera (Teledyne, USA). Images were acquired using a Zeiss 100× 1.46 NA oil-immersion objective (Jena, Germany) with excitation from 488 and 640 nm solid-state lasers (LaserStack v4, Intelligent Imaging Innovations [3i]), both operated at 100 mW. Ring-TIRF acquisition was used through Vector3, in which the excitation beam was scanned circularly at the back focal plane of the objective during each camera exposure, producing an azimuthally averaged evanescent field at the glass–sample interface. The incident angle was set above the critical angle for total internal reflection and held constant throughout each experiment. For each field of view, we recorded 30-min movies at 1 min per frame at 50% power from each laser and 10 ms exposure at 488 nm (CF signal) and 50 ms exposure at 640 nm (Cy5-DOPE dye, liposome signal). The microscope was controlled by the SlideBook acquisition program (3i). Raw images were recorded without post-acquisition illumination correction.

*Image analysis* – We used previously described custom-made MATLAB (MathWorks) scripts (*50*) for automatic detection of fluorescent liposomes in the 640 nm channel (Cy5-DOPE dye, liposome signal) by fitting a 2D Gaussian function to diffraction-limited spots with intensities >1.5-fold above background. We extracted the XY position of each liposome at all time frames in the movie and calculated the integrated fluorescence intensity for the CF and lipid channels within a 4×4 pixel area around the XY position. We calculated background as the integrated fluorescence intensity in a 6×6 pixel frame around the 4×4 region containing the signal, normalized the background value by area, and subtracted it from the 4×4 pixel integrated intensity. Only liposomes with detectable lipid signal at all time points were considered for analysis. Signal loss was quantified by comparing the average fluorescence of the first two time points (min 1 and 2) to the average of the final two time points (min 29 and 30) in the movie and calculating the percentage of signal loss between these two values.

*Cryo-EM of liposomes incubated with VP7* – 3 µl of liposomes were incubated with 15 µl of VP7 (40 µM) or VP7$^{292\text{-}312}$ peptide (100 µM, synthetized by TUCF Peptide Synthesis) at 37 °C for 30 min. 4 µl of the sample was then placed on a glow-discharged 1.2/1.3 Quantifoil grid, blotted for 6 s at 90% humidity and plunge frozen in liquid ethane in a Vitrobot (Thermo-Fischer). Grids were stored in liquid nitrogen until imaging. Micrographs of liposomes were recorded on a Thermo Fisher Scientific Talos Arctica equipped with a Gatan K3 detector at a pixel size of 1.44 Å$^2$ with a total dose of 50 electrons/Å$^2$ at a defocus of -2.0 to -2.5 µm. SerialEM version 4.2.12 was used for data acquisition (*51*).

*Tomographic tilt series collection* – Tilt series of liposomes incubated with VP7 were acquired on a Thermo Fisher Scientific Talos Arctica equipped with a Gatan K3 detector at a pixel size of 1.1 Å. Data were collected following a dose-symmetric bidirectional tilt scheme (*33*), with a nominal defocus range of -1 to -3 µm. Tilt angles spanned from +60° to -60° in 3° increments, yielding 41 tilts per series. Each tilt was acquired at a dose of 4.17 electrons per Å$^2$, for a total cumulative dose of ~170 electrons per Å$^2$. Four frames were recorded per tilt image.

*Liposome tomographic tilt series processing* – Tilt image movie frames were aligned and averaged with WarpTools version warp_2.0.0dev36 (fs_motion, grid 1×1×5) and used for contrast transfer function (CTF) parameter estimation (fs_ctf, grid 2×2×1) (*34*). To exclude bad tilt images from downstream processing, we calculated the pixel value distributions from the tilt image averages using IMOD (clip histogram -F 0.5,1) (*35*) and excluded bad tilt images from the corresponding mdoc files based on pathologic falloff of the density histograms. We made tilt series image stacks with WarpTools (ts_stack) and then used AreTomo2 version 1.1.2 (*36*) for initial tomographic alignment (VolZ 4400, AlignZ 1400). Alignments were imported with WarpTools (ts_import_alignments), and the handedness of the CTF model was checked (ts_defocus_hand) before fitting it for the entire tilt series (ts_ctf). We then reconstructed 10×-binned (~10.0 Å per pixel) tomograms with WarpTools (ts_reconstruct).

*Membrane segmentation of liposome data* – We segmented membrane in tomograms (reconstructed at 10 Å per pixel) with MemBrain (*42*) using a pre-trained MemBrain segmentation model (MemBrain_seg_v10_beta) (movie S4).

**Infectivity assays**

*Infectivity assay of TLPs* – Focus-forming unit (FFU) concentrations for TLP were determined by infectious focus assays as previously described (*52*). Specific infectivity was calculated by normalizing FFU titers to particle concentrations determined by densitometry of Coomassie blue-stained SDS-PAGE gels. For the experiments with Latrunculin A (Lat A), cells were incubated for 1 hour with 5 µM Lat A, which was diluted in Dulbecco's Modified Eagle's Medium (DMEM, Thermo Fisher Scientific) from a 1 mM stock solution in dimethyl sulfoxide (DMSO). BSC-1 cells, grown overnight in a 96-well plate, were washed with DMEM and inoculated with serial dilutions of trypsin activated virus in DMEM containing 5 µM Lat A and 1 µg/ml trypsin. For the assays testing trypsin inhibitor, cells were washed and inoculated with serial dilutions of trypsin activated virus in presence of either 1 µg/ml trypsin or 1 µg/ml Defined Trypsin Inhibitor (Gibco). After 1 h at 37 °C incubation with virus particles, the inoculum was removed and replaced with DMEM containing 10% fetal bovine serum and 2.5 μg/ml neutralizing monoclonal antibody M159 to prevent secondary infection. At 16 h post-infection, cells were washed with phosphate-buffered saline (PBS) and fixed with methanol. Infectious FFUs were detected by immunoperoxidase staining using monoclonal antibody M60 (anti-VP7) as the primary detection antibody, and peroxidase labeled Goat anti-Mouse IgG (SeraCare) as secondary antibody. Once developed, FFU were counted by light microscopy.

**GCaMP live-cell fluorescence microscopy experiments**

*Cell culture and transfection* – BSC-1 cells were plated on glass-bottom 18-well dishes at 70% confluence and allowed to adhere overnight in DMEM Glutamax, supplemented with 10% Hi-FBS (Thermo Fisher Scientific). The following day, cells were transiently transfected with plasmid DNA (Addgene plasmid # 52228 – pGP-CMV-GCaMP6s-CAAX, a gift from Tobias Meyer) using Lipofectamine 2000 (Thermo Fisher Scientific). Briefly, plasmid DNA and Lipofectamine 2000 were diluted separately in antibiotic-free OPTI-MEM, combined, and incubated for 20 min to allow complex formation before addition to cells. Cells were incubated with transfection complexes for 6 h, after which the medium was replaced with fresh DMEM Glutamax, supplemented with 10% FBS. Cells were then allowed to recover and express the transgene overnight.

*Labeling of TLPs* – TLPs were diluted to a final concentration of 1.0 mg/ml in a total volume of 60 μl using HNC buffer (20 mM HEPES, pH 8.0, 100 mM NaCl, 1 mM $CaCl_2$). Subsequently, 6.7 μl of 1 M $NaHCO_3$ (pH 8.3) was added, followed by 1.9 μl of 0.0076 mg/ml Atto 565 NHS ester and incubated for 1 h at room temperature. The reaction was quenched by addition of 5 μl of 1 M Tris (pH 8.0). Labeled particles were buffer exchanged into 20 mM Tris (pH 8.0), 100 mM NaCl, and 1 mM $CaCl_2$ using Zeba Spin Desalting Columns (Thermo Fisher Scientific).

*Confocal microscopy and live-cell imaging* – Labeled TLPs were added to cells at a final concentration of 0.030 mg/ml and incubated during 5 min before starting imaging. Movies of 10 min with a time-lapse interval of 10 s were acquired using a Zeiss Axio Observer Z1 inverted microscope equipped with environmental control (37 °C, 5% $CO_2$) and a CSU-X1 spinning disk confocal unit (Yokogawa), operated by the Marianas (3i) platform. The system included a spherical aberration correction module and a 3i LaserStack with diode lasers at 488, 561 (150 mW), and 640 nm (100 mW). Emission was filtered using 525/40, 609/54, and 692/40 filters (Semrock). Single plane movies were acquired with 50 ms exposure using a dual-camera sCMOS system (Prime 95B, Teledyne Photometrics) and a 40× objective. Final XY resolution was 0.22×0.22 μm/pixel. Background intensity was subtracted using Fiji (*53*) and fluorescence intensity of the GCaMP signal in in the cell was plotted as a function of time.

**Figure preparation**

Figures were prepared with PyMOL (The PyMOL Molecular Graphics System, Version 2.3 Schrödinger, LLC), ChimeraX (*43*), Fiji (*53*), and matplotlib (*54*).

**Supplementary Text**

*Subtomogram averaged map of VP5* in reversed-conformation spike*s *(outline of procedure)* – Full details are in the cited Supplementary Figures and Methods. We found particle positions in the tomograms by 3D template matching (*55*), refined their positions and orientations with RELION (*56*) and M (*39*) (see Methods, figs. S2 to S3, and table S1), and extracted 3D subparticles corresponding to VP4 spike positions (figs. S4 to S6). To identify reversed, membrane-bound spikes, we defined an overlap factor between each spike location and the surrounding membrane, which we segmented using MemBrain (*42, 57*) (see Methods and fig. S7, A to D). Viruses with high overall membrane overlap showed extensive membrane wrapping (fig. S7, E and F). Moreover, spikes that classified as reversed showed on average substantially higher membrane overlap than did upright ones (fig. S7, G to J, and data S1), confirming the correlation between the VP5* conformational change from upright to reversed and the close contact with membrane. We also calculated the distance of segmented membrane from spike reference points (see Materials and Methods). The distribution maximum for distances at spike positions that classified as upright was longer than that for positions that classified as reversed (fig. S8).

*Initiation and growth of VP7-generated pore* – Initiation of pore formation might start by association of the VP7 amphipathic helix with the membrane surface, with the hydrophobic side facing inward and the hydrophilic side facing outward, followed by transition to a small, transmembrane pore when the surface concentration has reached a critical level. Growth could be fed by recruitment of additional, surface-bound VP7. Unlike the large, β-barrel pores of cholesterol-dependent toxins such as perfringolysin O, there is no pre-pore ring (*58*). Proposals for formation of small pores by amphipathic helices include either a "barrel-stave" arrangement, in which the helices, approximately normal to the plane of the membrane, form a barrel that defines the circumference of the pore, with hydrophobic residues facing the fatty-acyl chains and hydrophilic residues facing the aqueous channel (*59, 60*), or a related alternative, a "toroidal pore", in which the helices are not in tight lateral contact, and lipid headgroups project between them, as at the edges of a bicelle (*61*). It is possible that the VP7 perforations initiate as barrel-stave or toroidal pores, with the rim turning outward as the pore grows, but our current structural data are not of sufficient resolution or contrast to show early events. Cryo-EM images of pores induced in mitochondrial outer membranes and liposomes by the pro-apoptotic protein Bax are very similar to those in Fig. 4D (*62, 63*), but most models for pore formation by Bax and related proteins involve association of globular domains along with insertion of an amphipathic peptide (*64, 65*). Similar apparent pore structures from intact VP7 monomers and amphipathic peptide alone rule out such interactions for the perforations seen here.

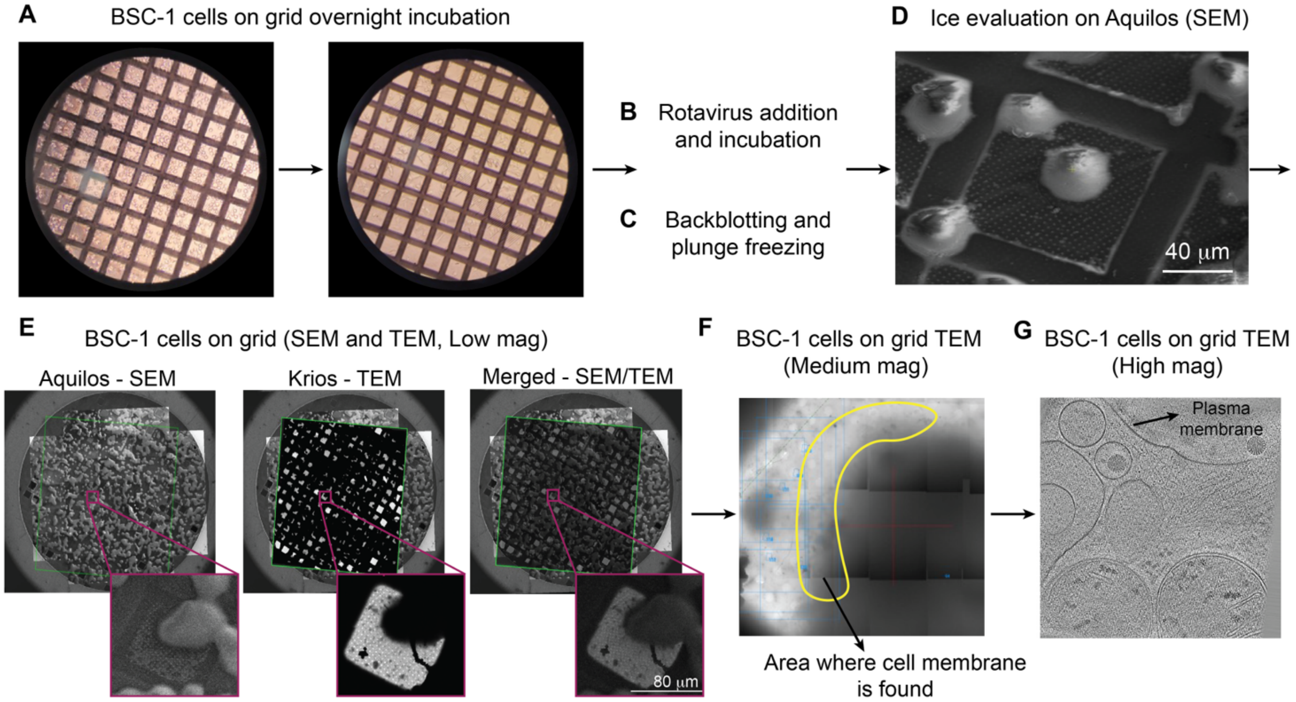


**Fig. S1. Cryo-ET sample preparation workflow for rotavirus infected BSC-1 cells.** (**A**) Sample preparation protocol. BSC-1 cells were seeded on gold EM grids and incubated overnight. (**B** and **C**) Cells were incubated with RRV for 10 min and with virus-free medium for another 10 min, then back-blotted and plunge-frozen in liquid ethane. (**D**) Representative scanning electron micrograph (SEM) of vitrified BSC-1 cells on a grid square, acquired on an Aquilos cryo-SEM. Ice quality was evaluated by assessing cell morphology and ice thickness. (**E**) Low-magnification images were acquired on an Aquilos (scanning electron microscopy: SEM, left) and Krios (transmission electron microscopy: TEM, center) to evaluate the quality and quantity of cells on the grid. Insets show higher magnification views of selected cells. (**F**) Medium-magnification TEM image montage of a BSC-1 cell on a grid square. The yellow outline shows the cell membrane boundary and suitable regions for tomogram acquisition. (**G**) High-magnification TEM image showing the quality of cellular preservation with clearly visible membrane bilayers and intracellular structures suitable for cryo-ET.

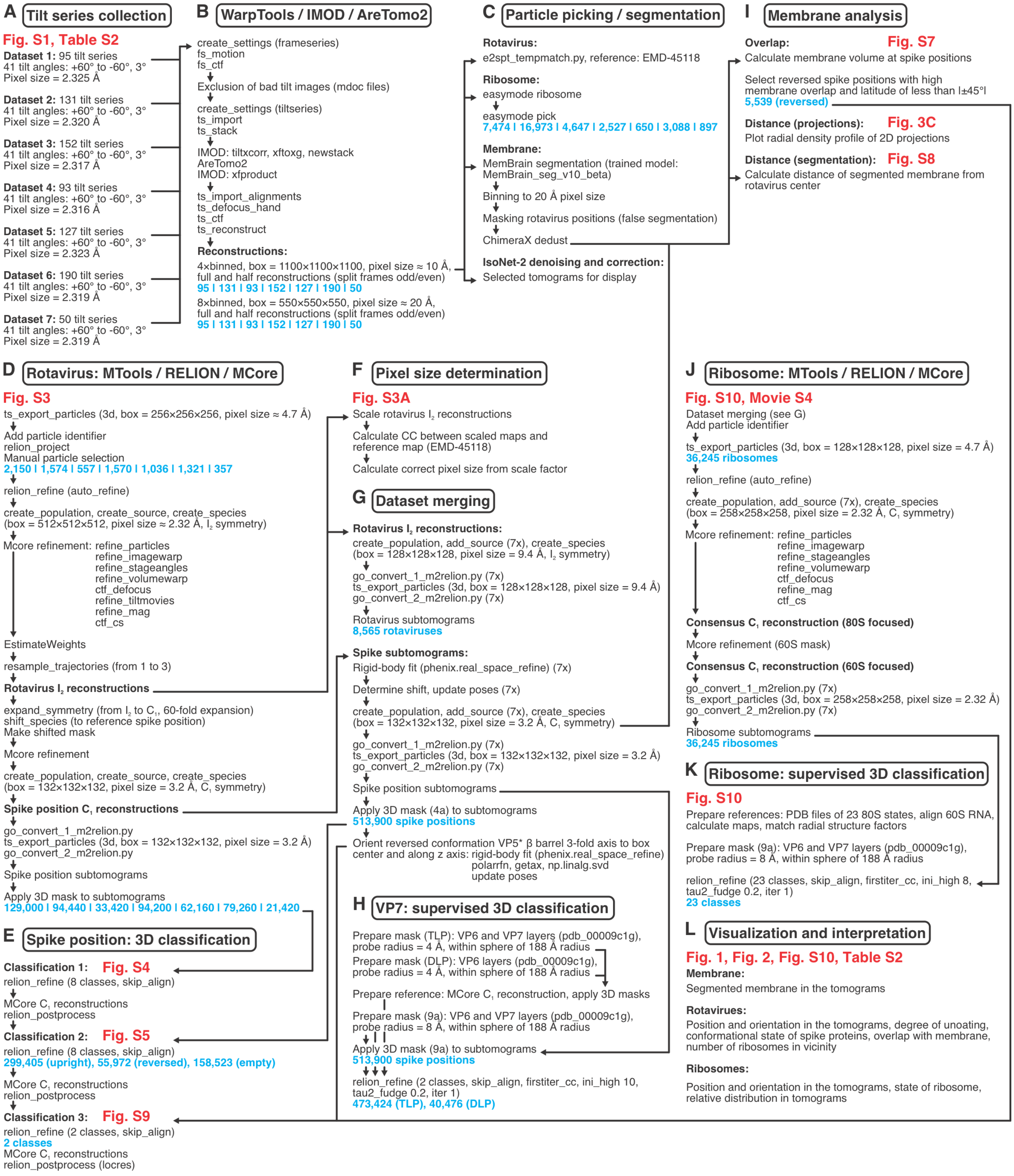


**Fig. S2. Summary of the cryo-ET data processing workflow.** (**A** to **L**) See Materials and Methods for additional details of the processing steps.

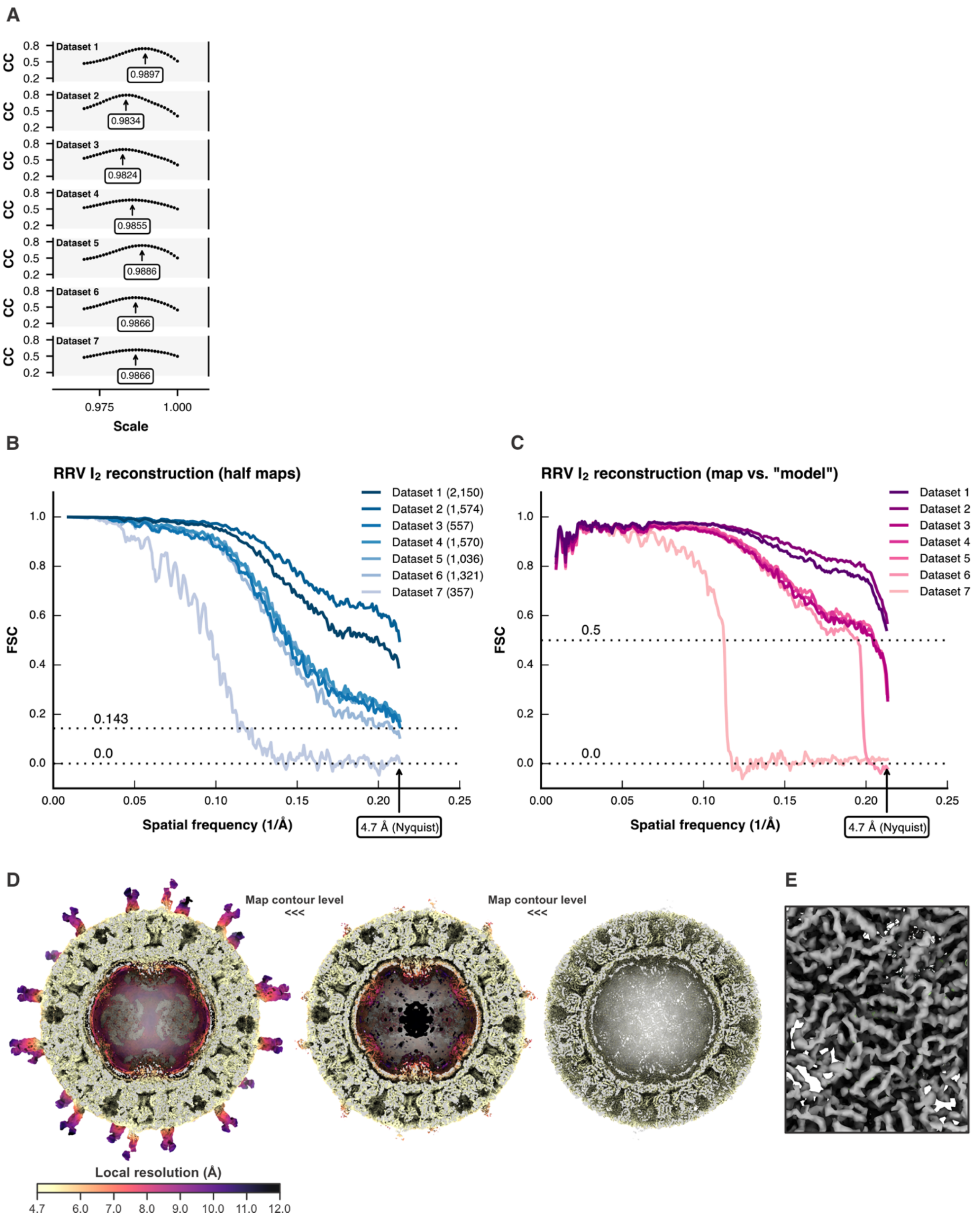


**Fig. S3. Icosahedral (I$_2$) rhesus rotavirus (RRV) reconstructions from subtomograms.** (A) Determination of the true pixel size for each dataset (table S1) using a 2.36 Å resolution cryo-EM RRV map (EMD-45118) as reference. Map correlation coefficients (CC) were calculated between differently scaled subtomogram reconstructions and the reference map. The scale factors corresponding to the true pixel size are highlighted with an arrow. (**B**) Fourier shell correlation (FSC) curves calculated from the final M reconstruction half maps for each of the 7 datasets. Maps were scaled to the true pixel sizes (table S1 and fig. S3A). The number of particles is given in parenthesis. The conventional cutoff of 0.143 for meaningful half map correlation is indicated. Nyquist frequency corresponds to 4.7 Å. (**C**) FSC curves calculated from the final M reconstruction map and a 2.36 Å high-resolution cryo-EM RRV map (EMD-45118), serving as "model". Maps were scaled to the true pixel sizes (table S1 and fig. S3A). The conventional cutoff of 0.5 for meaningful map-model correlation is indicated. Nyquist frequency corresponds to 4.7 Å. (**D**) Dataset 2 $I_2$ RRV reconstruction, filtered and colored according to local resolution, at increasing contour level from left to right. (**E**) Close-up view of the VP2A/VP2B inner layer density (dataset 2, M-sharpened map) showing molecular features (helical grooves and appearance of bulky side chains) consistent with a 4.7 Å-resolution map.

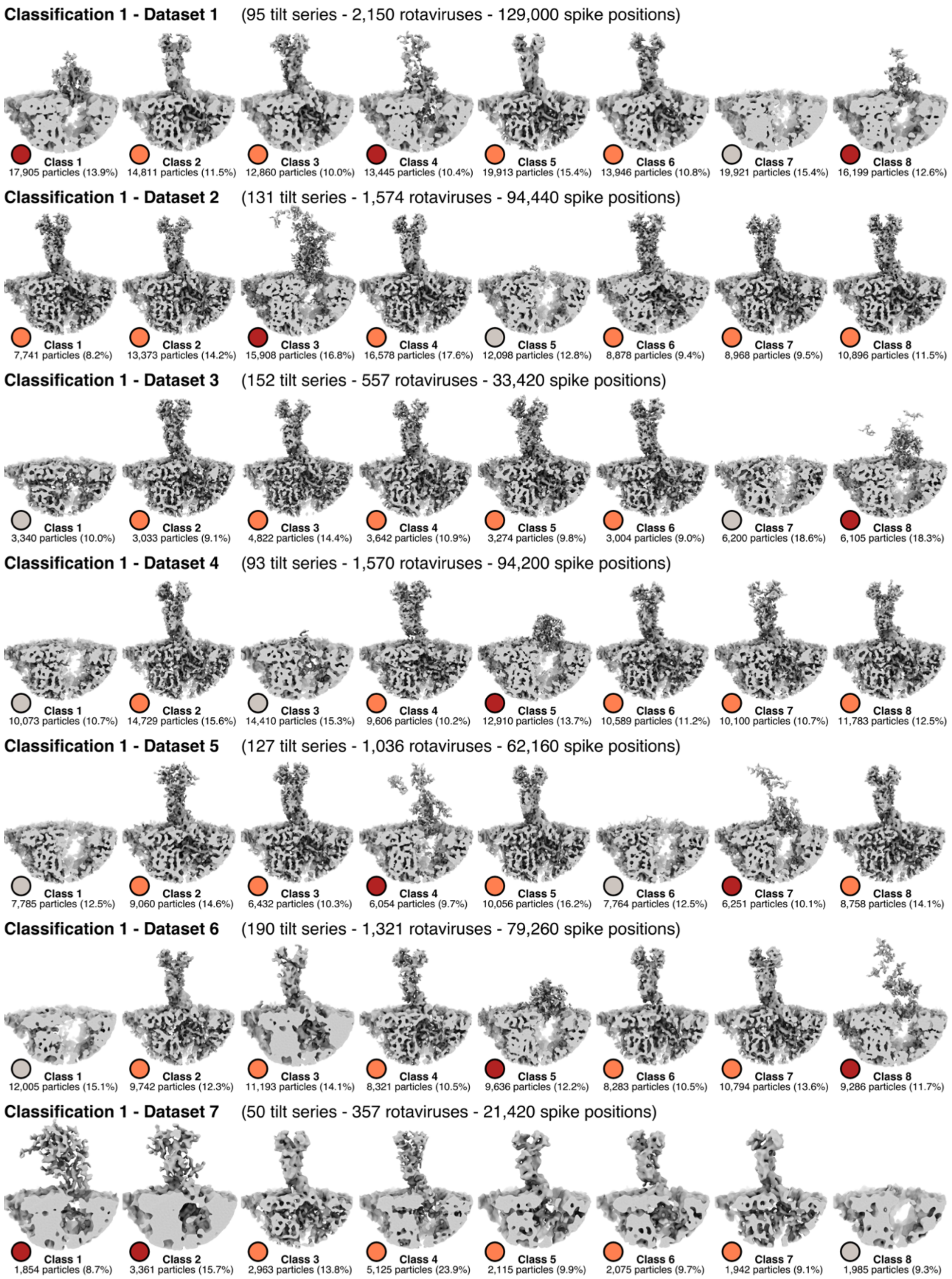


**Fig. S4. 3D classification 1 of spike positions.** For each of the 7 datasets, 3D classification 1 was done with RELION, and maps shown here were reconstructed with M (see Methods). For each class, the number of subparticles and the percentage of the total number of particles is given. Spike position configuration is color coded for each class: upright, light red; reversed, dark red; empty (unoccupied, no VP4 present), gray.

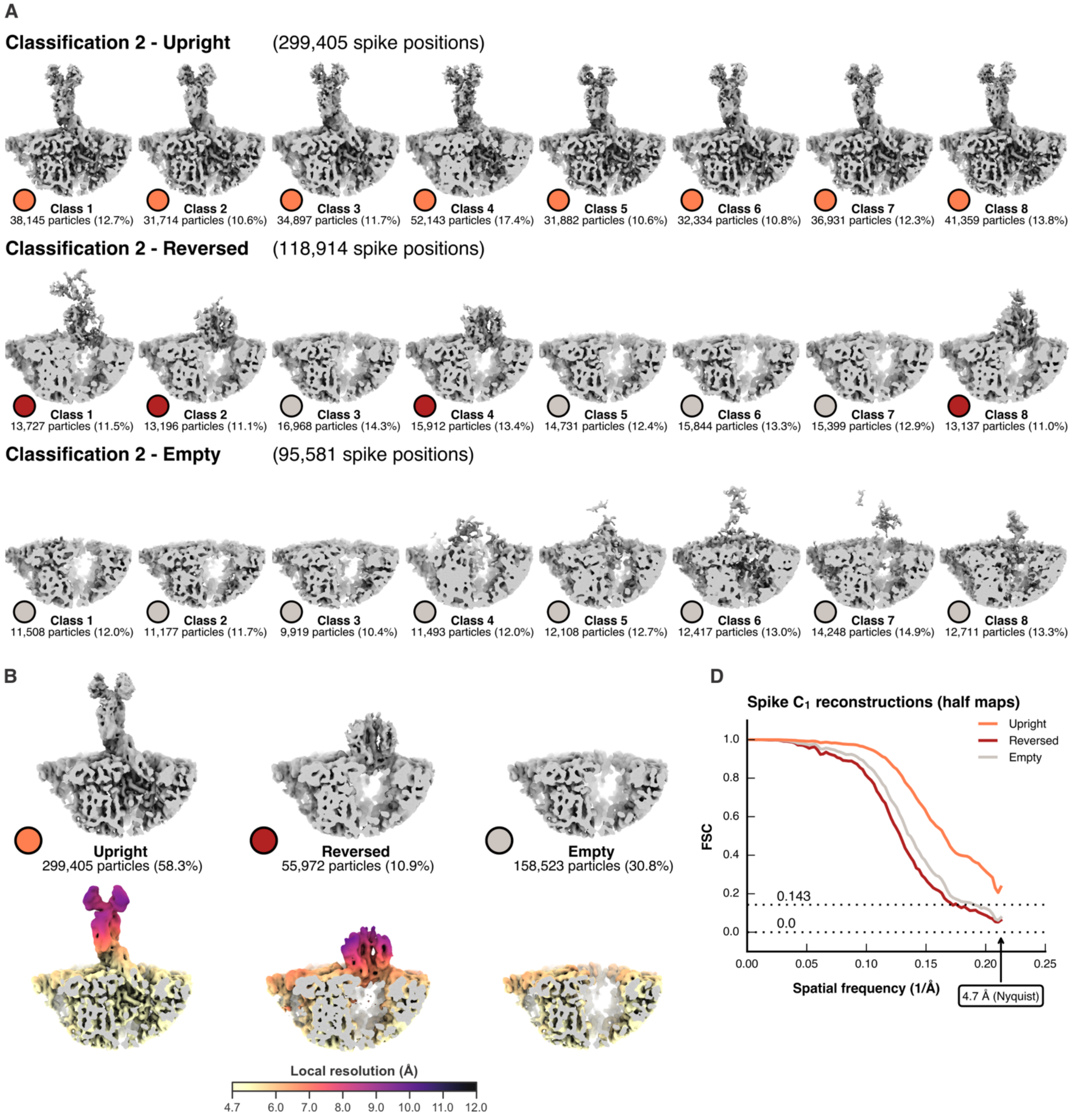


**Fig. S5. 3D classification 2 of spike positions.** (**A**) After merging upright, reversed, and empty classes (fig. S4) from each dataset, 3D classification 2 was done with RELION, and maps shown here were reconstructed with M (see Materials and Methods). Spike position configuration is color coded: upright, light red; reversed, dark red; empty (unoccupied, no VP4 present), gray. (**B**) 3D classification 2, the particles partitioned into 299k upright, 56k reversed, and 159k empty spike positions (top row). Local resolution mapped on the reconstructions (bottom row). (**C**) Fourier shell correlation (FSC) curves calculated from the half maps for each of the three states.

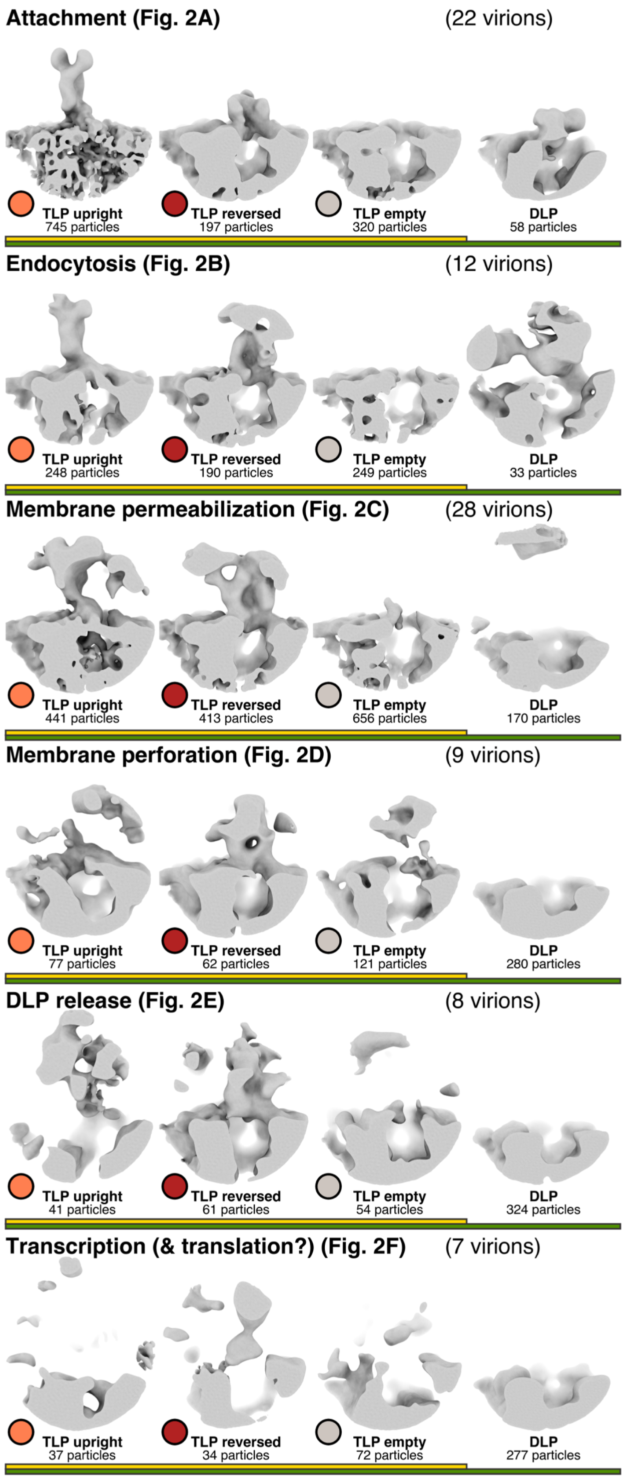


**Fig. S6. Subtomogram averages calculated for spike positions from set of virions assigned to distinct entry steps in table S2.** For each entry step shown in Fig. 2, A to F, and table S2, subtomogram averages were calculated (maps are local-resolution filtered) from the set of particles (spike position subtomograms) that classified either as TLP (yellow) upright (light red), TLP (yellow) reversed (dark red), TLP (yellow) empty (gray) or DLP (green).

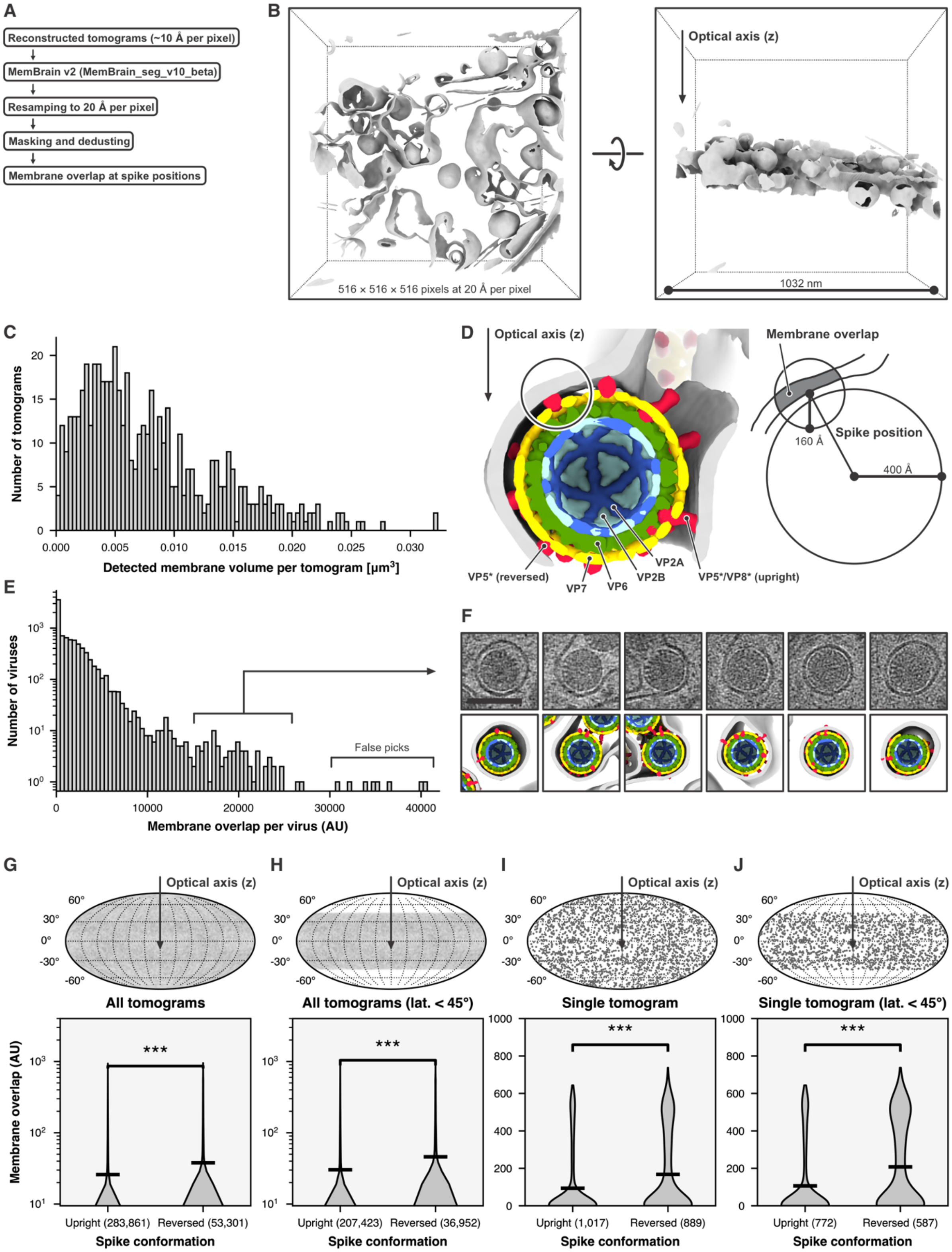


**Fig. S7. Overlap of spike positions with segmented membrane.** (**A**) Overview of the steps performed to calculate the overlap between spike location and segmented membrane. (**B**) Membrane segmentation result for a representative tomogram, viewed along the optical axis of the microscope (left) and from the side (right). (**C**) Histogram showing the distribution of total detected membrane volume per tomogram.

(**D**) Illustration how the overlap between spike location and segmented membrane was calculated for each spike position. The membrane overlap is defined as the volume of segmented membrane within a sphere centered at the spike position with a radius of 160 Å. (**E**) Histogram showing the distribution of total membrane overlap per virus in arbitrary units (AU). Particles with the highest values were false template matching picks. (**F**) Representative examples of rotaviruses with high membrane overlap (extend close membrane contacts), shown as 10-nm thick tomographic slices (top row, scale bar = 100 nm) with the corresponding tomogram interpretations (bottom row). (**G** to **J**) Violin plots showing the distribution of membrane overlap in arbitrary units (AU) for upright and reversed spikes (bottom row). The number of spikes is given in parenthesis. The black bar is the mean of the distribution. ***, $p < 0.001$. Distribution of the spikes on the viral surface included in the analysis (top row). In panels (G) and (I), all spike positions were included. In panels (H) and (J), spikes with high latitude were excluded, because the missing wedge in Fourier space leads to essentially undetectable membrane (when oriented perpendicular to the optical axis of the microscope) for those spikes. (G) Analysis for all tomograms and all upright and reversed spikes. (H) Analysis for all tomograms and spikes with latitude smaller than |±45°|. (I) Analysis for a single tomogram and all upright and reversed spikes. (J) Analysis for a single tomograms and spikes with latitude smaller than |±45°|. The data shown in (I) and (J) are for the same tomogram.

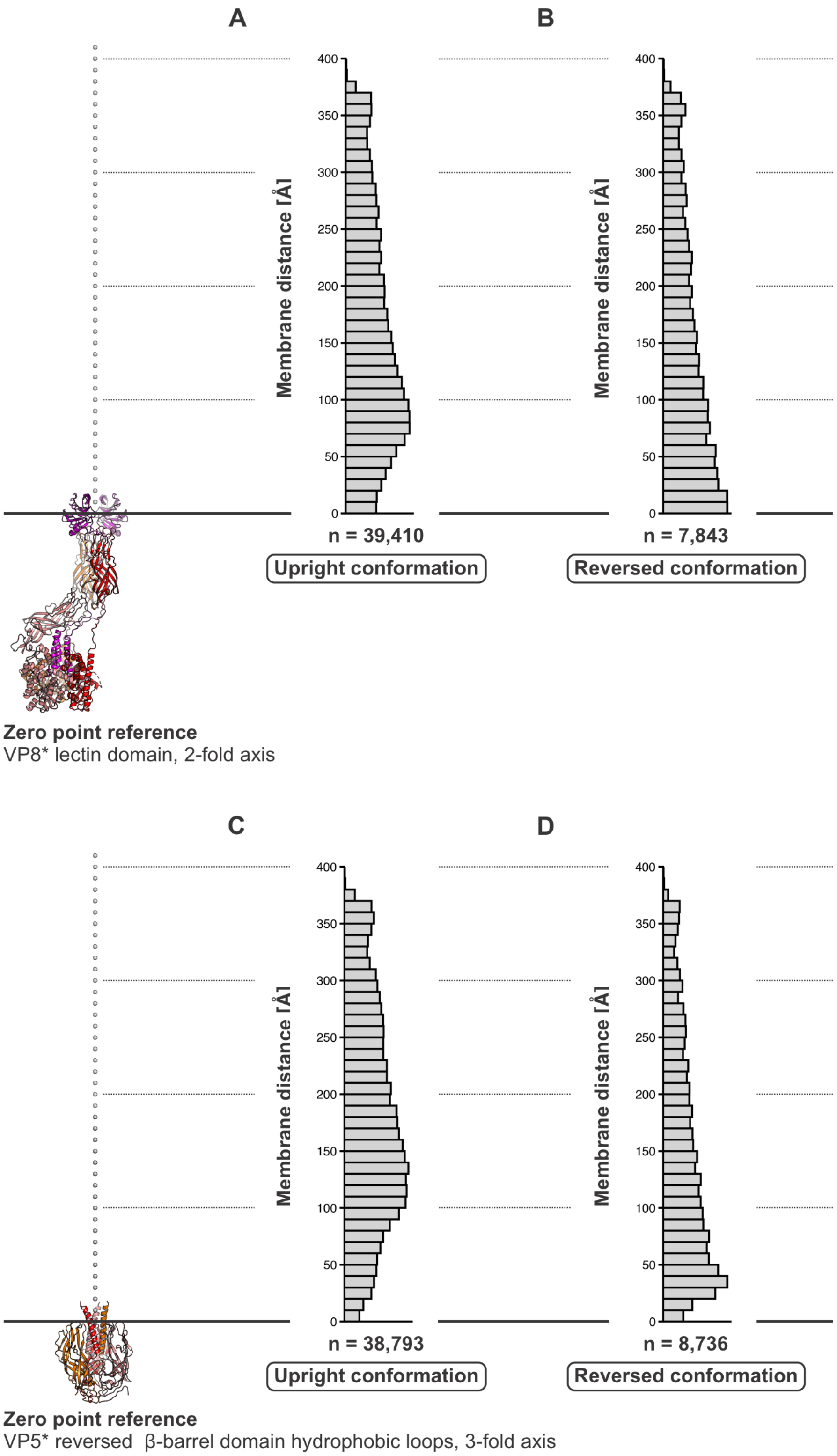


**Fig. S8. Distances between spikes and segmented membrane.** See Materials and Methods for details. (**A** and **B**) Histograms for upright and reversed spike positions, respectively, with the upright conformation as reference (zero point and 2-fold axis). (**C** and **D**) Histograms for upright and reversed spike positions, respectively, with the reversed conformation as reference (zero point and 3-fold axis).

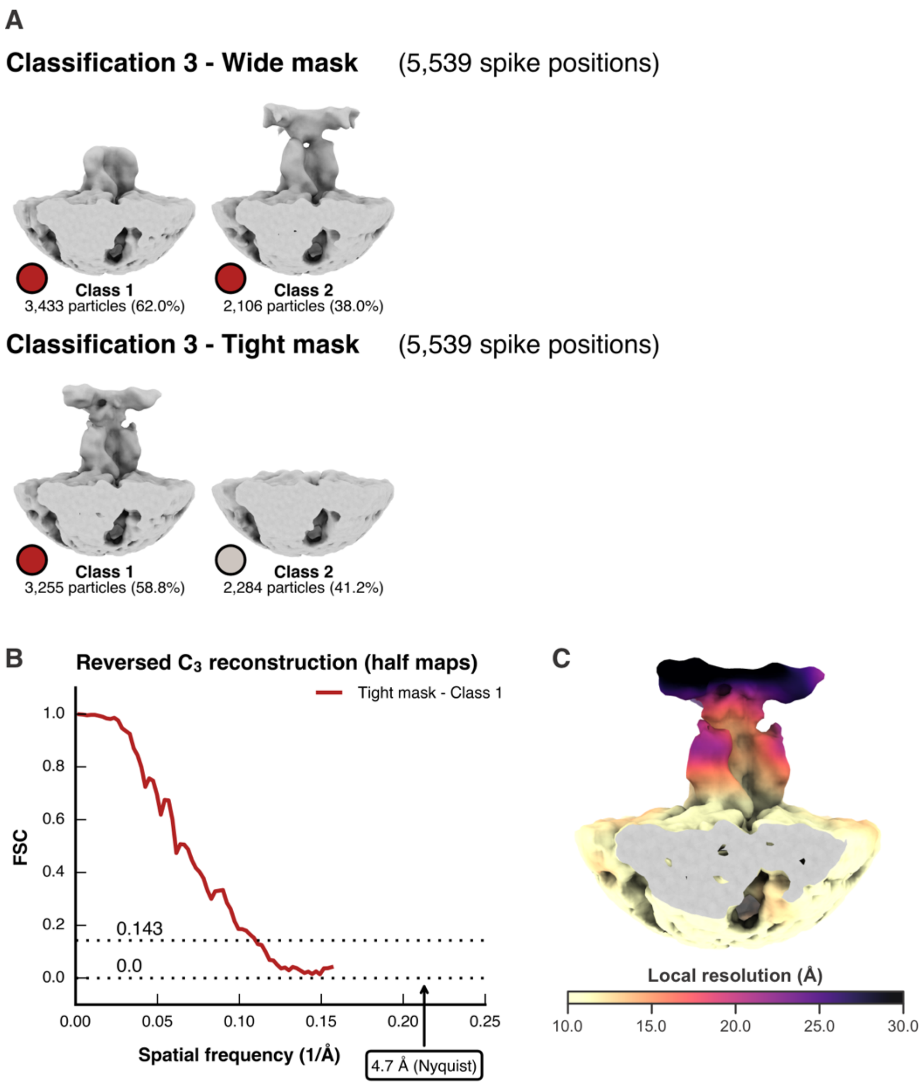


**Fig. S9. 3D classification 3 of a selected (high membrane overlap) set of VP5* reversed spikes.** (**A**) 3D classification 3 was done with RELION without alignment and $C_3$ symmetry imposed, and maps shown here were reconstructed with M and filtered according to local resolution (see Materials and Methods). Top, classification with a wide mask. Bottom, classification with a tight mask. Spike position configuration is color coded: reversed, dark red; empty (unoccupied, no VP4 present), gray. The map of class 1 obtained after classification with a tight mask is shown in Fig. 3A. (**B**) Fourier shell correlation (FSC) curves calculated from the half maps. (**C**) Local resolution mapped on the Fig. 3A reconstruction.

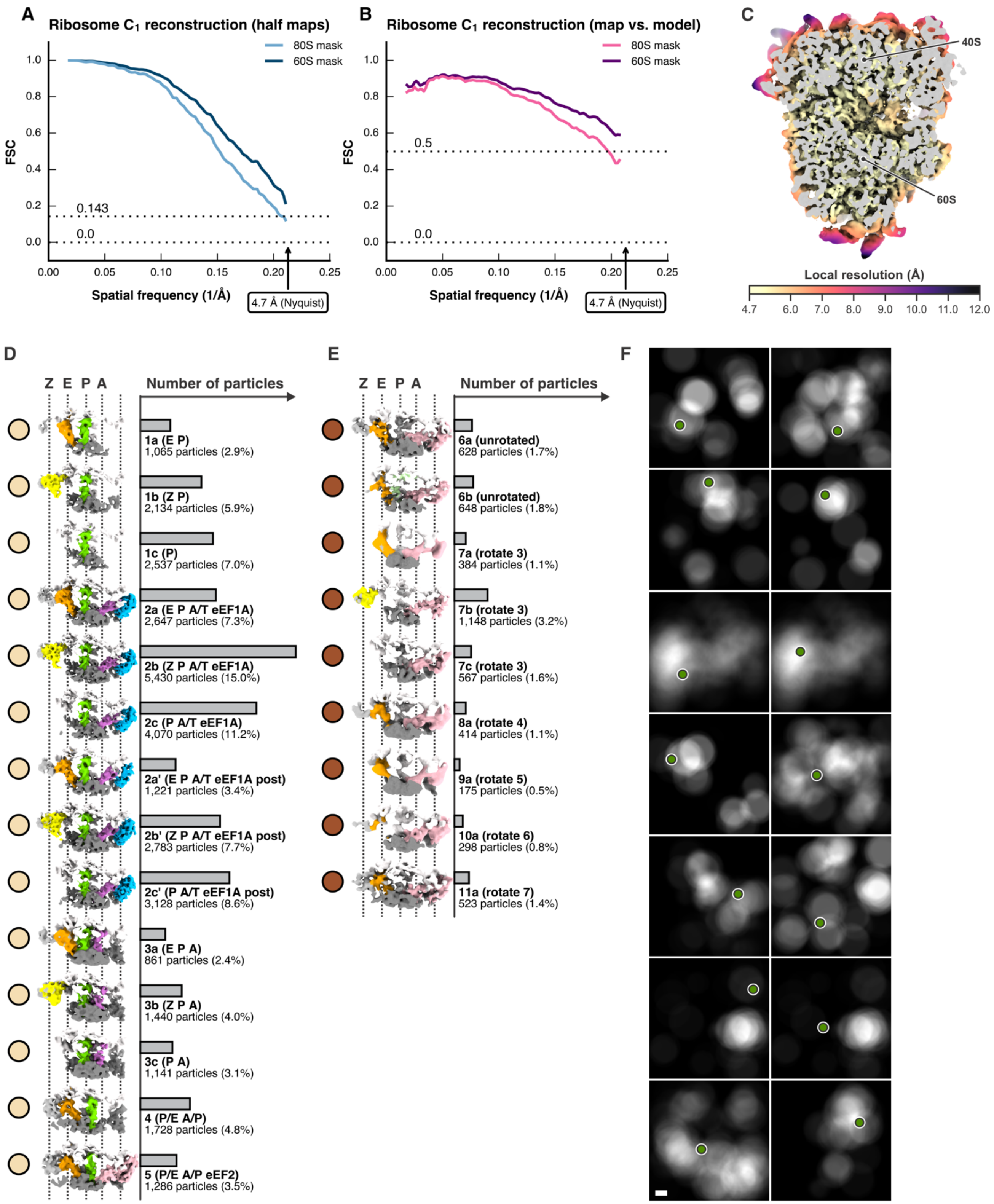


**Fig. S10. Analysis and supervised classification of ribosomes.** (**A**) Fourier shell correlation (FSC) curves after M refinement (with the initial 80S and final 60S masks, respectively) calculated from the half maps of the consensus reconstructions. (**B**) FSC between the full maps and a high-resolution model (PDB-ID pdb_00009p72). (**C**) Local resolution mapped on the consensus reconstruction (60S mask). (**D**

and **E**) Ribosomal state distribution determined by supervised classification (see Materials and Methods for details). For each of the 23 states, a reconstruction from the assigned subtomograms is shown, masked to focus on the tRNA and elongation factor binding sites between the 60S (white) and 40S (gray) subunits. The ribosomes states are named according to previous work (*20*) and so are the tRNAs and elongation factors colored (Z tRNA: yellow; E tRNA: orange; P tRNA: green; A tRNA: magenta; eEF1A: blue; eEF2 pink. The approximate positions of the Z, E, P, and A sites are indicated by dashed lines. Beige dots indicate states associated with translating ribosomes (D), brown dots indicated non-translating states (E). (**F**) Ribosome density in tomograms (see Materials and Methods) in relation to the positions of 14 virions assigned to the "Transcription (& translation?)" entry step (Fig. 2F and table S2). Only the z section at the center of each virion is shown from the 3D density maps. The position of the virions is indicated by green dots. The scale bar corresponds to 100 nm. Note that this analysis contains an additional 7 virions that were not included in the analysis of table S2, because they were initially not picked by the picking algorithm.

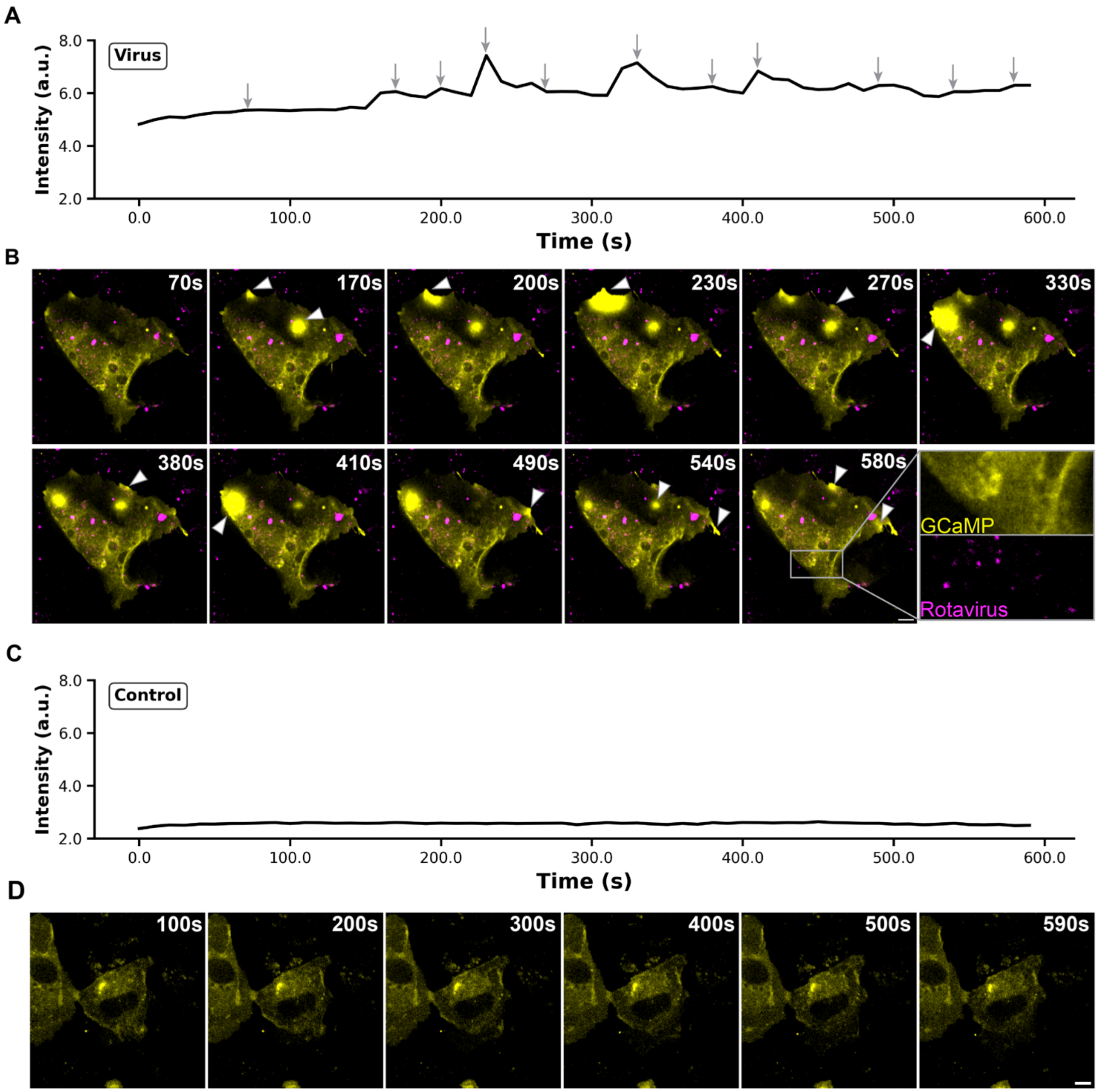


**Fig. S11. Rotavirus induces localized calcium transients in BSC-1 cells.** (**A**) GCaMP fluorescence intensity over time in a representative BSC-1 cell after 5 min incubation with rotavirus. Arrows indicate transient calcium spikes. (**B**) Confocal microscopy images corresponding to the different time points indicated with gray arrows in (A). GCaMP calcium indicator is shown in yellow and fluorescently labeled rotavirus in magenta. White arrowheads indicate regions of elevated GCaMP fluorescence corresponding to calcium transients. Inset shows magnified view of both channels separately in a region of the cell. Scale bar, 10 µm. (**C**) GCaMP fluorescence intensity over time in a representative uninfected control BSC-1 cell showing stable baseline calcium levels. (**D**) Time-lapse images of the control cell shown in (C). Scale bar, 10 µm.

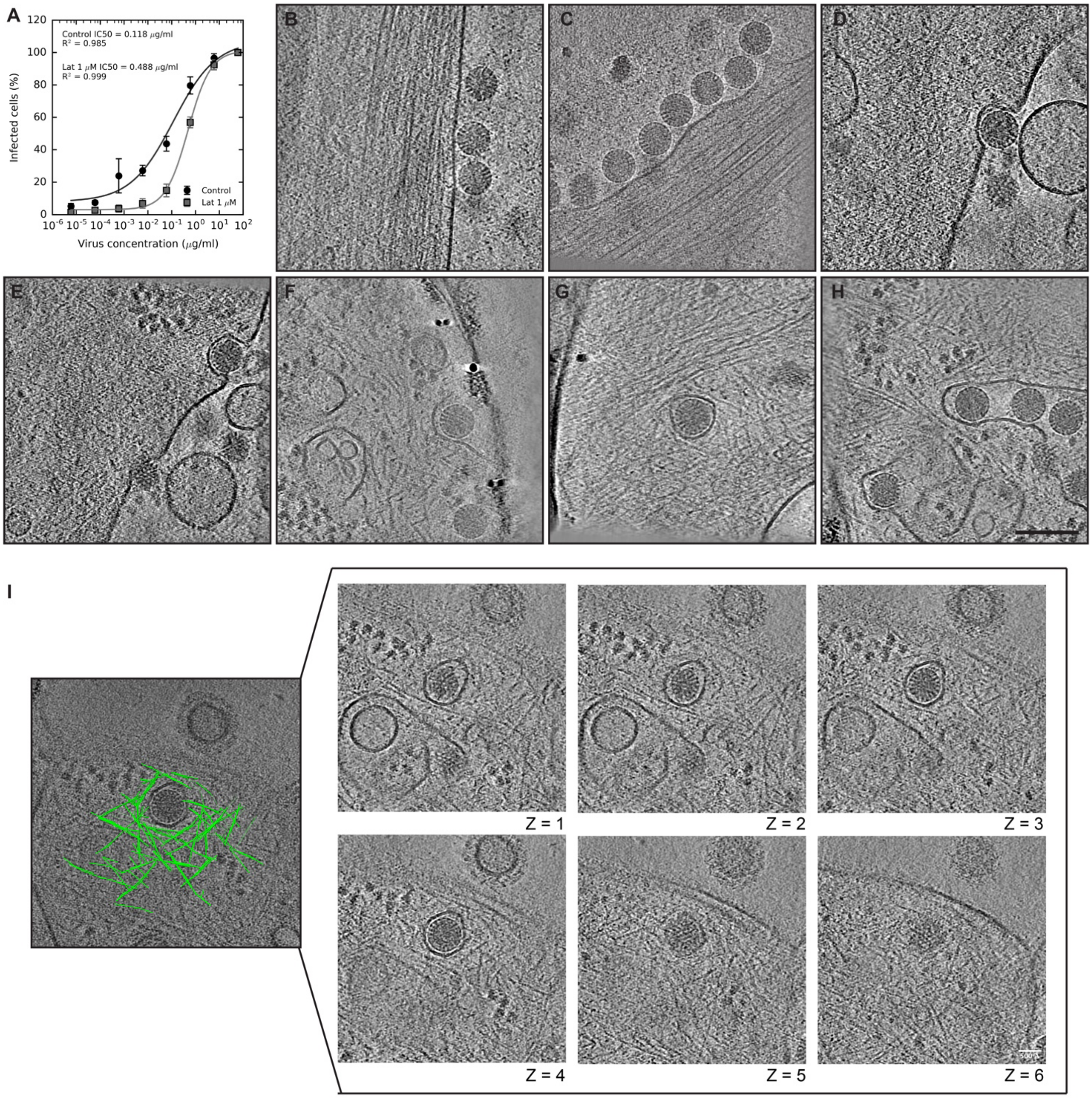


**Fig. S12. Actin reorganization at rotavirus entry sites.** (**A**) Rotavirus infectivity in the absence (control, black circles) or presence (gray squares) of 1 μM latrunculin A, an actin polymerization inhibitor. Data represent the mean ± standard deviation from three independent experiments. We carried out a two-way analysis of variance (ANOVA) to determine the significance of latrunculin A treatment, using the standard method (OLS) within the statsmodels Python library. The p-value was $<0.001$. (**B** to **H**) Representative tomographic slices of rotavirus-infected BSC-1 cells showing viruses close to cytoskeletal networks. In panels (D) to (H), branched actin networks surround entering virus particles. The scale bar corresponds to 100 nm (H). (**I**) Manual segmentation of actin filaments (green) surrounding a rotavirus particle. Serial tomographic slices along the z axis of the same virus particle are shown, scale bar = 50 nm.

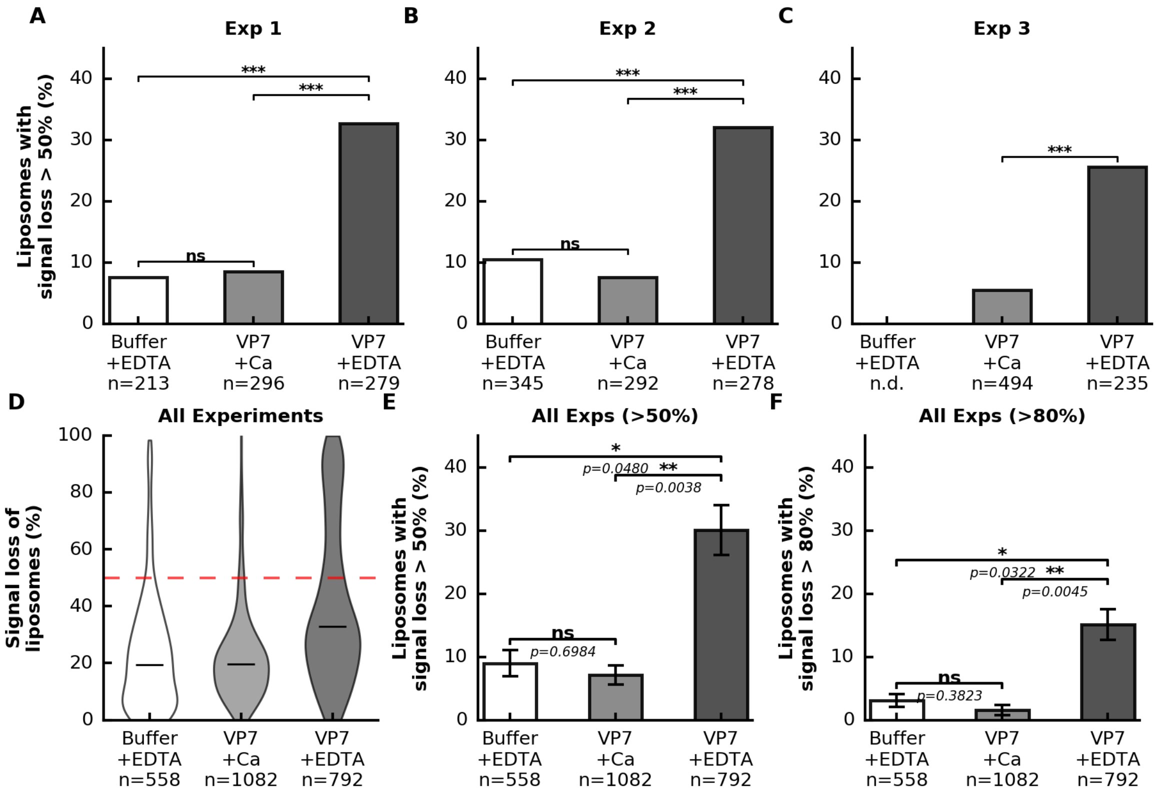


**Fig. S13. Quantification of TIRF microscopy data.** (**A** to **C**) Percentage of liposomes with carboxyfluorescein (CF) signal loss greater than 50% after 30 min incubation with EDTA buffer only (white bars), VP7 trimer in $Ca^{2+}$ buffer (light grey bars), and VP7 monomer in EDTA buffer (dark grey bars), for three independent experiments (A to C). Total number of liposomes is shown at the bottom of each bar. Error bars represent the standard deviation of the mean (SD, n = 3 independent experiments); Two sided t-test statistical analysis was used between samples. *, $p < 0.05$; **, $p < 0.01$; ***, $p < 0.001$; ns, not significant. (**D**) Liposome signal loss (%) for all liposomes in the three independent experiments. Total number of liposomes per condition (n) is shown for each sample. The red dotted line indicates the cut off of 50%. (**E** and **F**) Percentage of liposomes with signal loss of more than 50% (E) or 80% (F) after 30 min incubation with EDTA buffer only (white bar), VP7 trimer in $Ca^{2+}$ buffer (light grey bar) or VP7 monomer (dark grey bar). Plotted data are from three independent experiments. Total number of liposomes (n) is shown for each condition. Error bars represent the standard deviation of the mean (SD, n = 3 independent experiments); t-test statistical analysis was used between samples, with p values indicated.

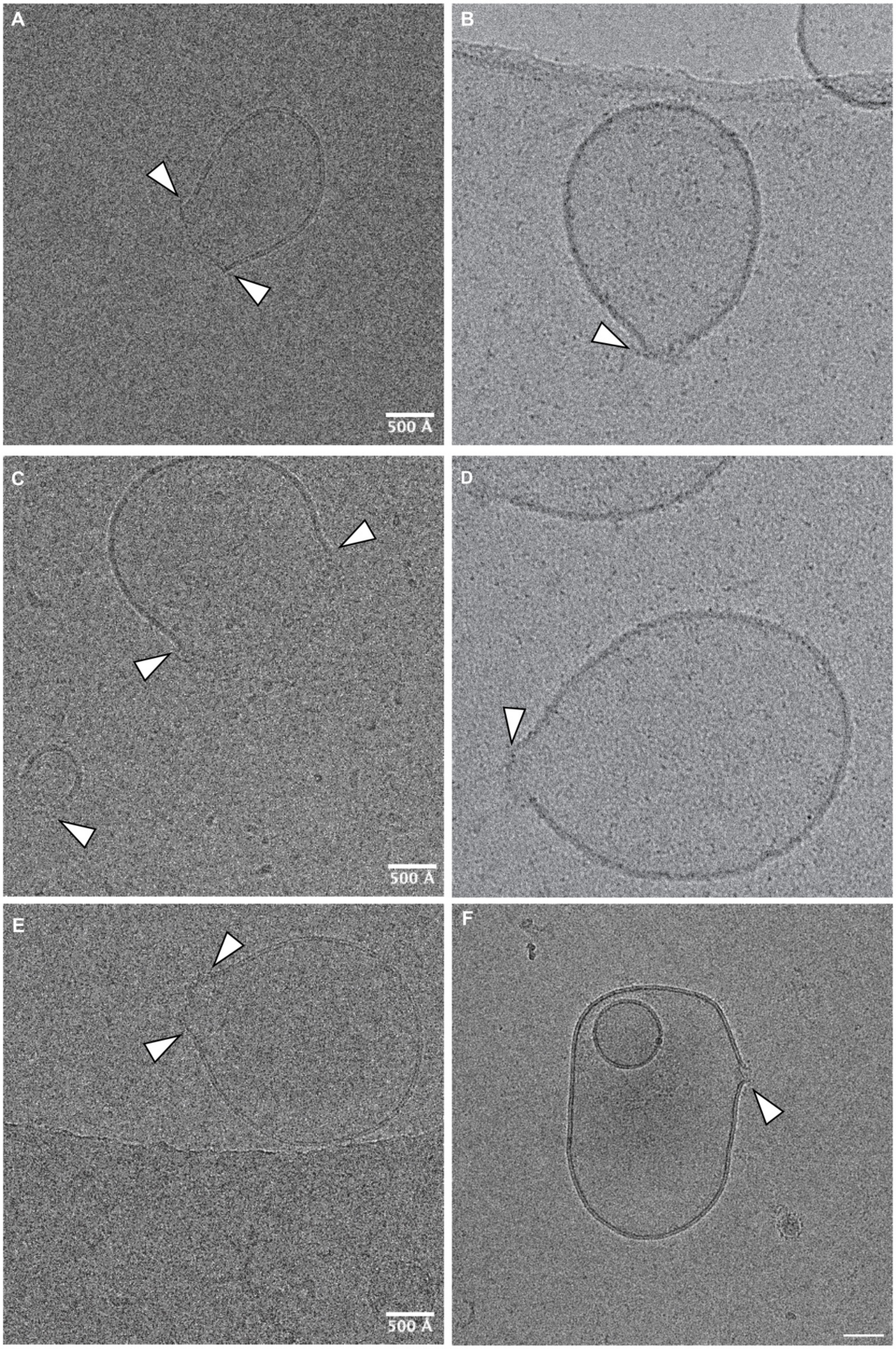


**Fig. S14. cryo-EM of VP7-disrupted liposomes.** (**A** to **F**) Representative cryo-EM images of liposomes after 30 min incubation with VP7 monomer in EDTA buffer. White arrows show perforation of the liposomes. Scale bar = 500 Å.

**A**

| Sample | FFU/mL | Average FFU/mL | Std Dev |
|---|---|---|---|
| TLP | 6.33e+11<br>2.50e+12<br>8.75e+11 | 1.34e+12 | 1.02e+12 |
| TLP + Trypsin Inhibitor | 8.00e+11<br>5.17e+11<br>1.00e+12 | 7.72e+11 | 2.43e+11 |

**B**

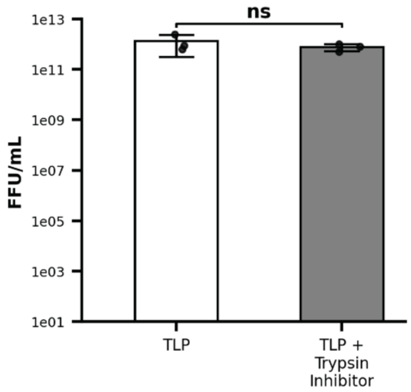


**Fig. S15. Infectivity assay of virus particles incubated with trypsin inhibitor.** (**A**) The table shows the infectivity values as focus-forming units per milliliter (FFU/ml) for TLP sample and TLP incubated with trypsin inhibitor for three independent experiments, determined as described in Methods. (**B**) Average and standard deviation (Std Dev) of the three independent experiments are displayed in a bar graph with the focus-forming unit concentration (FFU/ml) for TLPs in white and TLPs incubated with trypsin inhibitor in gray. A t-test statistical analysis was performed (p value = 0.4; ns, not significant).

#### Table S1. Cryo-ET data collection and tilt series refinement.

| | Dataset 1<br>BSC-1 + RRV | Dataset 2<br>BSC-1 + RRV | Dataset 3<br>BSC-1 + RRV | Dataset 4<br>BSC-1 + RRV | Dataset 5<br>BSC-1 + RRV | Dataset 6<br>BSC-1 + RRV | Dataset 7<br>BSC-1 + RRV |
|---|---|---|---|---|---|---|---|
| **Data collection** | | | | | | | |
| Electron microscope | Titan Krios G3i | Titan Krios G3i | Titan Krios G3i | Titan Krios G3i | Titan Krios G3i | Titan Krios G3i | Titan Krios G3i |
| Energy filter width (eV) | 10 | 10 | 10 | 10 | 10 | 10 | 10 |
| Camera | Falcon 4i | Falcon 4i | Falcon 4i | Falcon 4i | Falcon 4i | Falcon 4i | Falcon 4i |
| Magnification | 53,000 | 53,000 | 53,000 | 53,000 | 53,000 | 53,000 | 53,000 |
| Voltage (kV) | 300 | 300 | 300 | 300 | 300 | 300 | 300 |
| Defocus range (µm) * | 3.0–5.0 | 3.0–5.0 | 3.0–5.0 | 3.0–5.0 | 3.0–5.0 | 3.0–5.0 | 3.0–5.0 |
| # of tilt series (TS) | 95 | 131 | 152 | 93 | 127 | 190 | 50 |
| Tilt angles † | +60° to -60°, 3° | +60° to -60°, 3° | +60° to -60°, 3° | +60° to -60°, 3° | +60° to -60°, 3° | +60° to -60°, 3° | +60° to -60°, 3° |
| # of tilt images (TI) | 41 | 41 | 41 | 41 | 41 | 41 | 41 |
| Dose per TI ($e^-/Å^2$) | 3.7–3.9 | 3.7–3.9 | 3.7–3.9 | 3.7–3.9 | 3.7–3.9 | 3.7–3.9 | 3.7–3.9 |
| Pixel size (Å) ‡ | 2.349 | 2.359 | 2.359 | 2.350 | 2.350 | 2.350 | 2.350 |
| **M refinement ($I_2$)** | | | | | | | |
| Symmetry imposed | $I_2$ | $I_2$ | $I_2$ | $I_2$ | $I_2$ | $I_2$ | $I_2$ |
| # of RRV | 2,150 | 1,574 | 557 | 1,570 | 1,036 | 1,321 | 357 |
| Box size (pixels) | 512 | 512 | 512 | 512 | 512 | 512 | 512 |
| True pixel size (Å) § | 2.325 | 2.320 | 2.317 | 2.316 | 2.323 | 2.319 | 2.319 |
| Map resolution (Å) \|\| | 4.7 (Nyquist) | 4.7 (Nyquist) | 4.7 (Nyquist) | 4.7 (Nyquist) | 4.7 (Nyquist) | 4.8 | 8.8 |
| Map resolution (Å) ¶ | 4.7 (Nyquist) | 4.7 (Nyquist) | 4.9 | 4.9 | 4.9 | 5.1 | 8.9 |
| **M refinement ($C_1$)** | | | | | | | |
| Symmetry imposed | $C_1$ | $C_1$ | $C_1$ | $C_1$ | $C_1$ | $C_1$ | $C_1$ |
| # of RRV spikes | 129,000 | 94,440 | 33,420 | 94,200 | 62,160 | 79,260 | 21,420 |
| Box size (pixels) | 512 | 512 | 512 | 512 | 512 | 512 | 512 |
| Map resolution (Å) # | 5.9 | 4.8 | 7.6 | 7.3 | 7.2 | 7.9 | 11.0 |

* Approximate range of underfocus.
† Collected with a dose-symmetric bidirectional tilt scheme. The minimal, maximal tilt angle, and the increment between adjacent tilt angels are given.
‡ Pixel size used for processing of each dataset.
§ Pixel size determined by scaling the final map to a high-resolution cryo-EM reconstruction.
|| Resolution where the Fourier shell correlation (FSC) between half-maps drops below 0.143 after applying a mask, made from a low-resolution envelope of the proteinaceous triple layers (VP2, VP6, VP7) and the VP4 spikes in upright conformation of the virus.
¶ Resolution where the FSC between the final map and a high-resolution cryo-EM reconstruction drops below 0.5 after applying a mask, made from a low-resolution envelope of the proteinaceous triple layers (VP2, VP6, VP7) and the VP4 spikes in upright conformation of the virus.
### Resolution where the FSC between half-maps drops below 0.143 after applying a spherical mask with a radius of 193 Å.

#### Table S2. Analysis of particles assigned to distinct entry steps.

| | N * | t/d † | # of VP7 ‡ | VP7 lost (%) § | u / r / e \|\| | Reversed (%) ¶ | m # | 80S-n<br>80S-t ** | Transl. (%) †† |
|---|---|---|---|---|---|---|---|---|---|
| **Attachment**<br>(Fig. 2A) | 22 | 57.4 ± 5.8<br>2.6 ± 5.8 | 746 / 780 | 4.4 | 33.9 ± 11.1<br>9.0 ± 4.2<br>14.5 ± 6.0 | 21.0 | 4.0 ± 4.0 | 0.1 ± 0.3<br>1.0 ± 2.8 | 91.7 |
| **Endocytosis**<br>(Fig. 2B) | 12 | 57.2 ± 4.6<br>2.8 ± 4.6 | 744 / 780 | 4.6 | 20.7 ± 11.9<br>15.8 ± 9.7<br>20.8 ± 8.6 | 43.4 | 14.6 ± 4.0 | 0.2 ± 0.4<br>5.7 ± 8.9 | 91.7 |
| **Membrane permeabilization**<br>(Fig. 2C) | 28 | 53.9 ± 6.8<br>6.1 ± 6.8 | 701 / 780 | 10.1 | 15.8 ± 10.5<br>14.8 ± 7.1<br>23.4 ± 7.5 | 48.4 | 16.0 ± 6.0 | 1.0 ± 1.6<br>6.4 ± 8.3 | 86.1 |
| **Membrane perforation**<br>(Fig. 2D) | 9 | 28.9 ± 9.9<br>31.1 ± 9.9 | 376 / 780 | 51.9 | 8.6 ± 5.9<br>6.9 ± 4.5<br>13.4 ± 6.4 | 44.6 | 11.0 ± 4.3 | 0.9 ± 1.0<br>4.3 ± 9.2 | 83.0 |
| **DLP release**<br>(Fig. 2E) | 8 | 19.5 ± 9.9<br>40.0 ± 9.9 | 254 / 780 | 67.5 | 5.1 ± 5.8<br>7.6 ± 5.0<br>6.8 ± 4.4 | 59.8 | 7.2 ± 4.4 | 0.4 ± 0.5<br>5.9 ± 4.7 | 94.0 |
| **Transcription (& translation?)**<br>(Fig. 2F) | 7 | 20.4 ± 11.0<br>39.6 ± 11.0 | 266 / 780 | 66.0 | 5.3 ± 5.8<br>4.9 ± 6.6<br>10.3 ± 9.3 | 47.9 | 0.5 ± 0.9 | 3.7 ± 5.0<br>17.9 ± 18.3 | 82.8 |

* Number of particles included in the analysis for each entry step. Because each tomogram represented a single-time snapshot, there are fewer particles in the perforation and release steps, which are relatively fast (*15*) compared to the preceding steps.
† Average ± standard deviation of the number of asymmetric subunits per particle (total of 60) classified as TLP (t, VP7 present) or DLP (d, VP7 absent).
‡ Average number of VP7 subunits per rotavirus particle, out of a maximum of 780.
§ Average degree of uncoating per rotavirus particle, as % of VP7 lost. The number is probably an underestimate, as the classifier would have interpreted as VP7 any noise or random presence of other protein in the relevant volume around the particle surface. From the particles we expect to be fully uncoated in the last two steps, a rough correction would adjust these numbers upward by about 1.5.
|| Average ± standard deviation of the number of upright (u), reversed (r), and empty (e) spike position per particle.
¶ Average percentage of reversed spike positions, excluding empty ones.
### Average total membrane overlap of all spike positions per rotavirus particle in arbitrary units (see fig. S7).
** Average number of non-translating (80S-n) and translating (80S-t) 80S ribosomes within a distance of 1500 Å from the particle center.
†† Average percent of translating ribosomes.

**Movie S1. 3D movie of the reconstructed tomogram shown in Fig. 1D.** The movie shows the raw reconstructed tomogram viewed along the Z axis. The segmented membrane is rendered in gray, translating and non-translating ribosomes in beige and brown, respectively, and rotavirus proteins are shown as follows: VP2A and VP2B in blue and cyan, respectively; VP6 in green; VP7 in yellow; and surface spikes in the upright (light red) and reversed (dark red) conformations.

**Movie S2. Molecular interpretation of the tomogram sections shown in Fig. 2 with varying degree of transparency of the membrane segmentation.** VP6 (green), VP7 (yellow), VP5*/VP8* in upright conformation (light red), and VP5* in reversed conformation (dark red). Segmented membranes are shown in gray, translating and non-translating ribosomes in beige and brown, respectively.

**Movie S3. Ribosomal state distribution determined by supervised classification.** See Materials and Methods for details. For each of the 23 states, a reconstruction from the assigned subtomograms is shown. 60S (white) and 40S (gray) subunits. The ribosomes states are named according to previous work (*20*), as are the tRNAs and elongation factors, colored Z tRNA: yellow; E tRNA: orange; P tRNA: green; A tRNA: magenta; eEF1A: blue; eEF2 pink. Beige dots indicate states associated with translating ribosomes, brown dots indicated non-translating states. The bar on top corresponds to the relative number of particles in each state.

**Movie S4. Tomogram reconstruction and membrane segmentation of VP7-disrupted liposomes.** The movie shows raw reconstructed tomograms viewed along the Z axis for three examples of liposomes incubated and disrupted by monomeric VP7 (EDTA buffer). The membrane was segmented with MemBrain and is shown in gray. Scale bar = 50 nm.

**Data S1. Overlap of spike positions with segmented membrane for each tomogram.** Violin plots showing the distribution of membrane overlap in arbitrary units (AU) for upright and reversed spikes for each tomogram containing rotavirus (537 tomograms). The number of spikes is given in parenthesis. The black bar is the mean of the distribution. Only spikes with latitude smaller than ±45° were included in the analysis (see also fig. S7, G to J). The inset horizonal bar plot shows the total detected membrane volume for each tomogram.
