## Supplementary material for "Mechanism of membrane perforation in rotavirus cell entry": data file (membrane overlap)

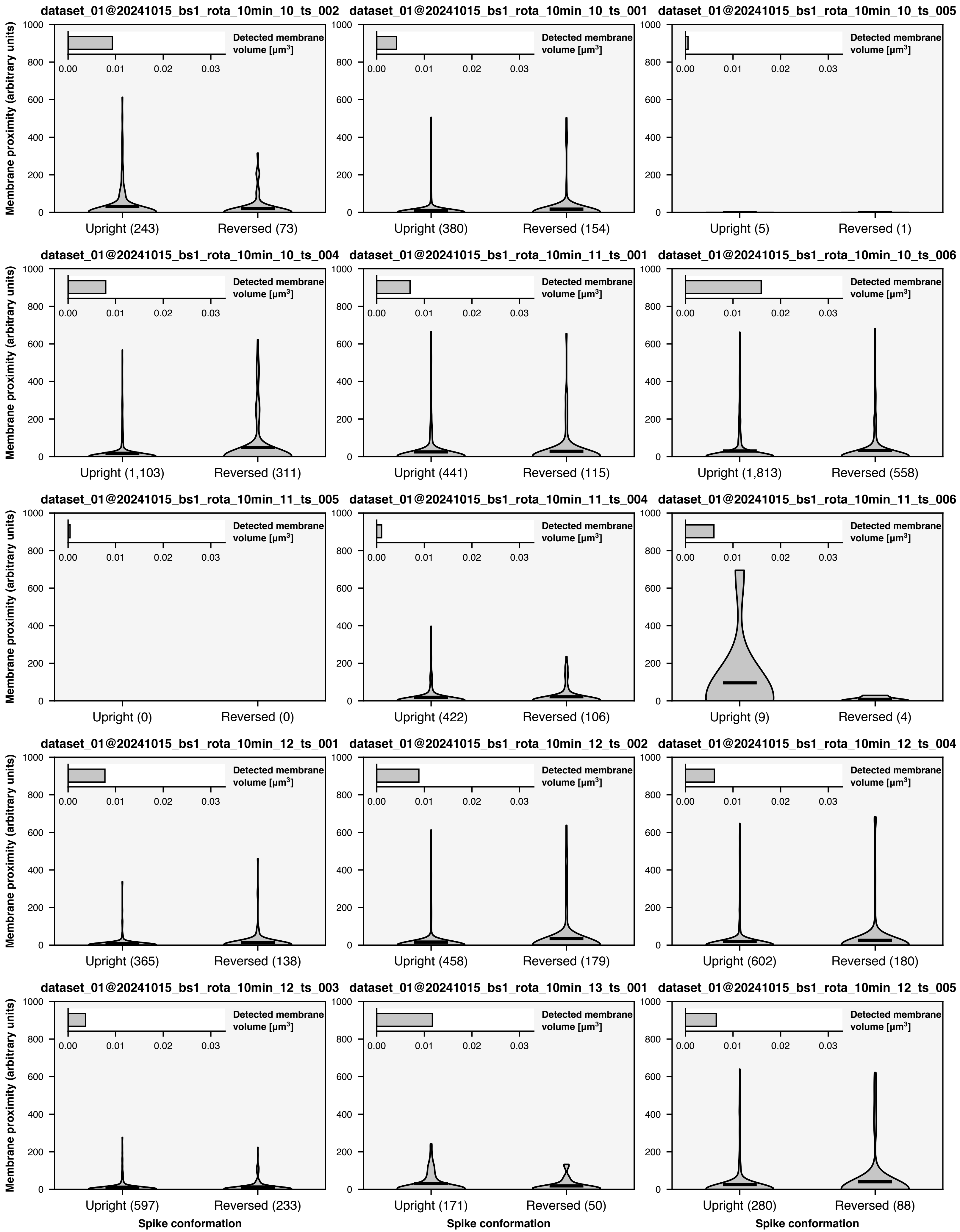
dataset_01@20241015_bs1_rota_10min_10_ts_002
Detected membrane volume [μm³]
Membrane proximity (arbitrary units)
Upright (243)
Reversed (73)
dataset_01@20241015_bs1_rota_10min_10_ts_001
Upright (380)
Reversed (154)
dataset_01@20241015_bs1_rota_10min_10_ts_005
Upright (5)
Reversed (1)
dataset_01@20241015_bs1_rota_10min_10_ts_004
Upright (1,103)
Reversed (311)
dataset_01@20241015_bs1_rota_10min_11_ts_001
Upright (441)
Reversed (115)
dataset_01@20241015_bs1_rota_10min_10_ts_006
Upright (1,813)
Reversed (558)
dataset_01@20241015_bs1_rota_10min_11_ts_005
Upright (0)
Reversed (0)
dataset_01@20241015_bs1_rota_10min_11_ts_004
Upright (422)
Reversed (106)
dataset_01@20241015_bs1_rota_10min_11_ts_006
Upright (9)
Reversed (4)
dataset_01@20241015_bs1_rota_10min_12_ts_001
Upright (365)
Reversed (138)
dataset_01@20241015_bs1_rota_10min_12_ts_002
Upright (458)
Reversed (179)
dataset_01@20241015_bs1_rota_10min_12_ts_004
Upright (602)
Reversed (180)
dataset_01@20241015_bs1_rota_10min_12_ts_003
Upright (597)
Reversed (233)
dataset_01@20241015_bs1_rota_10min_13_ts_001
Upright (171)
Reversed (50)
dataset_01@20241015_bs1_rota_10min_12_ts_005
Upright (280)
Reversed (88)
Spike conformation

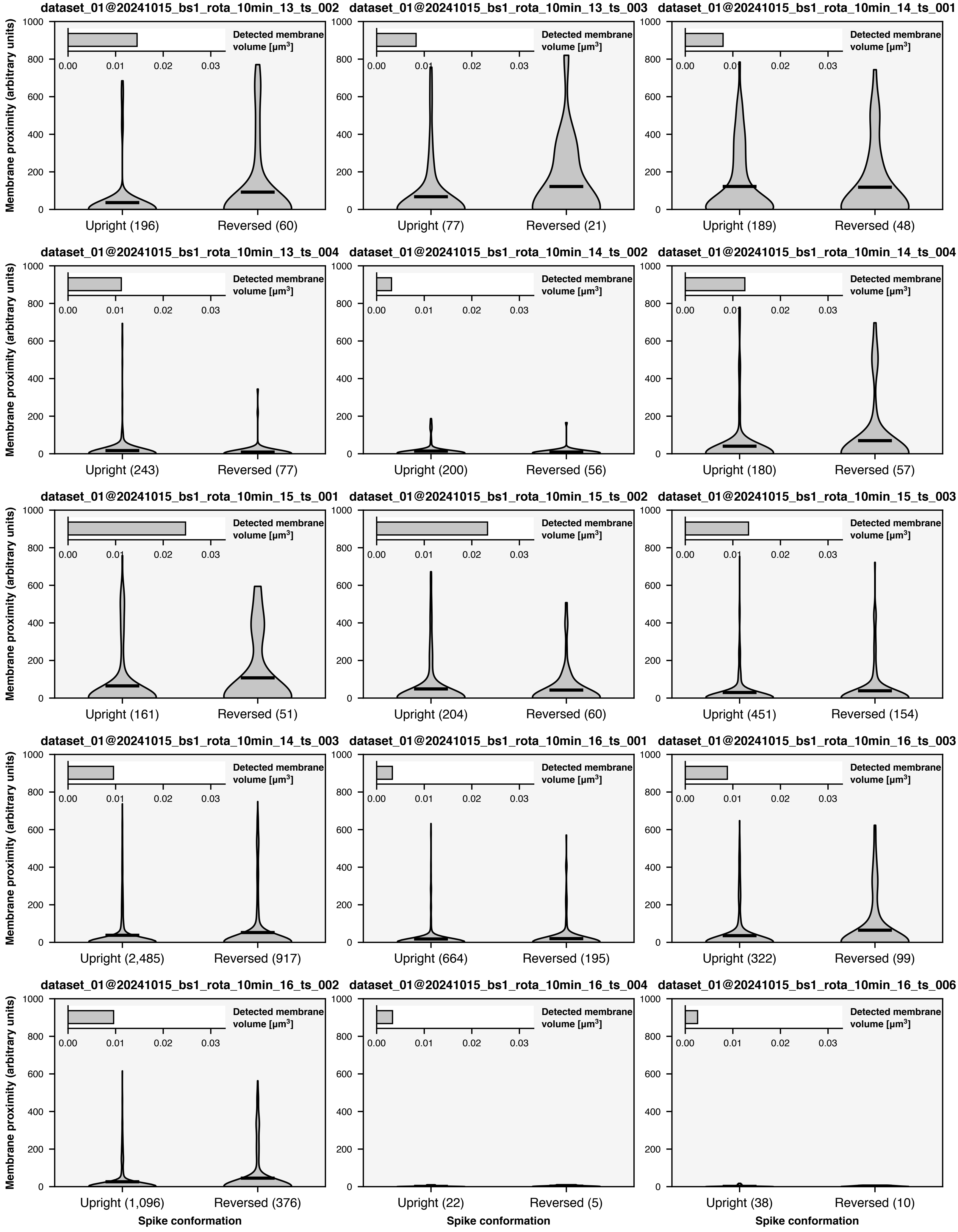
dataset_01@20241015_bs1_rota_10min_13_ts_002
dataset_01@20241015_bs1_rota_10min_13_ts_003
dataset_01@20241015_bs1_rota_10min_14_ts_001
dataset_01@20241015_bs1_rota_10min_13_ts_004
dataset_01@20241015_bs1_rota_10min_14_ts_002
dataset_01@20241015_bs1_rota_10min_14_ts_004
dataset_01@20241015_bs1_rota_10min_15_ts_001
dataset_01@20241015_bs1_rota_10min_15_ts_002
dataset_01@20241015_bs1_rota_10min_15_ts_003
dataset_01@20241015_bs1_rota_10min_14_ts_003
dataset_01@20241015_bs1_rota_10min_16_ts_001
dataset_01@20241015_bs1_rota_10min_16_ts_003
dataset_01@20241015_bs1_rota_10min_16_ts_002
dataset_01@20241015_bs1_rota_10min_16_ts_004
dataset_01@20241015_bs1_rota_10min_16_ts_006
Membrane proximity (arbitrary units)
Detected membrane volume [μm3]
Upright (196)
Reversed (60)
Upright (77)
Reversed (21)
Upright (189)
Reversed (48)
Upright (243)
Reversed (77)
Upright (200)
Reversed (56)
Upright (180)
Reversed (57)
Upright (161)
Reversed (51)
Upright (204)
Reversed (60)
Upright (451)
Reversed (154)
Upright (2,485)
Reversed (917)
Upright (664)
Reversed (195)
Upright (322)
Reversed (99)
Upright (1,096)
Reversed (376)
Upright (22)
Reversed (5)
Upright (38)
Reversed (10)
Spike conformation

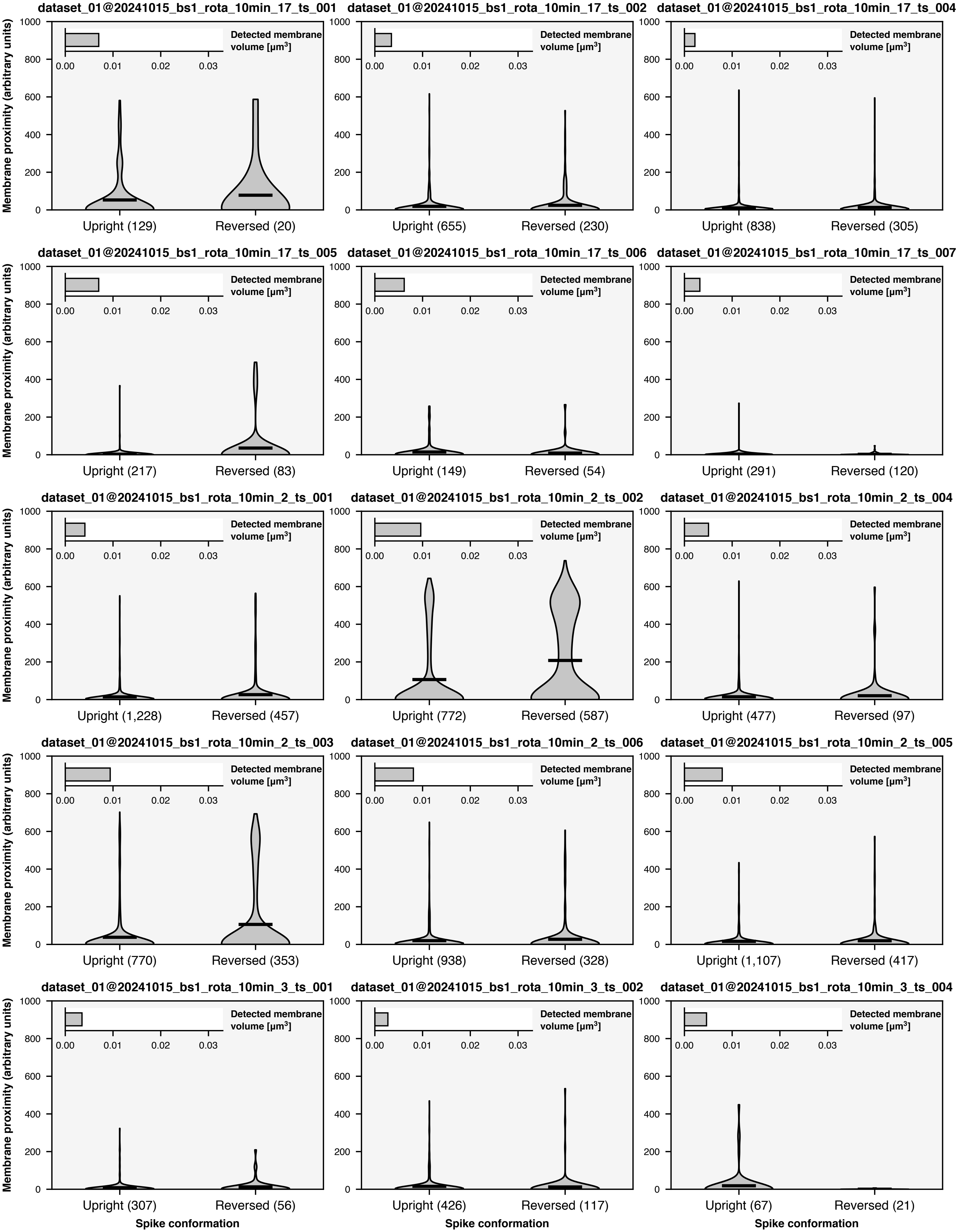

dataset_01@20241015_bs1_rota_10min_17_ts_001
Upright (129)
Reversed (20)
dataset_01@20241015_bs1_rota_10min_17_ts_002
Upright (655)
Reversed (230)
dataset_01@20241015_bs1_rota_10min_17_ts_004
Upright (838)
Reversed (305)
dataset_01@20241015_bs1_rota_10min_17_ts_005
Upright (217)
Reversed (83)
dataset_01@20241015_bs1_rota_10min_17_ts_006
Upright (149)
Reversed (54)
dataset_01@20241015_bs1_rota_10min_17_ts_007
Upright (291)
Reversed (120)
dataset_01@20241015_bs1_rota_10min_2_ts_001
Upright (1,228)
Reversed (457)
dataset_01@20241015_bs1_rota_10min_2_ts_002
Upright (772)
Reversed (587)
dataset_01@20241015_bs1_rota_10min_2_ts_004
Upright (477)
Reversed (97)
dataset_01@20241015_bs1_rota_10min_2_ts_003
Upright (770)
Reversed (353)
dataset_01@20241015_bs1_rota_10min_2_ts_006
Upright (938)
Reversed (328)
dataset_01@20241015_bs1_rota_10min_2_ts_005
Upright (1,107)
Reversed (417)
dataset_01@20241015_bs1_rota_10min_3_ts_001
Upright (307)
Reversed (56)
dataset_01@20241015_bs1_rota_10min_3_ts_002
Upright (426)
Reversed (117)
dataset_01@20241015_bs1_rota_10min_3_ts_004
Upright (67)
Reversed (21)
Membrane proximity (arbitrary units)
0
200
400
600
800
1000
Detected membrane volume [μm³]
0.00
0.01
0.02
0.03
Spike conformation

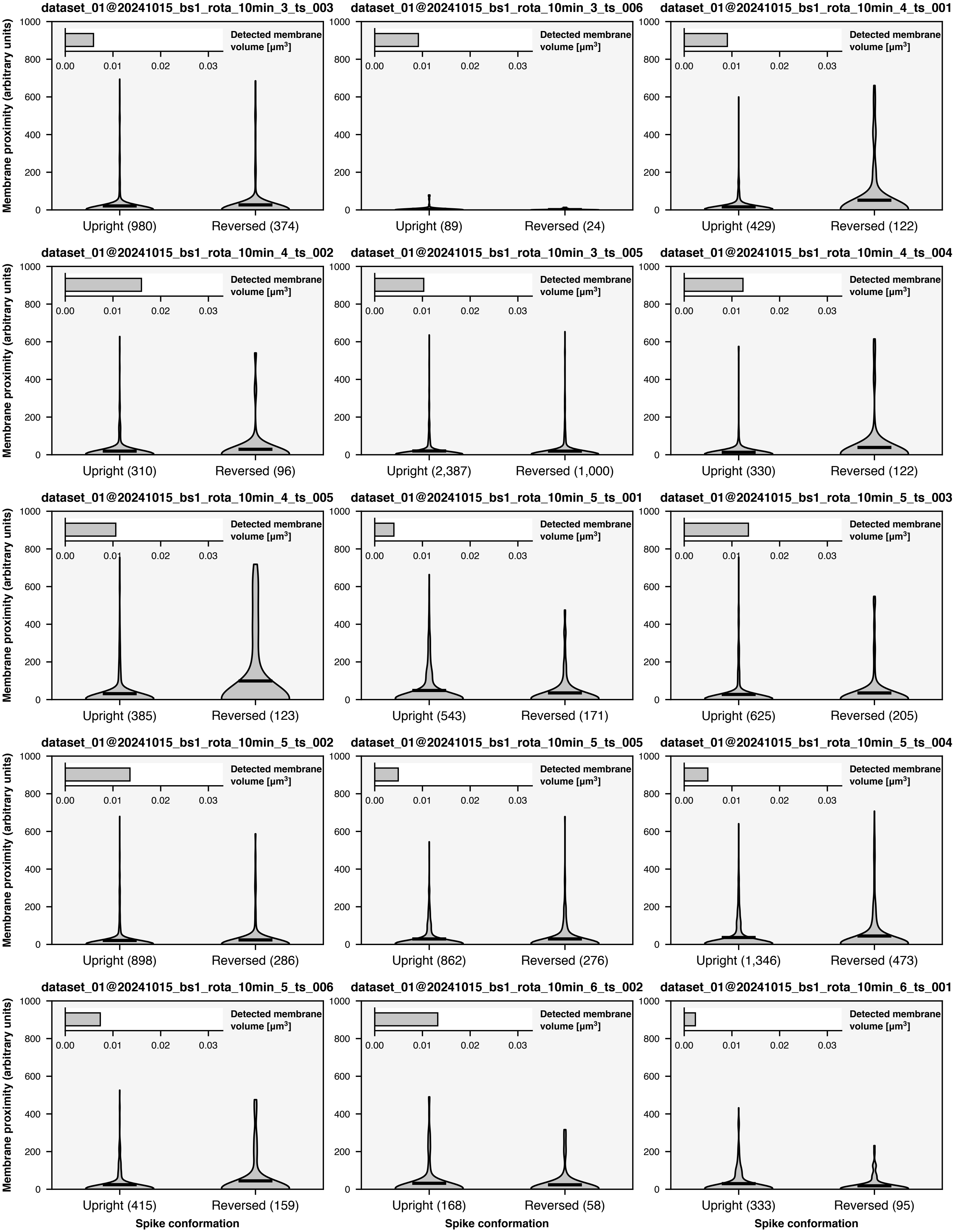
dataset_01@20241015_bs1_rota_10min_3_ts_003
dataset_01@20241015_bs1_rota_10min_3_ts_006
dataset_01@20241015_bs1_rota_10min_4_ts_001
dataset_01@20241015_bs1_rota_10min_4_ts_002
dataset_01@20241015_bs1_rota_10min_3_ts_005
dataset_01@20241015_bs1_rota_10min_4_ts_004
dataset_01@20241015_bs1_rota_10min_4_ts_005
dataset_01@20241015_bs1_rota_10min_5_ts_001
dataset_01@20241015_bs1_rota_10min_5_ts_003
dataset_01@20241015_bs1_rota_10min_5_ts_002
dataset_01@20241015_bs1_rota_10min_5_ts_005
dataset_01@20241015_bs1_rota_10min_5_ts_004
dataset_01@20241015_bs1_rota_10min_5_ts_006
dataset_01@20241015_bs1_rota_10min_6_ts_002
dataset_01@20241015_bs1_rota_10min_6_ts_001
Membrane proximity (arbitrary units)
Detected membrane volume [μm³]
Upright (980)
Reversed (374)
Upright (89)
Reversed (24)
Upright (429)
Reversed (122)
Upright (310)
Reversed (96)
Upright (2,387)
Reversed (1,000)
Upright (330)
Reversed (122)
Upright (385)
Reversed (123)
Upright (543)
Reversed (171)
Upright (625)
Reversed (205)
Upright (898)
Reversed (286)
Upright (862)
Reversed (276)
Upright (1,346)
Reversed (473)
Upright (415)
Reversed (159)
Upright (168)
Reversed (58)
Upright (333)
Reversed (95)
Spike conformation

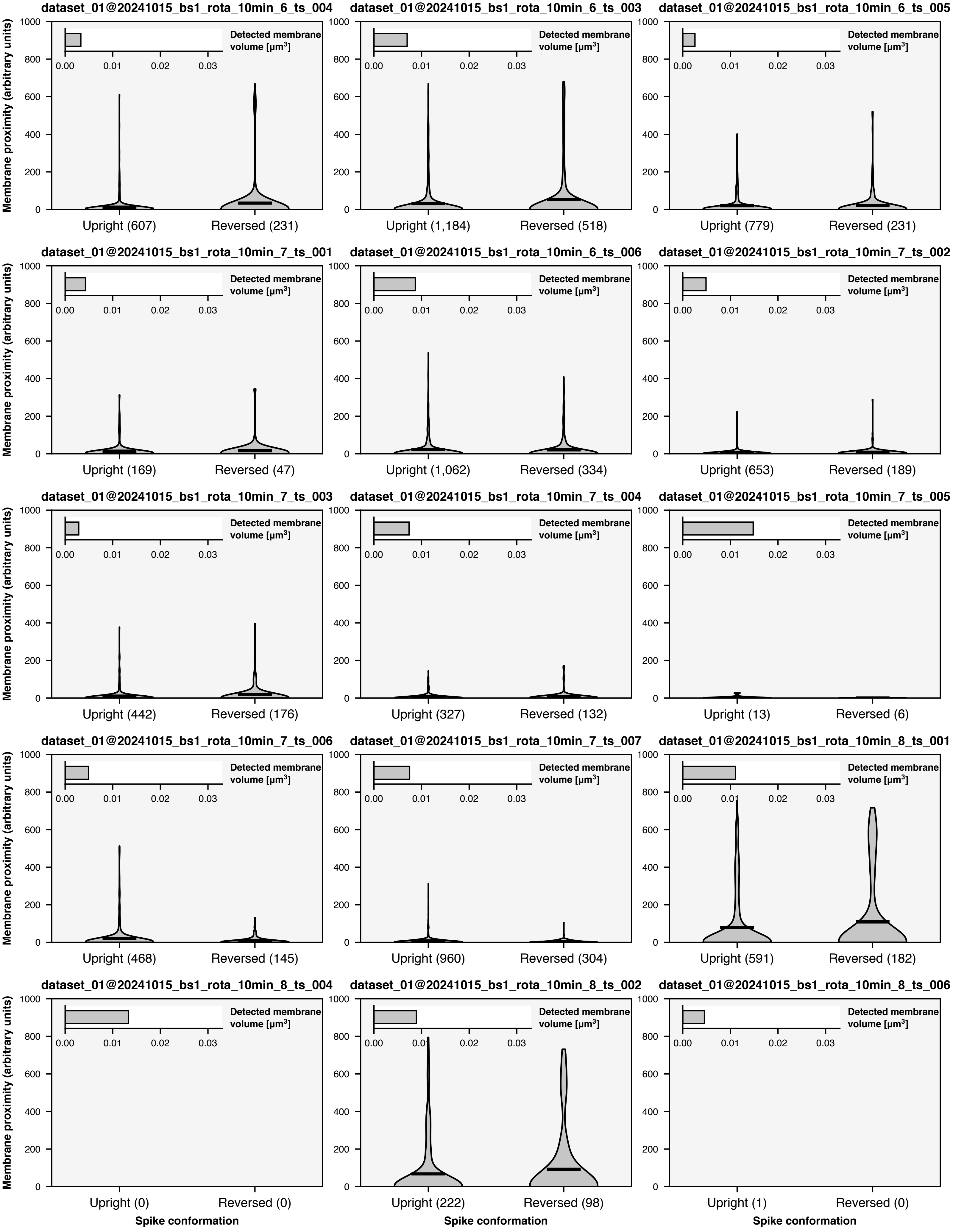
dataset_01@20241015_bs1_rota_10min_6_ts_004
Upright (607)
Reversed (231)
dataset_01@20241015_bs1_rota_10min_6_ts_003
Upright (1,184)
Reversed (518)
dataset_01@20241015_bs1_rota_10min_6_ts_005
Upright (779)
Reversed (231)
dataset_01@20241015_bs1_rota_10min_7_ts_001
Upright (169)
Reversed (47)
dataset_01@20241015_bs1_rota_10min_6_ts_006
Upright (1,062)
Reversed (334)
dataset_01@20241015_bs1_rota_10min_7_ts_002
Upright (653)
Reversed (189)
dataset_01@20241015_bs1_rota_10min_7_ts_003
Upright (442)
Reversed (176)
dataset_01@20241015_bs1_rota_10min_7_ts_004
Upright (327)
Reversed (132)
dataset_01@20241015_bs1_rota_10min_7_ts_005
Upright (13)
Reversed (6)
dataset_01@20241015_bs1_rota_10min_7_ts_006
Upright (468)
Reversed (145)
dataset_01@20241015_bs1_rota_10min_7_ts_007
Upright (960)
Reversed (304)
dataset_01@20241015_bs1_rota_10min_8_ts_001
Upright (591)
Reversed (182)
dataset_01@20241015_bs1_rota_10min_8_ts_004
Upright (0)
Reversed (0)
dataset_01@20241015_bs1_rota_10min_8_ts_002
Upright (222)
Reversed (98)
dataset_01@20241015_bs1_rota_10min_8_ts_006
Upright (1)
Reversed (0)
Detected membrane volume [µm³]
Membrane proximity (arbitrary units)
Spike conformation

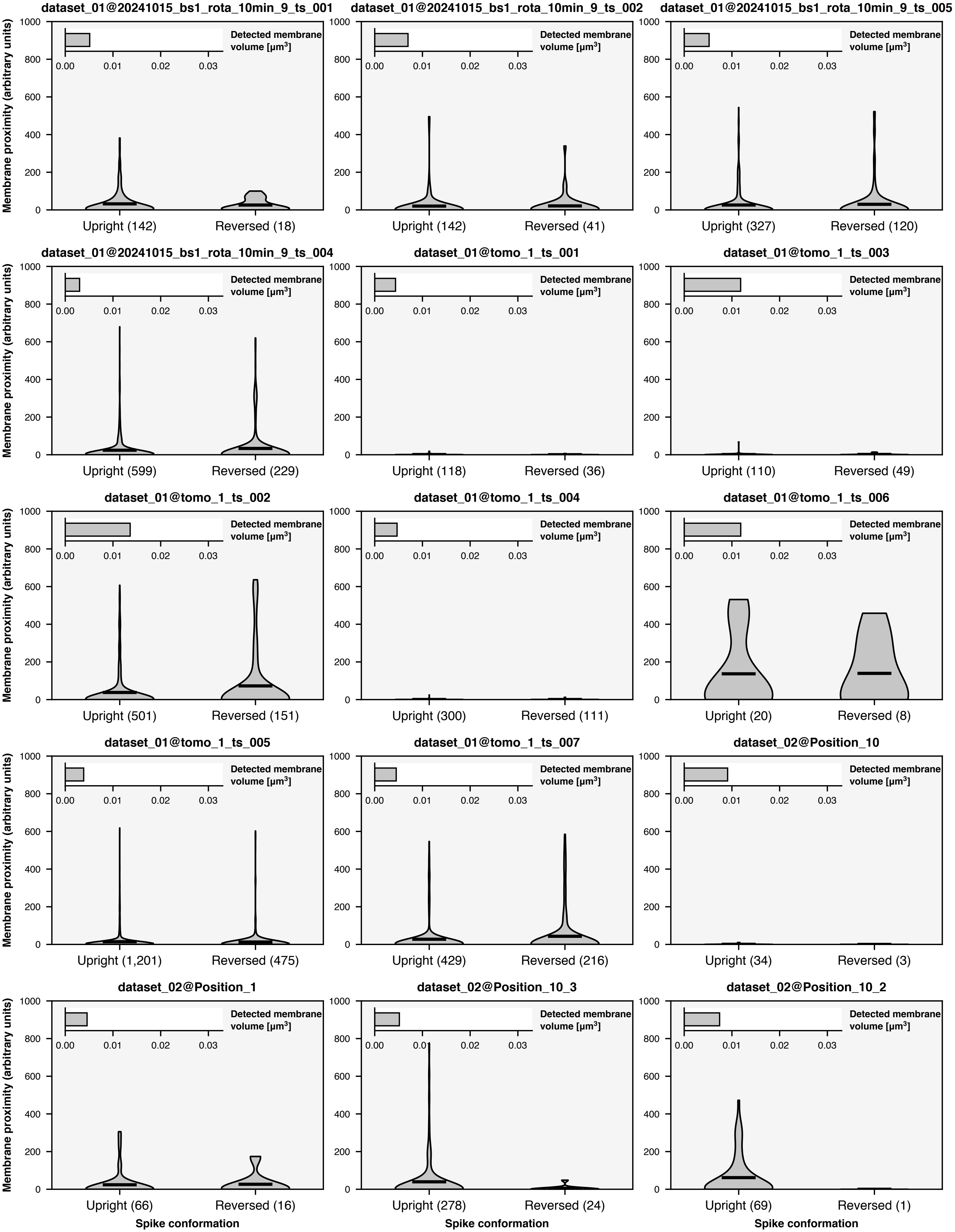
dataset_01@20241015_bs1_rota_10min_9_ts_001
Upright (142)
Reversed (18)
dataset_01@20241015_bs1_rota_10min_9_ts_002
Upright (142)
Reversed (41)
dataset_01@20241015_bs1_rota_10min_9_ts_005
Upright (327)
Reversed (120)
dataset_01@20241015_bs1_rota_10min_9_ts_004
Upright (599)
Reversed (229)
dataset_01@tomo_1_ts_001
Upright (118)
Reversed (36)
dataset_01@tomo_1_ts_003
Upright (110)
Reversed (49)
dataset_01@tomo_1_ts_002
Upright (501)
Reversed (151)
dataset_01@tomo_1_ts_004
Upright (300)
Reversed (111)
dataset_01@tomo_1_ts_006
Upright (20)
Reversed (8)
dataset_01@tomo_1_ts_005
Upright (1,201)
Reversed (475)
dataset_01@tomo_1_ts_007
Upright (429)
Reversed (216)
dataset_02@Position_10
Upright (34)
Reversed (3)
dataset_02@Position_1
Upright (66)
Reversed (16)
dataset_02@Position_10_3
Upright (278)
Reversed (24)
dataset_02@Position_10_2
Upright (69)
Reversed (1)
Membrane proximity (arbitrary units)
Detected membrane volume [µm³]
Spike conformation
0
200
400
600
800
1000
0.00
0.01
0.02
0.03

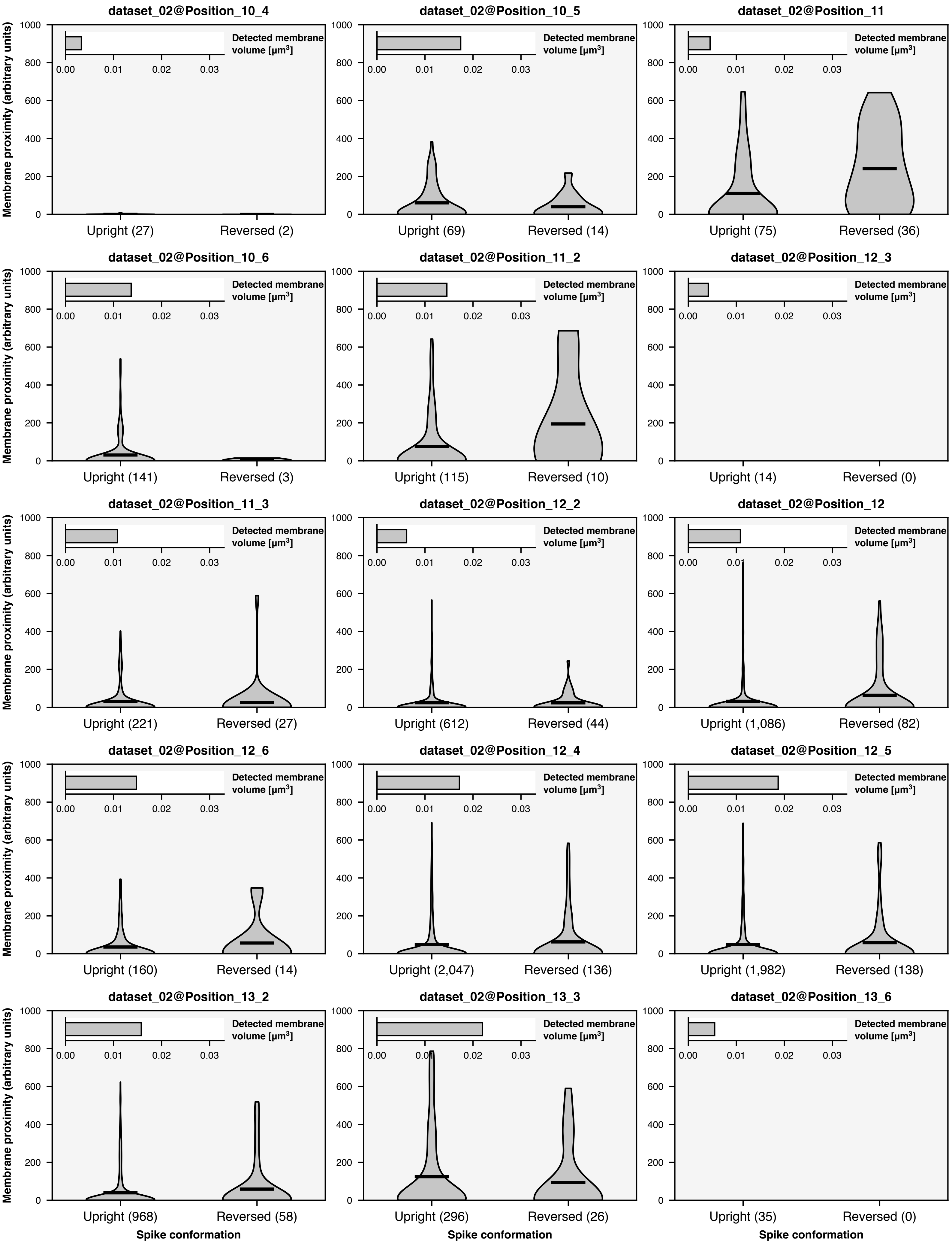
dataset_02@Position_10_4
Upright (27)
Reversed (2)
dataset_02@Position_10_5
Upright (69)
Reversed (14)
dataset_02@Position_11
Upright (75)
Reversed (36)
dataset_02@Position_10_6
Upright (141)
Reversed (3)
dataset_02@Position_11_2
Upright (115)
Reversed (10)
dataset_02@Position_12_3
Upright (14)
Reversed (0)
dataset_02@Position_11_3
Upright (221)
Reversed (27)
dataset_02@Position_12_2
Upright (612)
Reversed (44)
dataset_02@Position_12
Upright (1,086)
Reversed (82)
dataset_02@Position_12_6
Upright (160)
Reversed (14)
dataset_02@Position_12_4
Upright (2,047)
Reversed (136)
dataset_02@Position_12_5
Upright (1,982)
Reversed (138)
dataset_02@Position_13_2
Upright (968)
Reversed (58)
dataset_02@Position_13_3
Upright (296)
Reversed (26)
dataset_02@Position_13_6
Upright (35)
Reversed (0)
Detected membrane volume [μm³]
Membrane proximity (arbitrary units)
Spike conformation

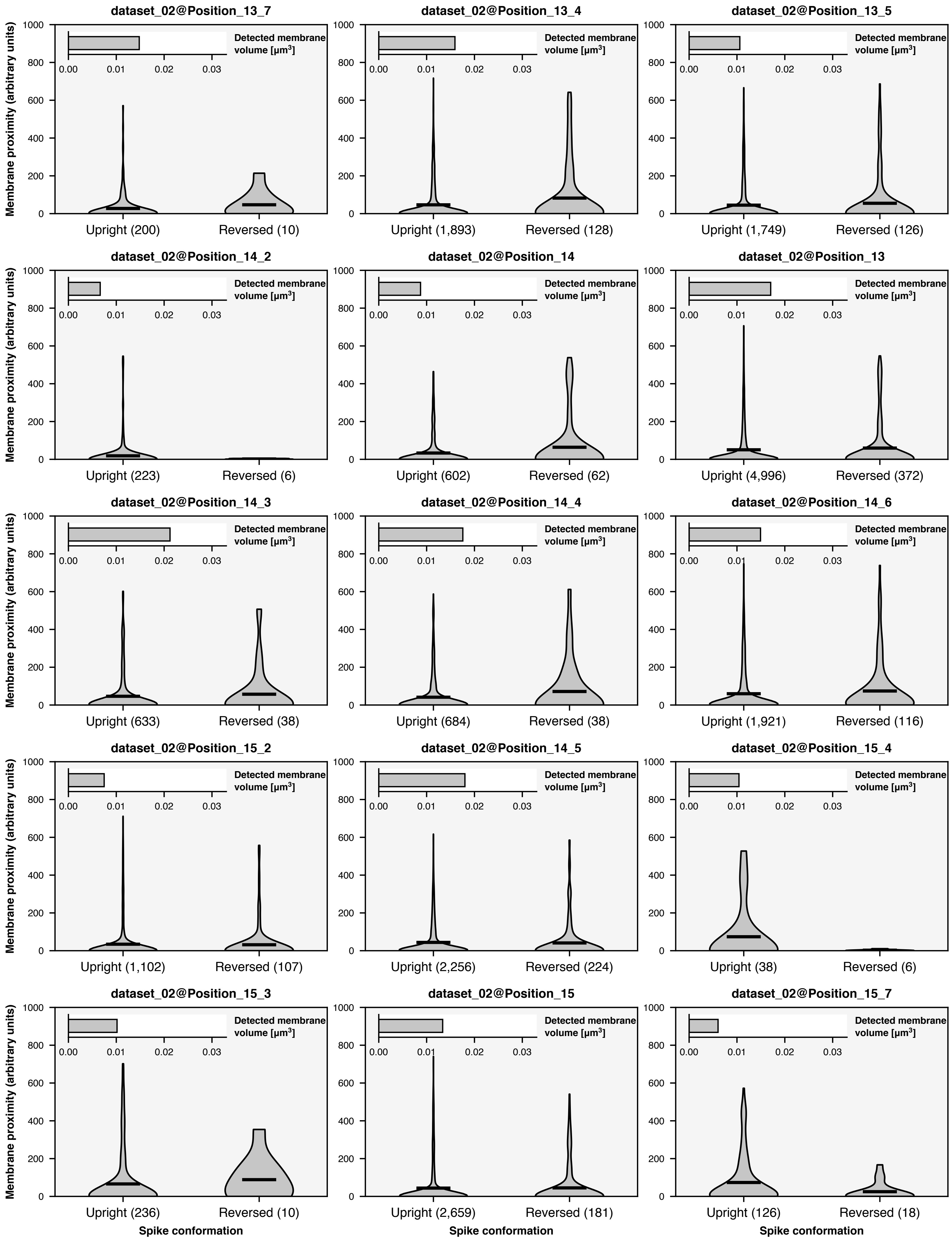

dataset_02@Position_13_7
Upright (200)
Reversed (10)
dataset_02@Position_13_4
Upright (1,893)
Reversed (128)
dataset_02@Position_13_5
Upright (1,749)
Reversed (126)
dataset_02@Position_14_2
Upright (223)
Reversed (6)
dataset_02@Position_14
Upright (602)
Reversed (62)
dataset_02@Position_13
Upright (4,996)
Reversed (372)
dataset_02@Position_14_3
Upright (633)
Reversed (38)
dataset_02@Position_14_4
Upright (684)
Reversed (38)
dataset_02@Position_14_6
Upright (1,921)
Reversed (116)
dataset_02@Position_15_2
Upright (1,102)
Reversed (107)
dataset_02@Position_14_5
Upright (2,256)
Reversed (224)
dataset_02@Position_15_4
Upright (38)
Reversed (6)
dataset_02@Position_15_3
Upright (236)
Reversed (10)
dataset_02@Position_15
Upright (2,659)
Reversed (181)
dataset_02@Position_15_7
Upright (126)
Reversed (18)
Detected membrane volume [μm³]
0.00
0.01
0.02
0.03
Membrane proximity (arbitrary units)
0
200
400
600
800
1000
Spike conformation

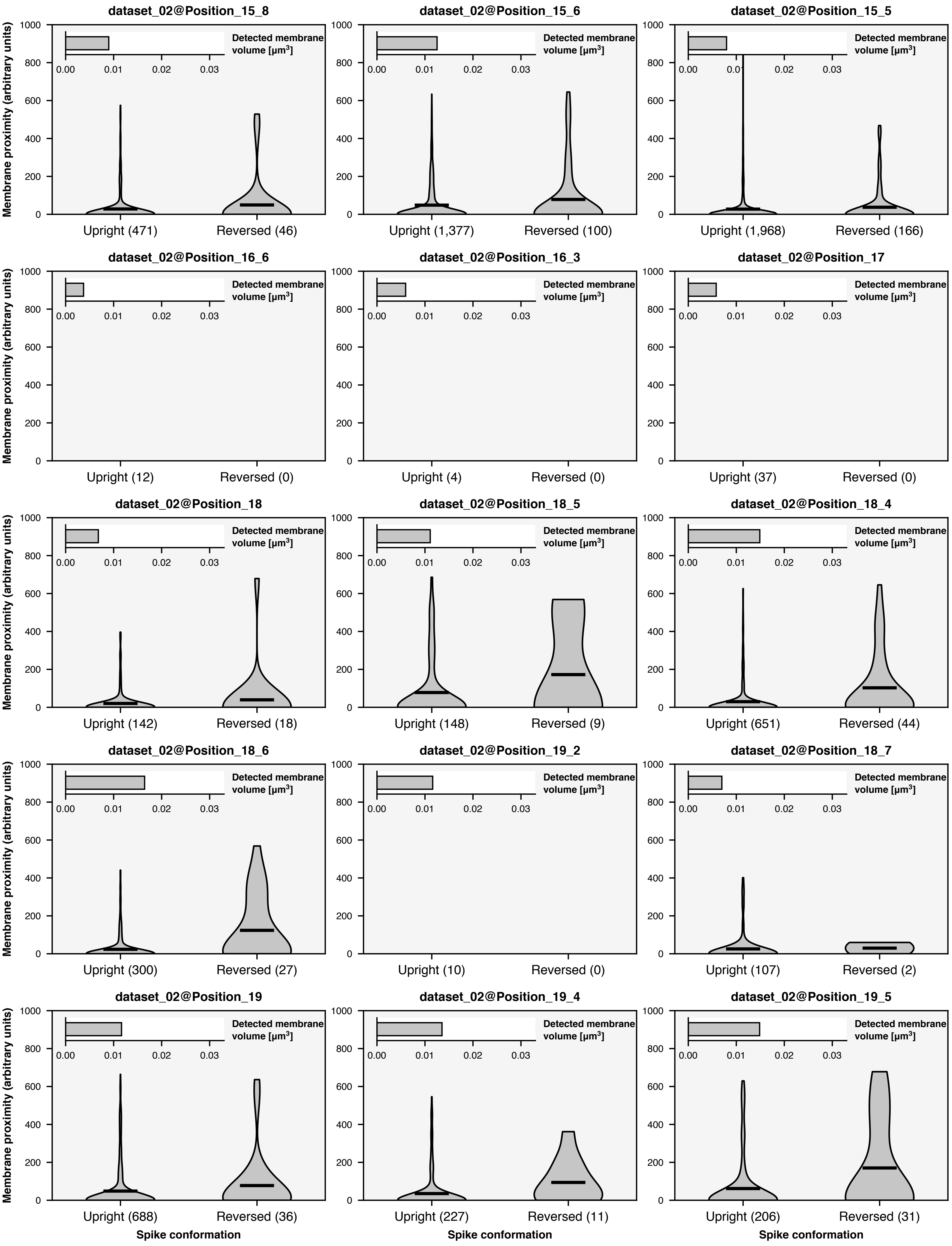
dataset_02@Position_15_8
dataset_02@Position_15_6
dataset_02@Position_15_5
dataset_02@Position_16_6
dataset_02@Position_16_3
dataset_02@Position_17
dataset_02@Position_18
dataset_02@Position_18_5
dataset_02@Position_18_4
dataset_02@Position_18_6
dataset_02@Position_19_2
dataset_02@Position_18_7
dataset_02@Position_19
dataset_02@Position_19_4
dataset_02@Position_19_5
Membrane proximity (arbitrary units)
Detected membrane volume [µm³]
Upright (471)
Reversed (46)
Upright (1,377)
Reversed (100)
Upright (1,968)
Reversed (166)
Upright (12)
Reversed (0)
Upright (4)
Reversed (0)
Upright (37)
Reversed (0)
Upright (142)
Reversed (18)
Upright (148)
Reversed (9)
Upright (651)
Reversed (44)
Upright (300)
Reversed (27)
Upright (10)
Reversed (0)
Upright (107)
Reversed (2)
Upright (688)
Reversed (36)
Upright (227)
Reversed (11)
Upright (206)
Reversed (31)
Spike conformation

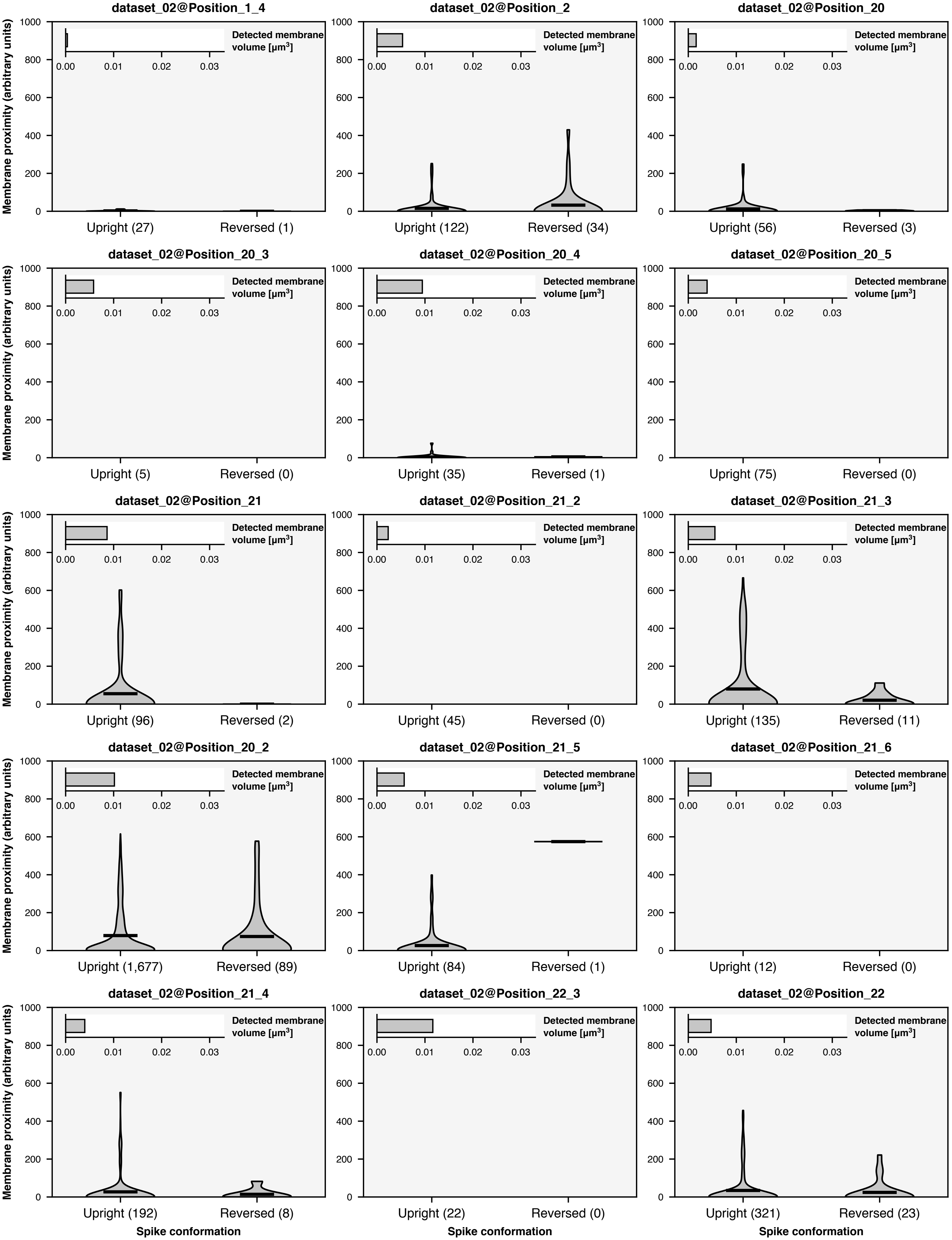
dataset_02@Position_1_4
dataset_02@Position_2
dataset_02@Position_20
dataset_02@Position_20_3
dataset_02@Position_20_4
dataset_02@Position_20_5
dataset_02@Position_21
dataset_02@Position_21_2
dataset_02@Position_21_3
dataset_02@Position_20_2
dataset_02@Position_21_5
dataset_02@Position_21_6
dataset_02@Position_21_4
dataset_02@Position_22_3
dataset_02@Position_22
Detected membrane volume [μm³]
Membrane proximity (arbitrary units)
Upright (27)
Reversed (1)
Upright (122)
Reversed (34)
Upright (56)
Reversed (3)
Upright (5)
Reversed (0)
Upright (35)
Reversed (1)
Upright (75)
Reversed (0)
Upright (96)
Reversed (2)
Upright (45)
Reversed (0)
Upright (135)
Reversed (11)
Upright (1,677)
Reversed (89)
Upright (84)
Reversed (1)
Upright (12)
Reversed (0)
Upright (192)
Reversed (8)
Upright (22)
Reversed (0)
Upright (321)
Reversed (23)
Spike conformation

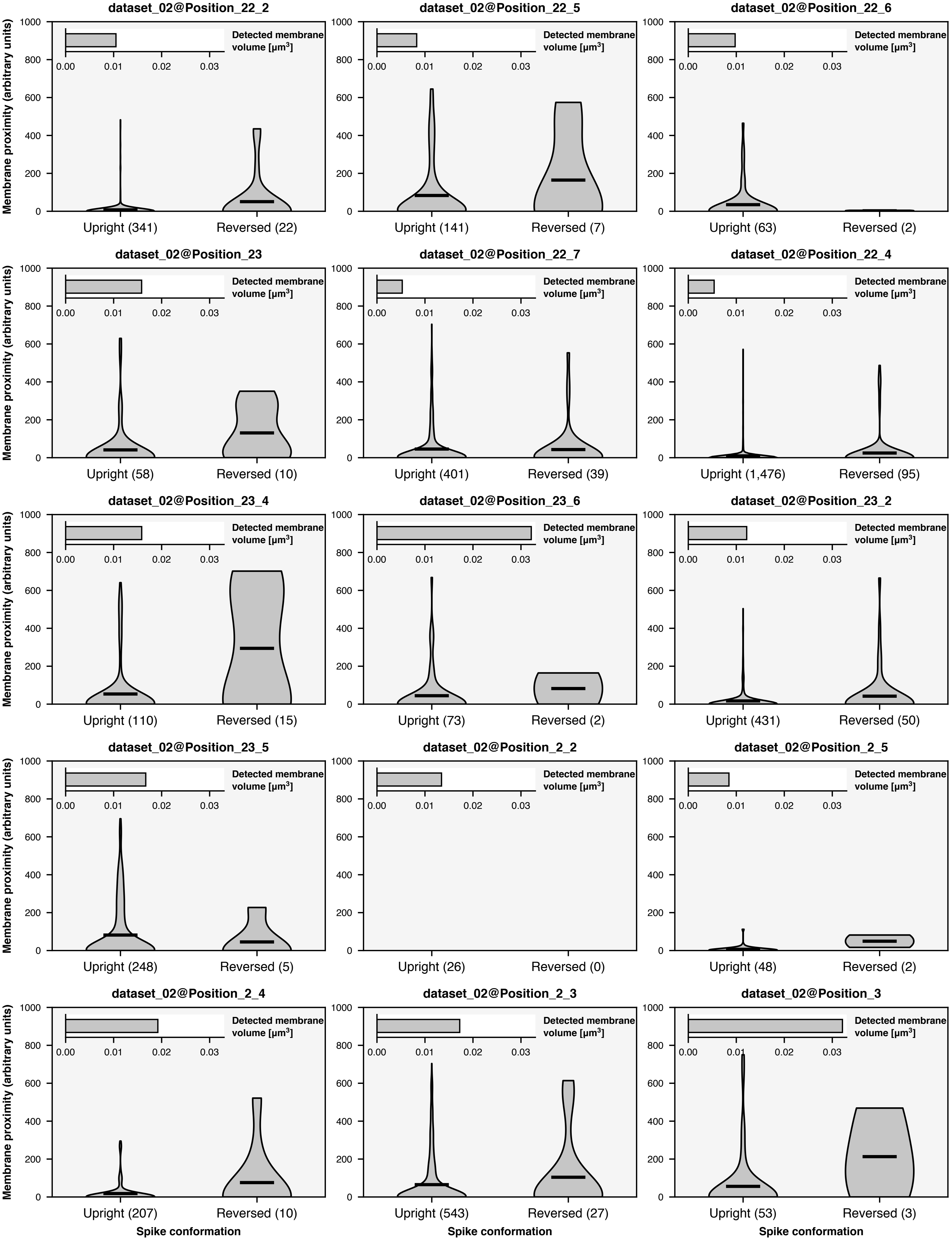
dataset_02@Position_22_2
dataset_02@Position_22_5
dataset_02@Position_22_6
dataset_02@Position_23
dataset_02@Position_22_7
dataset_02@Position_22_4
dataset_02@Position_23_4
dataset_02@Position_23_6
dataset_02@Position_23_2
dataset_02@Position_23_5
dataset_02@Position_2_2
dataset_02@Position_2_5
dataset_02@Position_2_4
dataset_02@Position_2_3
dataset_02@Position_3
Membrane proximity (arbitrary units)
Detected membrane volume [μm³]
Upright (341)
Reversed (22)
Upright (141)
Reversed (7)
Upright (63)
Reversed (2)
Upright (58)
Reversed (10)
Upright (401)
Reversed (39)
Upright (1,476)
Reversed (95)
Upright (110)
Reversed (15)
Upright (73)
Reversed (2)
Upright (431)
Reversed (50)
Upright (248)
Reversed (5)
Upright (26)
Reversed (0)
Upright (48)
Reversed (2)
Upright (207)
Reversed (10)
Upright (543)
Reversed (27)
Upright (53)
Reversed (3)
Spike conformation

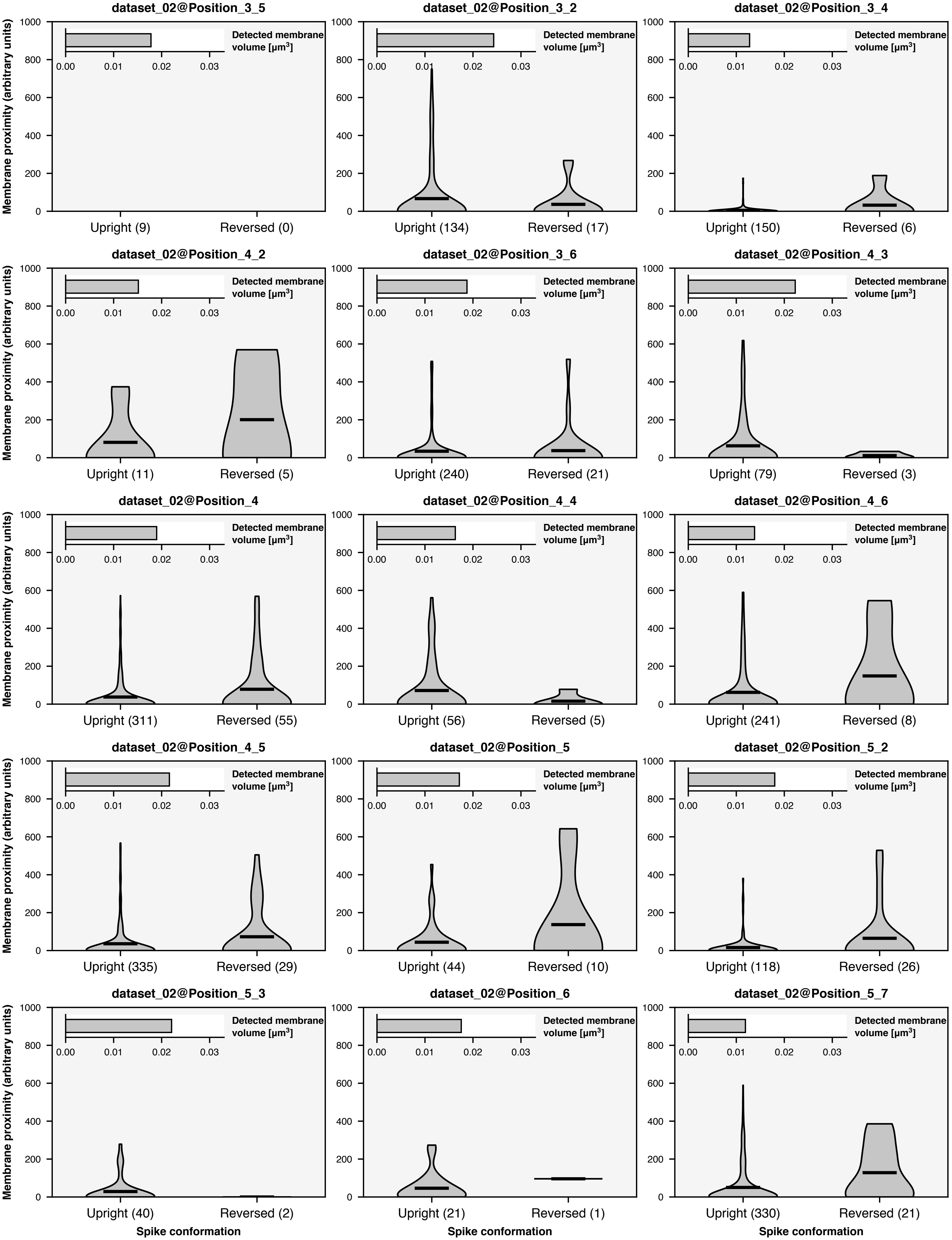
dataset_02@Position_3_5
dataset_02@Position_3_2
dataset_02@Position_3_4
dataset_02@Position_4_2
dataset_02@Position_3_6
dataset_02@Position_4_3
dataset_02@Position_4
dataset_02@Position_4_4
dataset_02@Position_4_6
dataset_02@Position_4_5
dataset_02@Position_5
dataset_02@Position_5_2
dataset_02@Position_5_3
dataset_02@Position_6
dataset_02@Position_5_7
Detected membrane volume [μm³]
Membrane proximity (arbitrary units)
Upright (9)
Reversed (0)
Upright (134)
Reversed (17)
Upright (150)
Reversed (6)
Upright (11)
Reversed (5)
Upright (240)
Reversed (21)
Upright (79)
Reversed (3)
Upright (311)
Reversed (55)
Upright (56)
Reversed (5)
Upright (241)
Reversed (8)
Upright (335)
Reversed (29)
Upright (44)
Reversed (10)
Upright (118)
Reversed (26)
Upright (40)
Reversed (2)
Upright (21)
Reversed (1)
Upright (330)
Reversed (21)
Spike conformation

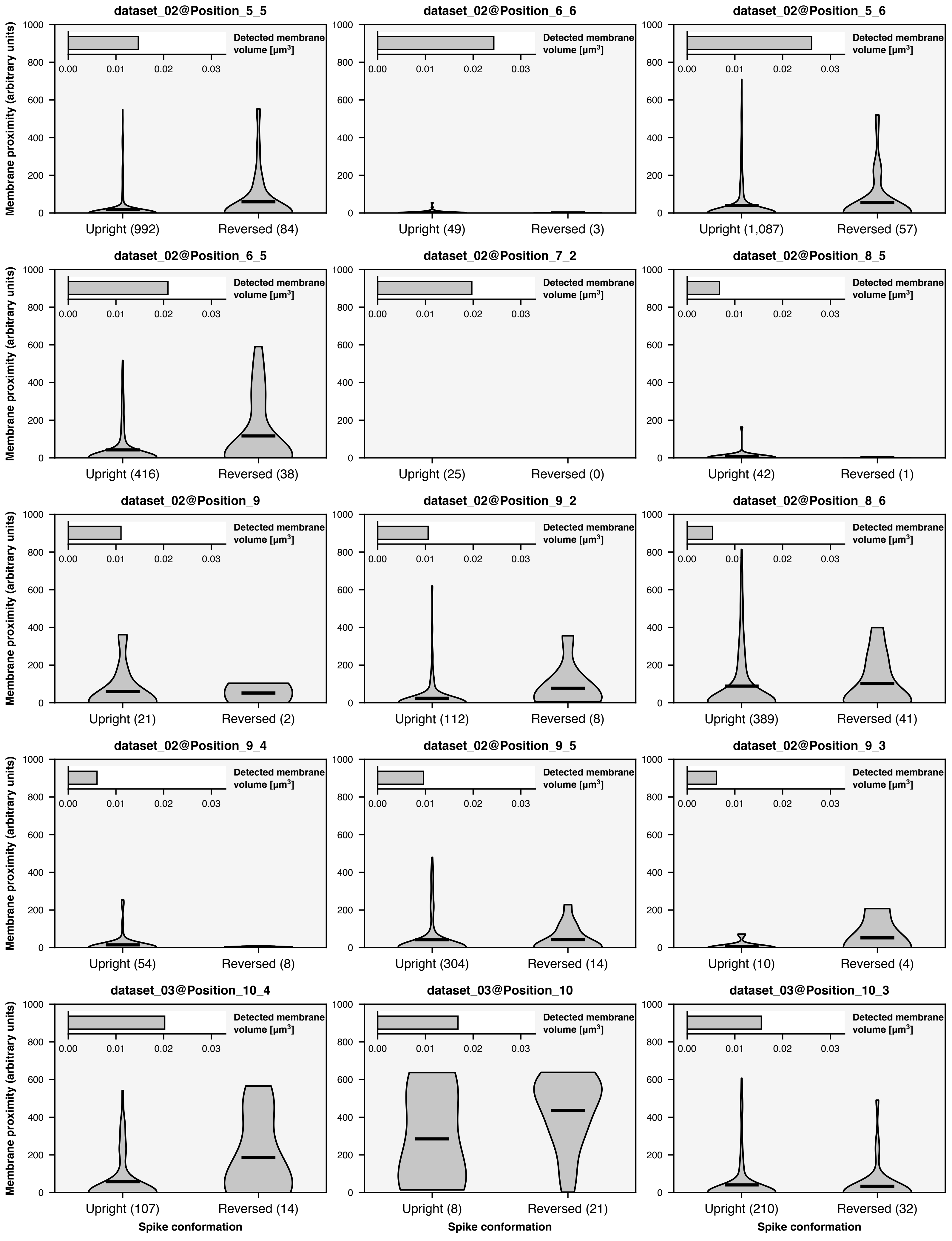

dataset_02@Position_5_5
dataset_02@Position_6_6
dataset_02@Position_5_6
dataset_02@Position_6_5
dataset_02@Position_7_2
dataset_02@Position_8_5
dataset_02@Position_9
dataset_02@Position_9_2
dataset_02@Position_8_6
dataset_02@Position_9_4
dataset_02@Position_9_5
dataset_02@Position_9_3
dataset_03@Position_10_4
dataset_03@Position_10
dataset_03@Position_10_3
Detected membrane volume [μm³]
Membrane proximity (arbitrary units)
Upright (992)
Reversed (84)
Upright (49)
Reversed (3)
Upright (1,087)
Reversed (57)
Upright (416)
Reversed (38)
Upright (25)
Reversed (0)
Upright (42)
Reversed (1)
Upright (21)
Reversed (2)
Upright (112)
Reversed (8)
Upright (389)
Reversed (41)
Upright (54)
Reversed (8)
Upright (304)
Reversed (14)
Upright (10)
Reversed (4)
Upright (107)
Reversed (14)
Upright (8)
Reversed (21)
Upright (210)
Reversed (32)
Spike conformation

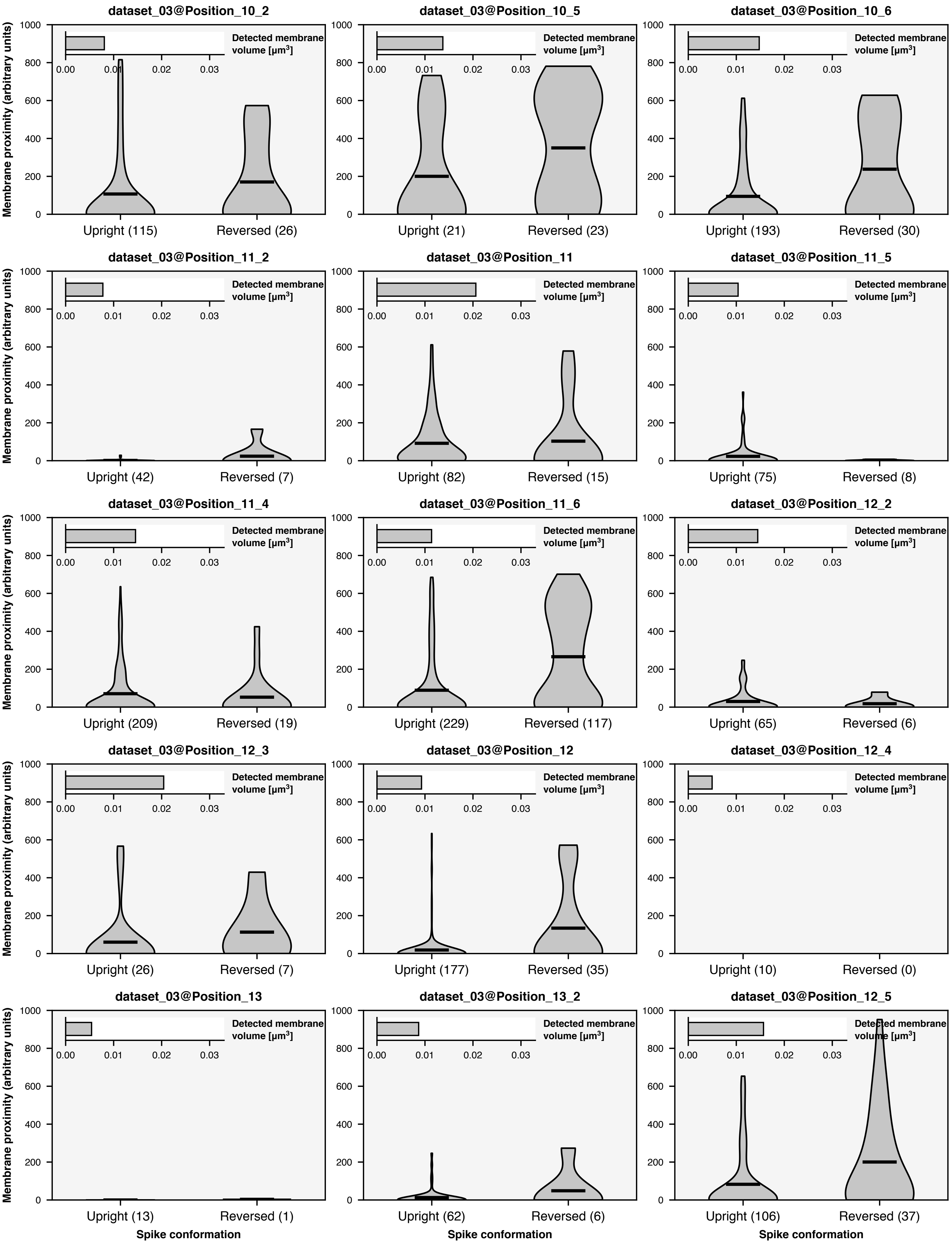
dataset_03@Position_10_2
dataset_03@Position_10_5
dataset_03@Position_10_6
dataset_03@Position_11_2
dataset_03@Position_11
dataset_03@Position_11_5
dataset_03@Position_11_4
dataset_03@Position_11_6
dataset_03@Position_12_2
dataset_03@Position_12_3
dataset_03@Position_12
dataset_03@Position_12_4
dataset_03@Position_13
dataset_03@Position_13_2
dataset_03@Position_12_5
Membrane proximity (arbitrary units)
Detected membrane volume [µm³]
Upright (115)
Reversed (26)
Upright (21)
Reversed (23)
Upright (193)
Reversed (30)
Upright (42)
Reversed (7)
Upright (82)
Reversed (15)
Upright (75)
Reversed (8)
Upright (209)
Reversed (19)
Upright (229)
Reversed (117)
Upright (65)
Reversed (6)
Upright (26)
Reversed (7)
Upright (177)
Reversed (35)
Upright (10)
Reversed (0)
Upright (13)
Reversed (1)
Upright (62)
Reversed (6)
Upright (106)
Reversed (37)
Spike conformation

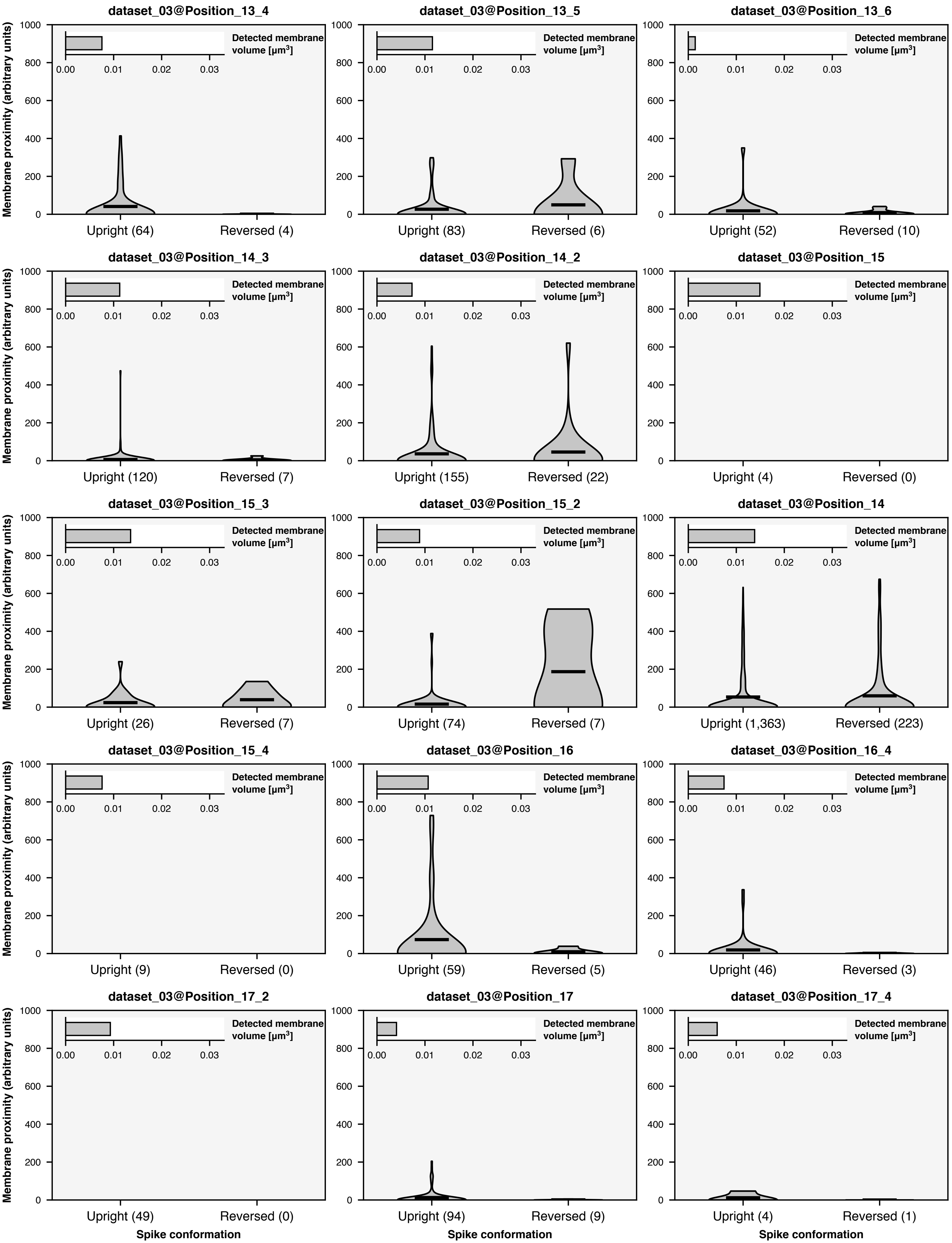

dataset_03@Position_13_4
dataset_03@Position_13_5
dataset_03@Position_13_6
dataset_03@Position_14_3
dataset_03@Position_14_2
dataset_03@Position_15
dataset_03@Position_15_3
dataset_03@Position_15_2
dataset_03@Position_14
dataset_03@Position_15_4
dataset_03@Position_16
dataset_03@Position_16_4
dataset_03@Position_17_2
dataset_03@Position_17
dataset_03@Position_17_4
Membrane proximity (arbitrary units)
Detected membrane volume [μm³]
Upright (64) Reversed (4)
Upright (83) Reversed (6)
Upright (52) Reversed (10)
Upright (120) Reversed (7)
Upright (155) Reversed (22)
Upright (4) Reversed (0)
Upright (26) Reversed (7)
Upright (74) Reversed (7)
Upright (1,363) Reversed (223)
Upright (9) Reversed (0)
Upright (59) Reversed (5)
Upright (46) Reversed (3)
Upright (49) Reversed (0)
Upright (94) Reversed (9)
Upright (4) Reversed (1)
Spike conformation

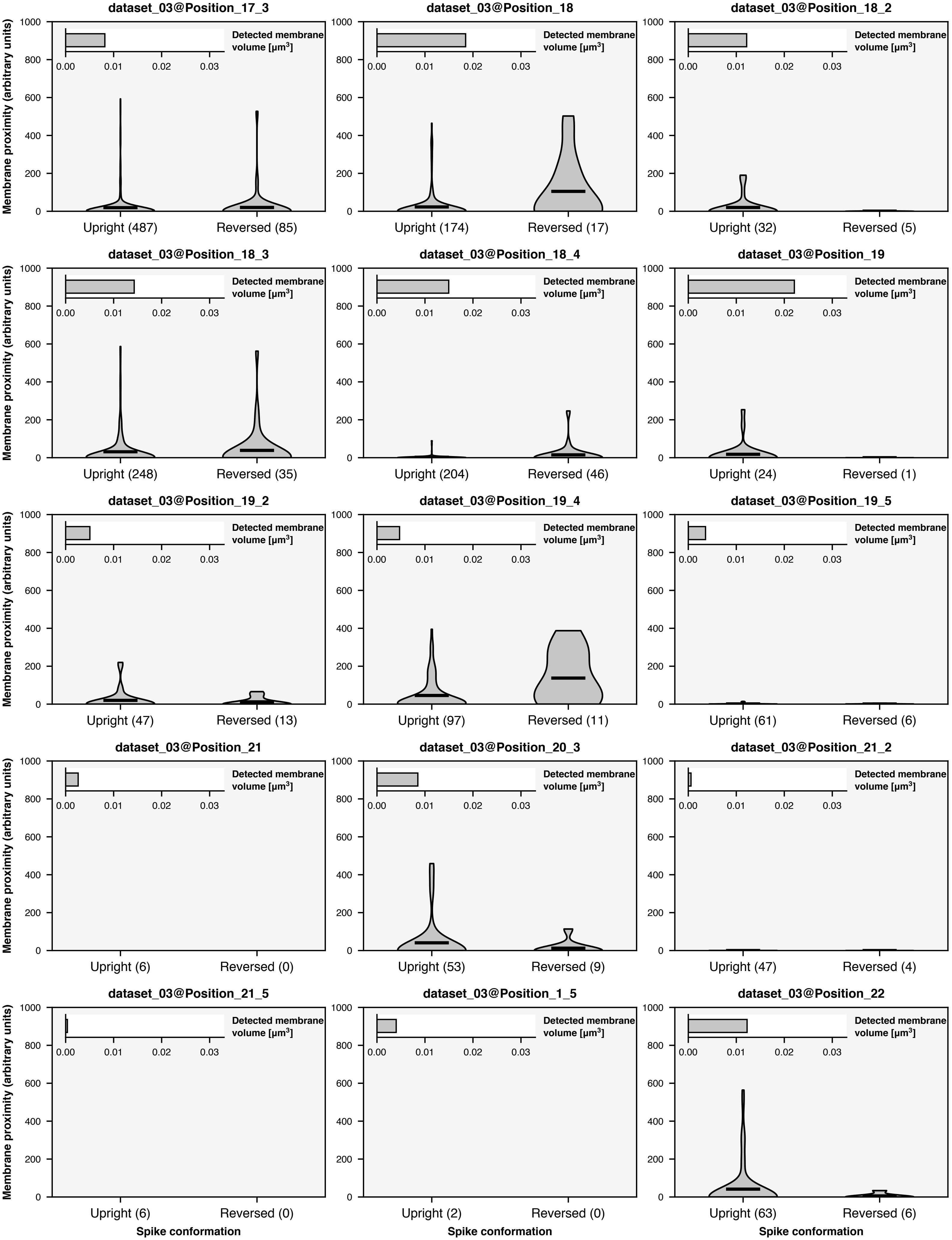
dataset_03@Position_17_3
dataset_03@Position_18
dataset_03@Position_18_2
dataset_03@Position_18_3
dataset_03@Position_18_4
dataset_03@Position_19
dataset_03@Position_19_2
dataset_03@Position_19_4
dataset_03@Position_19_5
dataset_03@Position_21
dataset_03@Position_20_3
dataset_03@Position_21_2
dataset_03@Position_21_5
dataset_03@Position_1_5
dataset_03@Position_22
Detected membrane volume [μm$^3$]
0.00 0.01 0.02 0.03
Membrane proximity (arbitrary units)
0 200 400 600 800 1000
Upright (487) Reversed (85)
Upright (174) Reversed (17)
Upright (32) Reversed (5)
Upright (248) Reversed (35)
Upright (204) Reversed (46)
Upright (24) Reversed (1)
Upright (47) Reversed (13)
Upright (97) Reversed (11)
Upright (61) Reversed (6)
Upright (6) Reversed (0)
Upright (53) Reversed (9)
Upright (47) Reversed (4)
Upright (6) Reversed (0)
Upright (2) Reversed (0)
Upright (63) Reversed (6)
Spike conformation

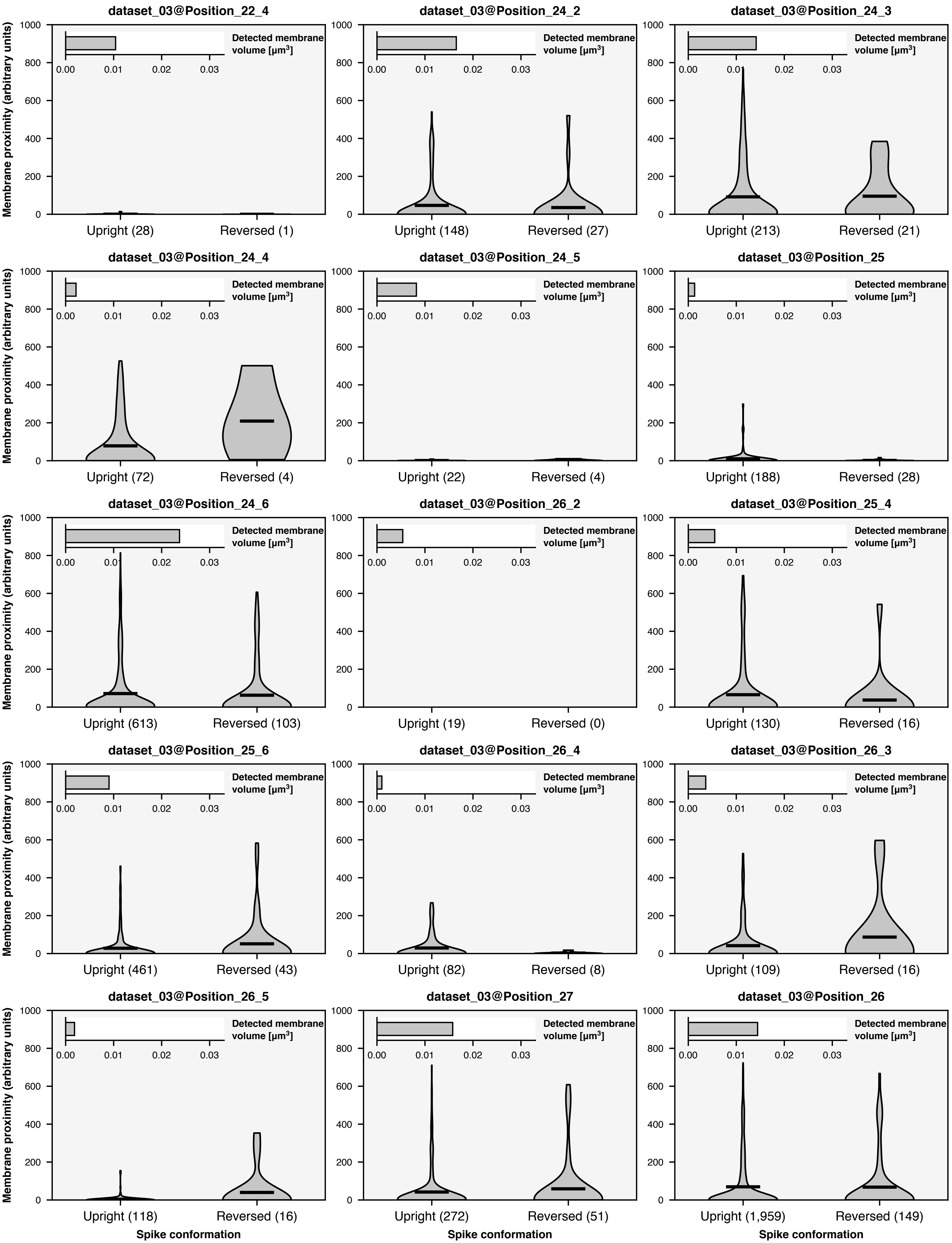
dataset_03@Position_22_4
dataset_03@Position_24_2
dataset_03@Position_24_3
dataset_03@Position_24_4
dataset_03@Position_24_5
dataset_03@Position_25
dataset_03@Position_24_6
dataset_03@Position_26_2
dataset_03@Position_25_4
dataset_03@Position_25_6
dataset_03@Position_26_4
dataset_03@Position_26_3
dataset_03@Position_26_5
dataset_03@Position_27
dataset_03@Position_26
Membrane proximity (arbitrary units)
Detected membrane volume [µm$^3$]
Upright (28)
Reversed (1)
Upright (148)
Reversed (27)
Upright (213)
Reversed (21)
Upright (72)
Reversed (4)
Upright (22)
Reversed (4)
Upright (188)
Reversed (28)
Upright (613)
Reversed (103)
Upright (19)
Reversed (0)
Upright (130)
Reversed (16)
Upright (461)
Reversed (43)
Upright (82)
Reversed (8)
Upright (109)
Reversed (16)
Upright (118)
Reversed (16)
Upright (272)
Reversed (51)
Upright (1,959)
Reversed (149)
Spike conformation

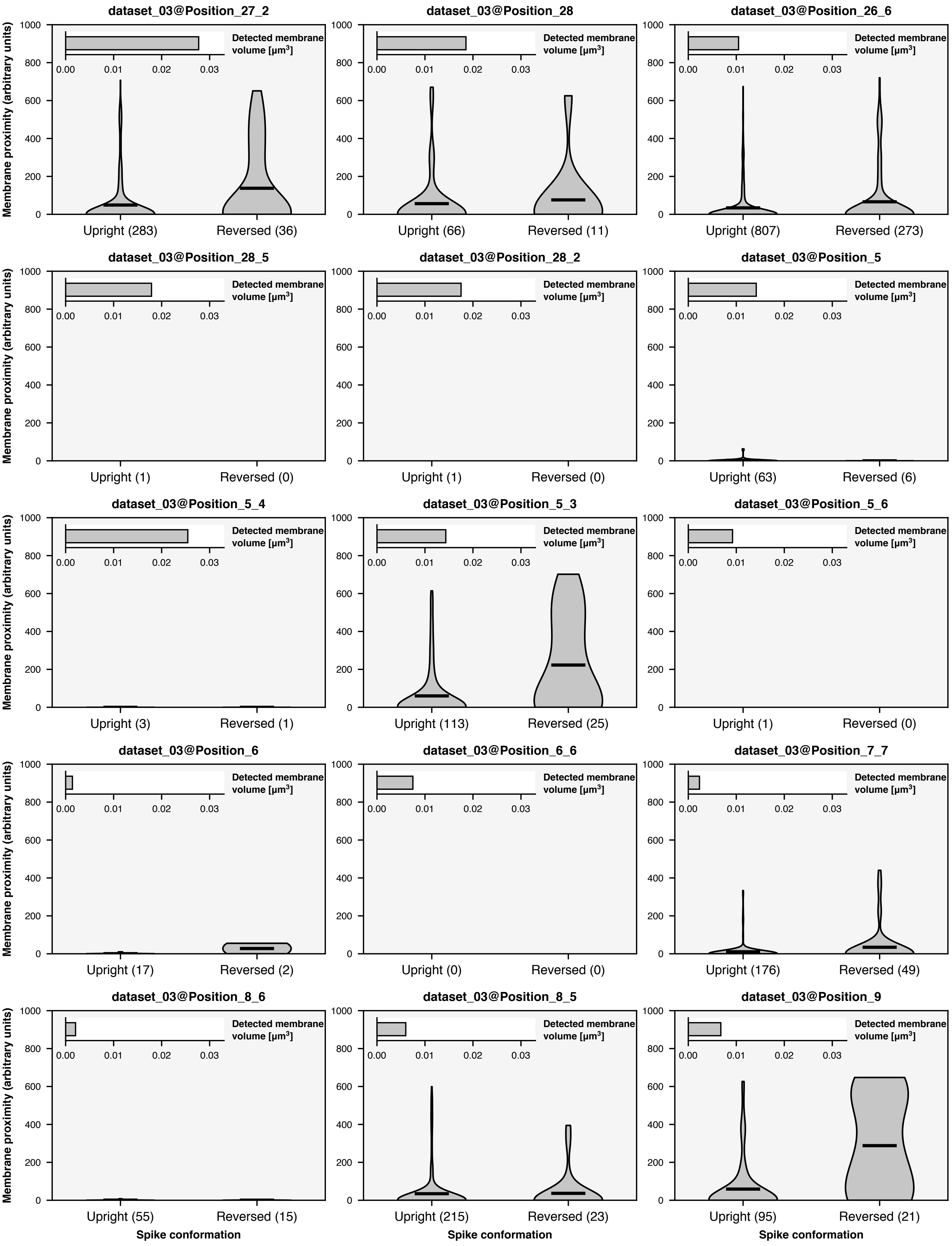
dataset_03@Position_27_2
Membrane proximity (arbitrary units)
Detected membrane volume [μm³]
0.00 0.01 0.02 0.03
0 200 400 600 800 1000
Upright (283)
Reversed (36)
dataset_03@Position_28
Upright (66)
Reversed (11)
dataset_03@Position_26_6
Upright (807)
Reversed (273)
dataset_03@Position_28_5
Upright (1)
Reversed (0)
dataset_03@Position_28_2
Upright (1)
Reversed (0)
dataset_03@Position_5
Upright (63)
Reversed (6)
dataset_03@Position_5_4
Upright (3)
Reversed (1)
dataset_03@Position_5_3
Upright (113)
Reversed (25)
dataset_03@Position_5_6
Upright (1)
Reversed (0)
dataset_03@Position_6
Upright (17)
Reversed (2)
dataset_03@Position_6_6
Upright (0)
Reversed (0)
dataset_03@Position_7_7
Upright (176)
Reversed (49)
dataset_03@Position_8_6
Upright (55)
Reversed (15)
dataset_03@Position_8_5
Upright (215)
Reversed (23)
dataset_03@Position_9
Upright (95)
Reversed (21)
Spike conformation

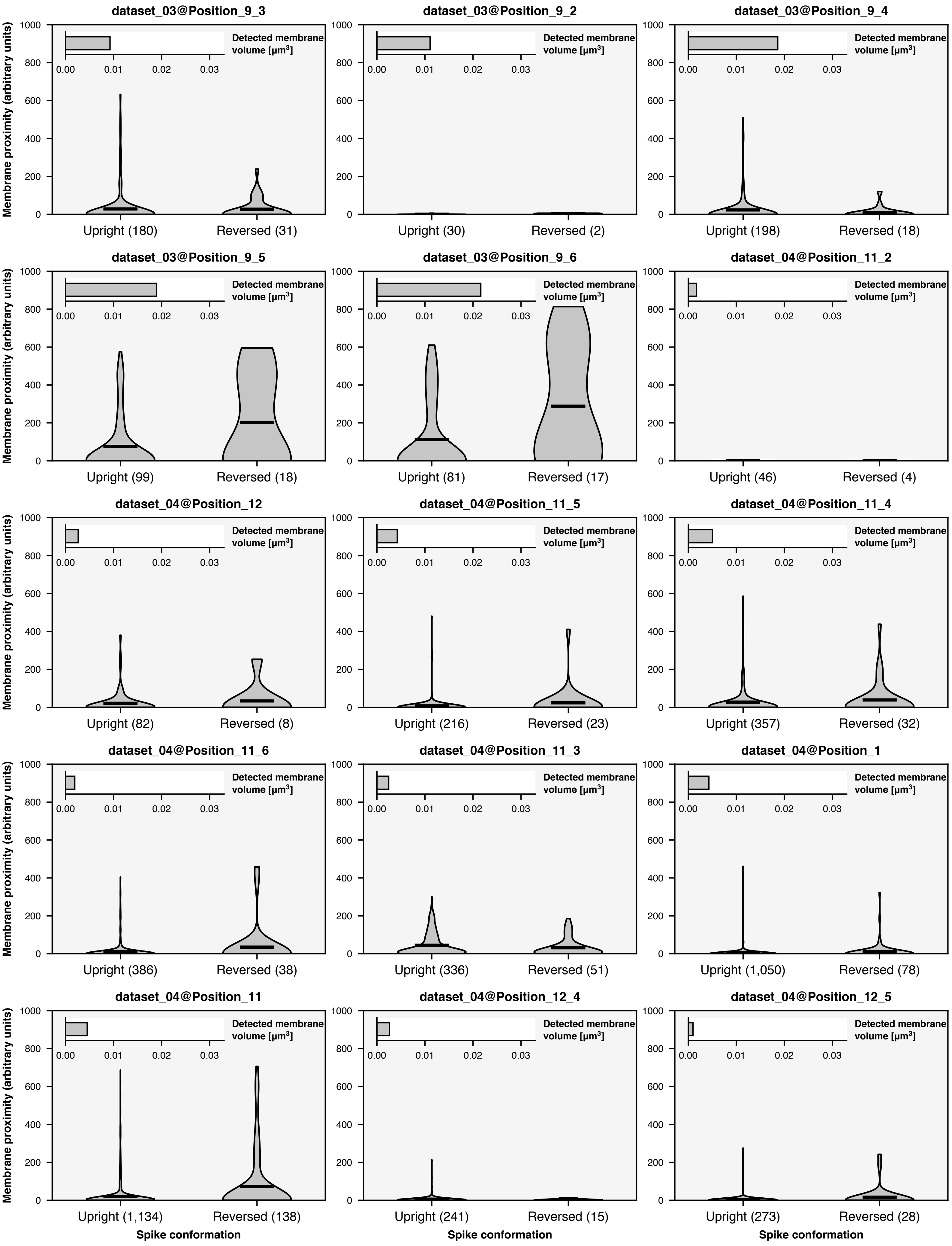
dataset_03@Position_9_3
Detected membrane volume [µm³]
Upright (180)
Reversed (31)
dataset_03@Position_9_2
Upright (30)
Reversed (2)
dataset_03@Position_9_4
Upright (198)
Reversed (18)
dataset_03@Position_9_5
Upright (99)
Reversed (18)
dataset_03@Position_9_6
Upright (81)
Reversed (17)
dataset_04@Position_11_2
Upright (46)
Reversed (4)
dataset_04@Position_12
Upright (82)
Reversed (8)
dataset_04@Position_11_5
Upright (216)
Reversed (23)
dataset_04@Position_11_4
Upright (357)
Reversed (32)
dataset_04@Position_11_6
Upright (386)
Reversed (38)
dataset_04@Position_11_3
Upright (336)
Reversed (51)
dataset_04@Position_1
Upright (1,050)
Reversed (78)
dataset_04@Position_11
Upright (1,134)
Reversed (138)
dataset_04@Position_12_4
Upright (241)
Reversed (15)
dataset_04@Position_12_5
Upright (273)
Reversed (28)
Membrane proximity (arbitrary units)
Spike conformation
0
200
400
600
800
1000
0.00
0.01
0.02
0.03

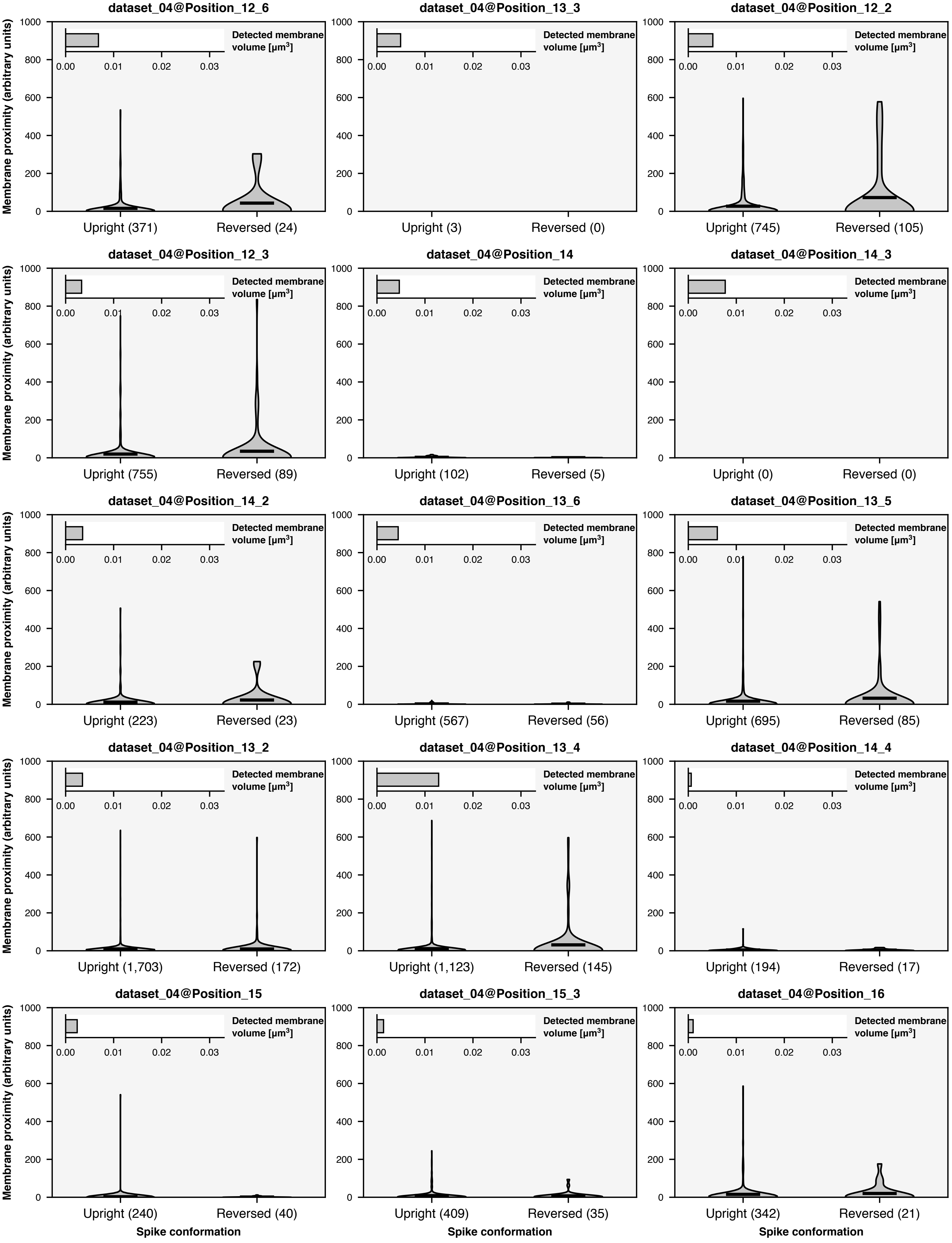
dataset_04@Position_12_6
Upright (371)
Reversed (24)
dataset_04@Position_13_3
Upright (3)
Reversed (0)
dataset_04@Position_12_2
Upright (745)
Reversed (105)
dataset_04@Position_12_3
Upright (755)
Reversed (89)
dataset_04@Position_14
Upright (102)
Reversed (5)
dataset_04@Position_14_3
Upright (0)
Reversed (0)
dataset_04@Position_14_2
Upright (223)
Reversed (23)
dataset_04@Position_13_6
Upright (567)
Reversed (56)
dataset_04@Position_13_5
Upright (695)
Reversed (85)
dataset_04@Position_13_2
Upright (1,703)
Reversed (172)
dataset_04@Position_13_4
Upright (1,123)
Reversed (145)
dataset_04@Position_14_4
Upright (194)
Reversed (17)
dataset_04@Position_15
Upright (240)
Reversed (40)
dataset_04@Position_15_3
Upright (409)
Reversed (35)
dataset_04@Position_16
Upright (342)
Reversed (21)
Membrane proximity (arbitrary units)
Detected membrane volume [μm³]
0.00
0.01
0.02
0.03
0
200
400
600
800
1000
Spike conformation

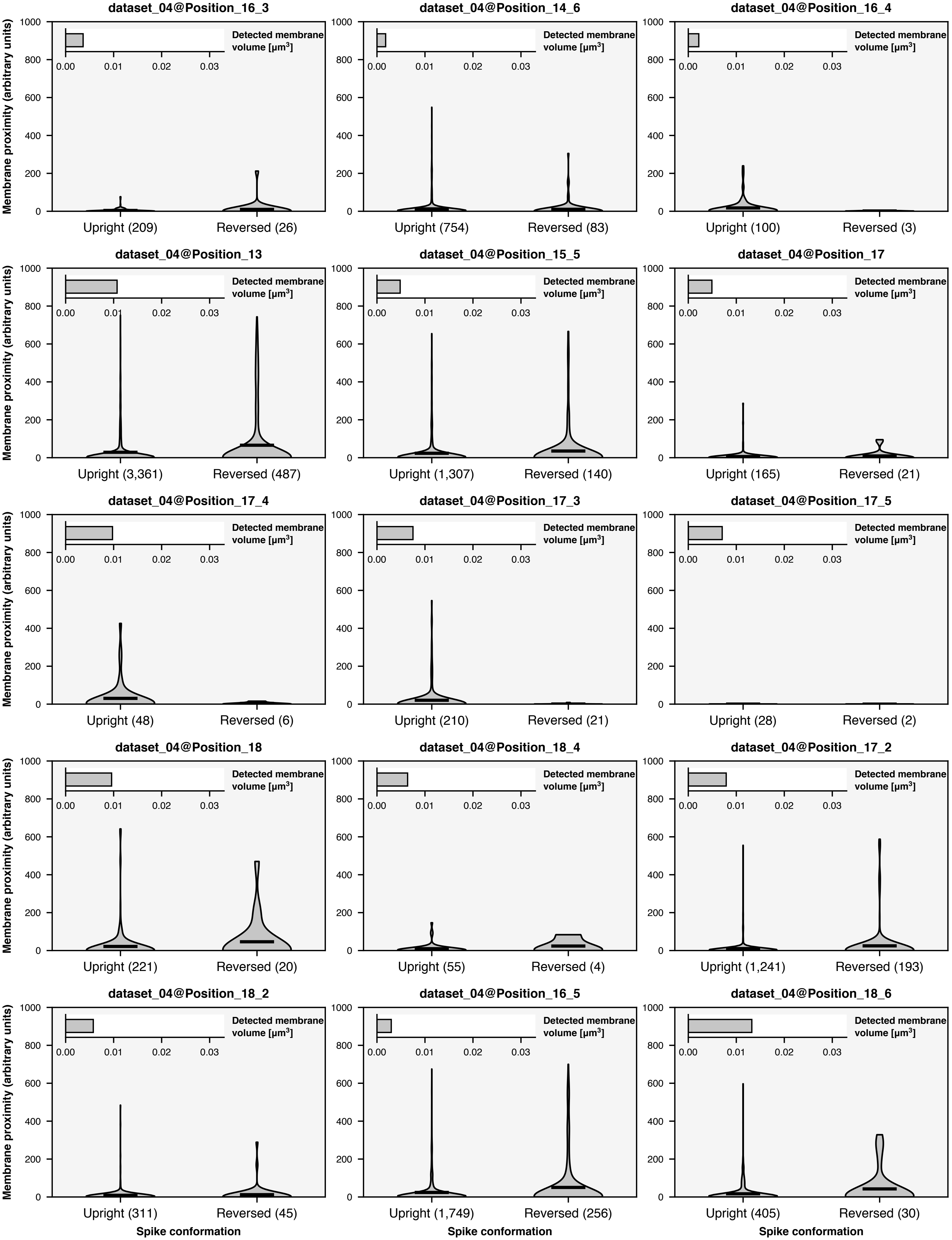

dataset_04@Position_16_3
dataset_04@Position_14_6
dataset_04@Position_16_4
dataset_04@Position_13
dataset_04@Position_15_5
dataset_04@Position_17
dataset_04@Position_17_4
dataset_04@Position_17_3
dataset_04@Position_17_5
dataset_04@Position_18
dataset_04@Position_18_4
dataset_04@Position_17_2
dataset_04@Position_18_2
dataset_04@Position_16_5
dataset_04@Position_18_6
Membrane proximity (arbitrary units)
Detected membrane volume [μm³]
0.00 0.01 0.02 0.03
0 200 400 600 800 1000
Upright (209) Reversed (26)
Upright (754) Reversed (83)
Upright (100) Reversed (3)
Upright (3,361) Reversed (487)
Upright (1,307) Reversed (140)
Upright (165) Reversed (21)
Upright (48) Reversed (6)
Upright (210) Reversed (21)
Upright (28) Reversed (2)
Upright (221) Reversed (20)
Upright (55) Reversed (4)
Upright (1,241) Reversed (193)
Upright (311) Reversed (45)
Upright (1,749) Reversed (256)
Upright (405) Reversed (30)
Spike conformation

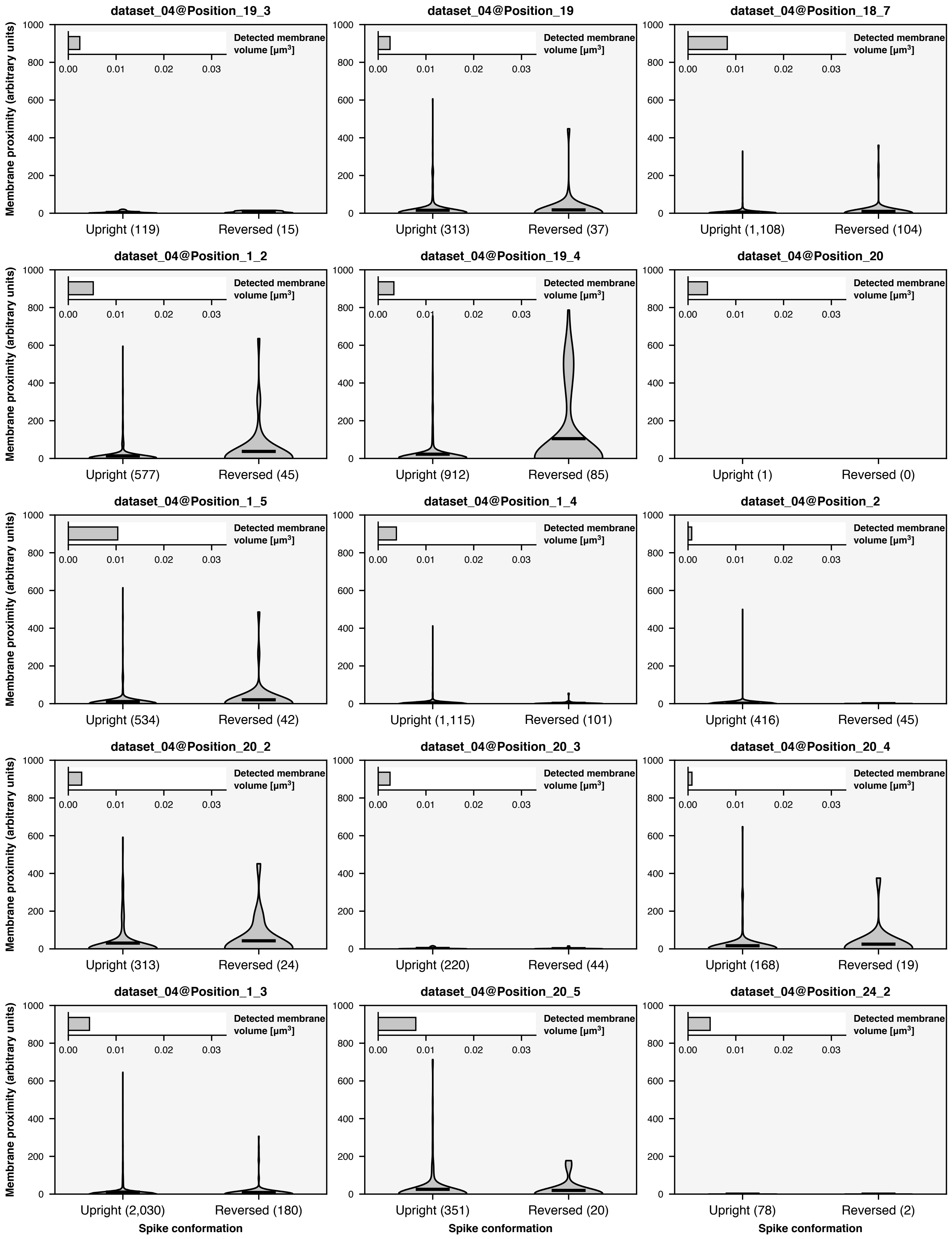
dataset_04@Position_19_3
dataset_04@Position_19
dataset_04@Position_18_7
dataset_04@Position_1_2
dataset_04@Position_19_4
dataset_04@Position_20
dataset_04@Position_1_5
dataset_04@Position_1_4
dataset_04@Position_2
dataset_04@Position_20_2
dataset_04@Position_20_3
dataset_04@Position_20_4
dataset_04@Position_1_3
dataset_04@Position_20_5
dataset_04@Position_24_2
Detected membrane volume [μm³]
Membrane proximity (arbitrary units)
Upright (119)
Reversed (15)
Upright (313)
Reversed (37)
Upright (1,108)
Reversed (104)
Upright (577)
Reversed (45)
Upright (912)
Reversed (85)
Upright (1)
Reversed (0)
Upright (534)
Reversed (42)
Upright (1,115)
Reversed (101)
Upright (416)
Reversed (45)
Upright (313)
Reversed (24)
Upright (220)
Reversed (44)
Upright (168)
Reversed (19)
Upright (2,030)
Reversed (180)
Upright (351)
Reversed (20)
Upright (78)
Reversed (2)
Spike conformation

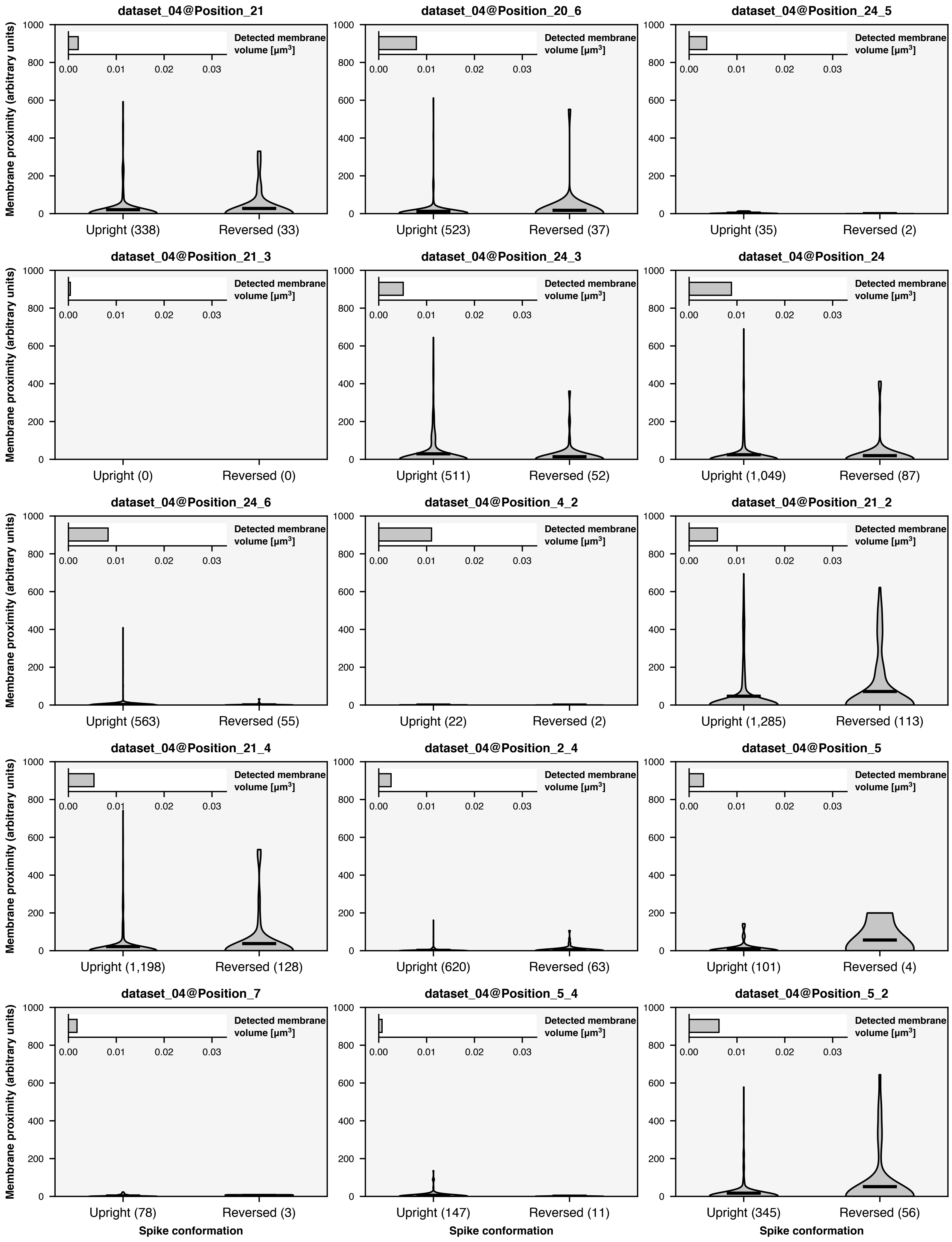
dataset_04@Position_21
Upright (338)
Reversed (33)
dataset_04@Position_20_6
Upright (523)
Reversed (37)
dataset_04@Position_24_5
Upright (35)
Reversed (2)
dataset_04@Position_21_3
Upright (0)
Reversed (0)
dataset_04@Position_24_3
Upright (511)
Reversed (52)
dataset_04@Position_24
Upright (1,049)
Reversed (87)
dataset_04@Position_24_6
Upright (563)
Reversed (55)
dataset_04@Position_4_2
Upright (22)
Reversed (2)
dataset_04@Position_21_2
Upright (1,285)
Reversed (113)
dataset_04@Position_21_4
Upright (1,198)
Reversed (128)
dataset_04@Position_2_4
Upright (620)
Reversed (63)
dataset_04@Position_5
Upright (101)
Reversed (4)
dataset_04@Position_7
Upright (78)
Reversed (3)
dataset_04@Position_5_4
Upright (147)
Reversed (11)
dataset_04@Position_5_2
Upright (345)
Reversed (56)
Membrane proximity (arbitrary units)
Detected membrane volume [μm³]
Spike conformation
0
200
400
600
800
1000
0.00
0.01
0.02
0.03

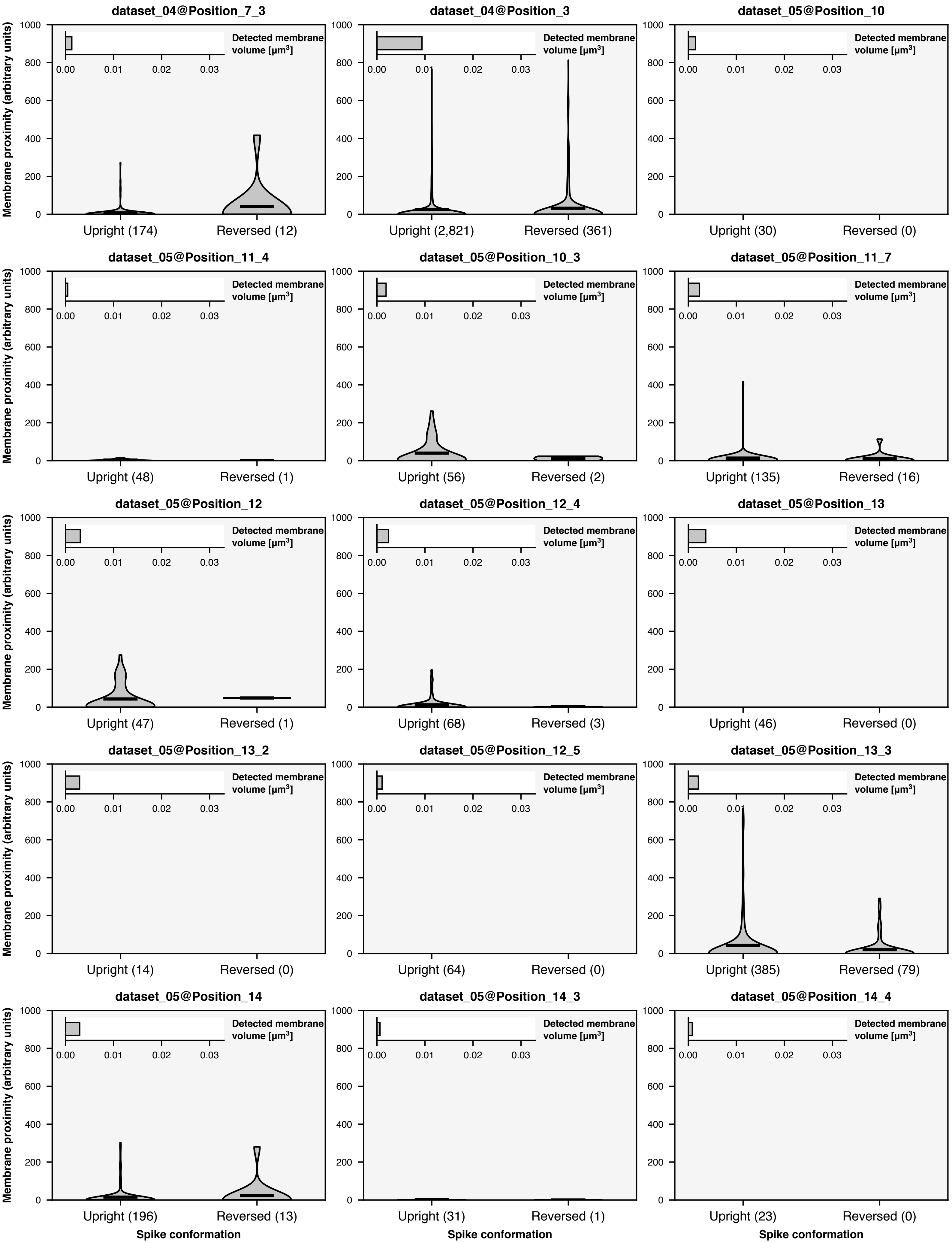

Membrane proximity (arbitrary units)
Detected membrane volume [µm³]
0.00 0.01 0.02 0.03
0 200 400 600 800 1000
dataset_04@Position_7_3
Upright (174)
Reversed (12)
dataset_04@Position_3
Upright (2,821)
Reversed (361)
dataset_05@Position_10
Upright (30)
Reversed (0)
dataset_05@Position_11_4
Upright (48)
Reversed (1)
dataset_05@Position_10_3
Upright (56)
Reversed (2)
dataset_05@Position_11_7
Upright (135)
Reversed (16)
dataset_05@Position_12
Upright (47)
Reversed (1)
dataset_05@Position_12_4
Upright (68)
Reversed (3)
dataset_05@Position_13
Upright (46)
Reversed (0)
dataset_05@Position_13_2
Upright (14)
Reversed (0)
dataset_05@Position_12_5
Upright (64)
Reversed (0)
dataset_05@Position_13_3
Upright (385)
Reversed (79)
dataset_05@Position_14
Upright (196)
Reversed (13)
dataset_05@Position_14_3
Upright (31)
Reversed (1)
dataset_05@Position_14_4
Upright (23)
Reversed (0)
Spike conformation

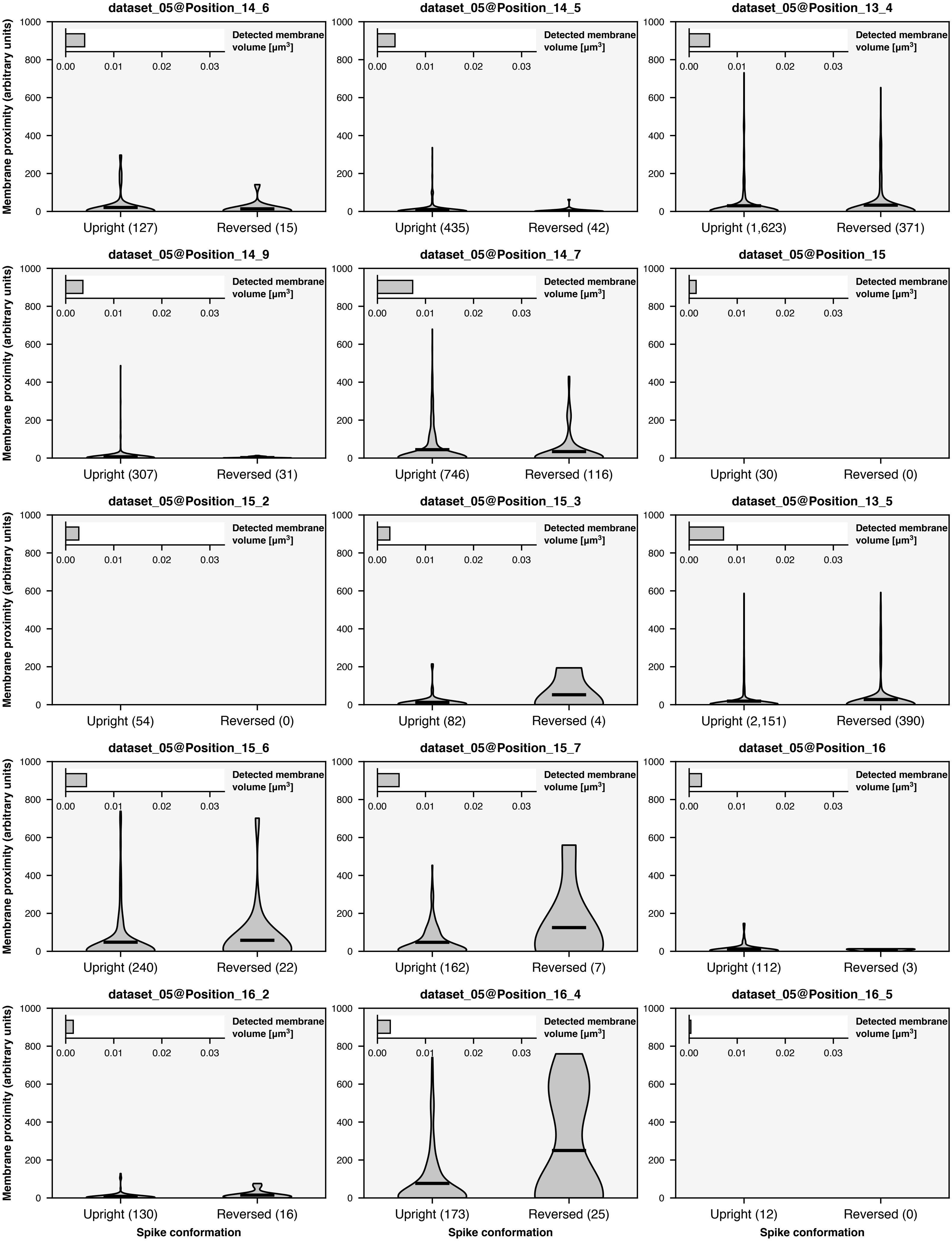
dataset_05@Position_14_6
dataset_05@Position_14_5
dataset_05@Position_13_4
dataset_05@Position_14_9
dataset_05@Position_14_7
dataset_05@Position_15
dataset_05@Position_15_2
dataset_05@Position_15_3
dataset_05@Position_13_5
dataset_05@Position_15_6
dataset_05@Position_15_7
dataset_05@Position_16
dataset_05@Position_16_2
dataset_05@Position_16_4
dataset_05@Position_16_5
Membrane proximity (arbitrary units)
Detected membrane volume [μm³]
Upright (127)
Reversed (15)
Upright (435)
Reversed (42)
Upright (1,623)
Reversed (371)
Upright (307)
Reversed (31)
Upright (746)
Reversed (116)
Upright (30)
Reversed (0)
Upright (54)
Reversed (0)
Upright (82)
Reversed (4)
Upright (2,151)
Reversed (390)
Upright (240)
Reversed (22)
Upright (162)
Reversed (7)
Upright (112)
Reversed (3)
Upright (130)
Reversed (16)
Upright (173)
Reversed (25)
Upright (12)
Reversed (0)
Spike conformation

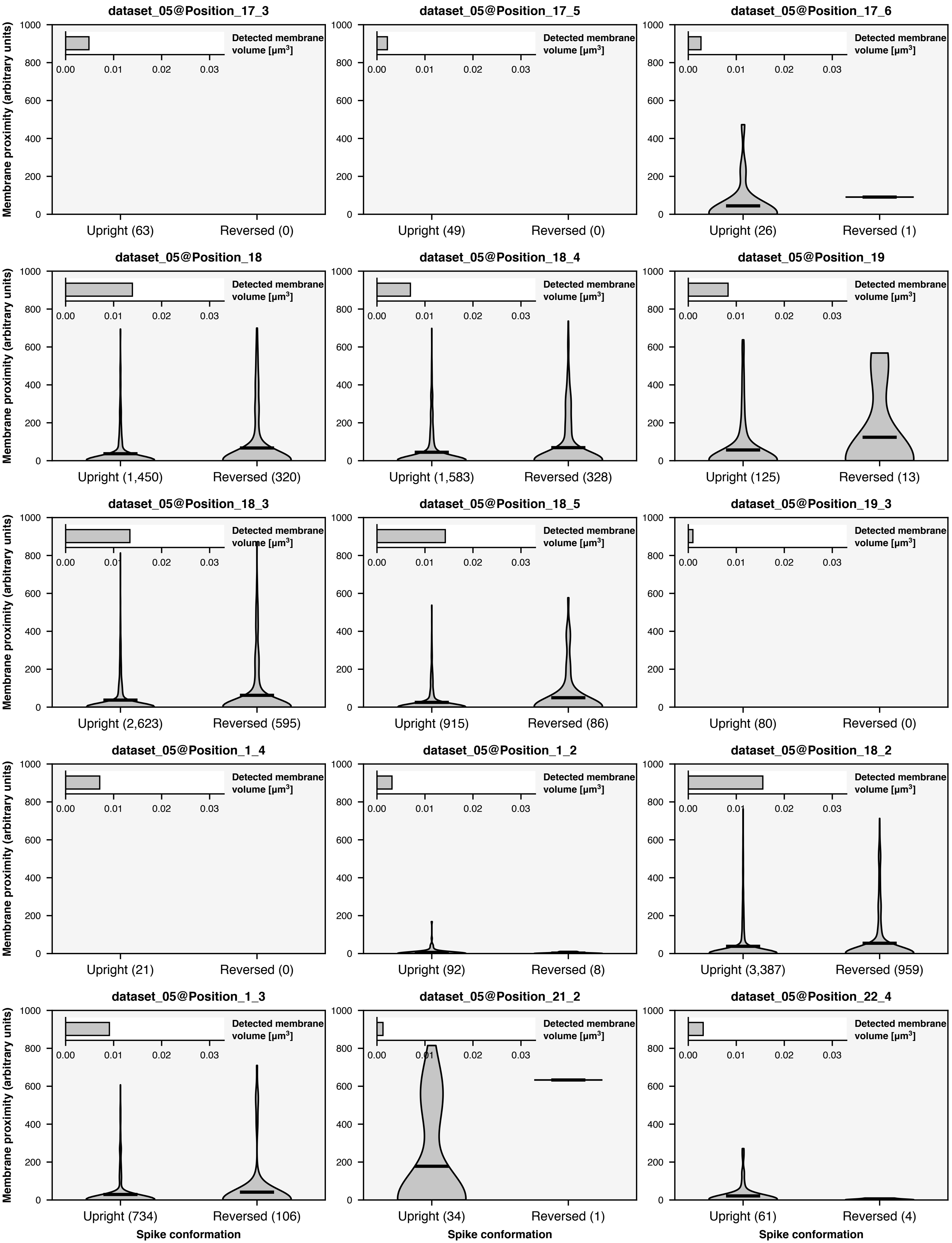
dataset_05@Position_17_3
dataset_05@Position_17_5
dataset_05@Position_17_6
dataset_05@Position_18
dataset_05@Position_18_4
dataset_05@Position_19
dataset_05@Position_18_3
dataset_05@Position_18_5
dataset_05@Position_19_3
dataset_05@Position_1_4
dataset_05@Position_1_2
dataset_05@Position_18_2
dataset_05@Position_1_3
dataset_05@Position_21_2
dataset_05@Position_22_4
Membrane proximity (arbitrary units)
Detected membrane volume [μm³]
Upright (63) Reversed (0)
Upright (49) Reversed (0)
Upright (26) Reversed (1)
Upright (1,450) Reversed (320)
Upright (1,583) Reversed (328)
Upright (125) Reversed (13)
Upright (2,623) Reversed (595)
Upright (915) Reversed (86)
Upright (80) Reversed (0)
Upright (21) Reversed (0)
Upright (92) Reversed (8)
Upright (3,387) Reversed (959)
Upright (734) Reversed (106)
Upright (34) Reversed (1)
Upright (61) Reversed (4)
Spike conformation

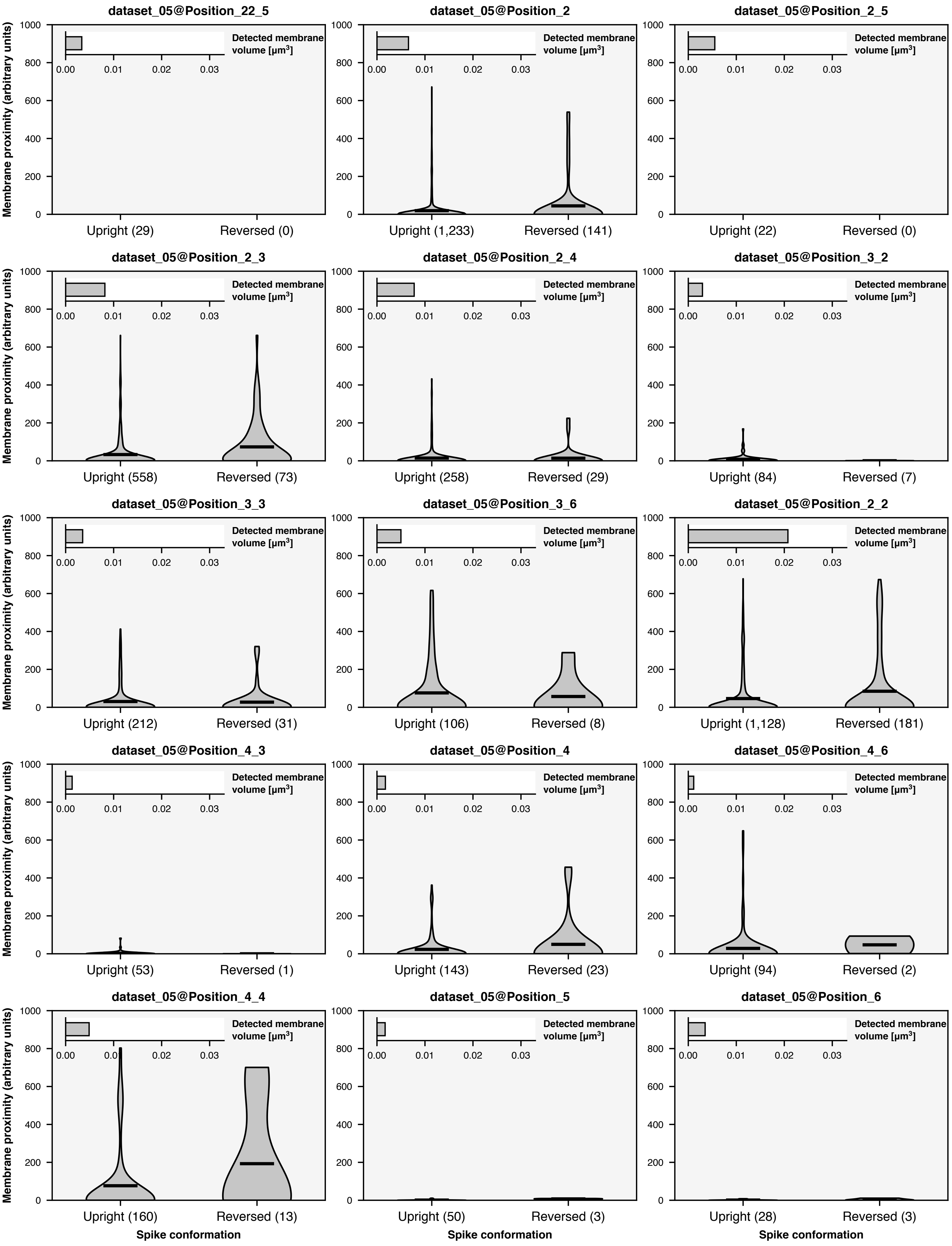

dataset_05@Position_22_5
Upright (29)
Reversed (0)
dataset_05@Position_2
Upright (1,233)
Reversed (141)
dataset_05@Position_2_5
Upright (22)
Reversed (0)
dataset_05@Position_2_3
Upright (558)
Reversed (73)
dataset_05@Position_2_4
Upright (258)
Reversed (29)
dataset_05@Position_3_2
Upright (84)
Reversed (7)
dataset_05@Position_3_3
Upright (212)
Reversed (31)
dataset_05@Position_3_6
Upright (106)
Reversed (8)
dataset_05@Position_2_2
Upright (1,128)
Reversed (181)
dataset_05@Position_4_3
Upright (53)
Reversed (1)
dataset_05@Position_4
Upright (143)
Reversed (23)
dataset_05@Position_4_6
Upright (94)
Reversed (2)
dataset_05@Position_4_4
Upright (160)
Reversed (13)
dataset_05@Position_5
Upright (50)
Reversed (3)
dataset_05@Position_6
Upright (28)
Reversed (3)
Membrane proximity (arbitrary units)
Detected membrane volume [μm³]
0.00
0.01
0.02
0.03
0
200
400
600
800
1000
Spike conformation

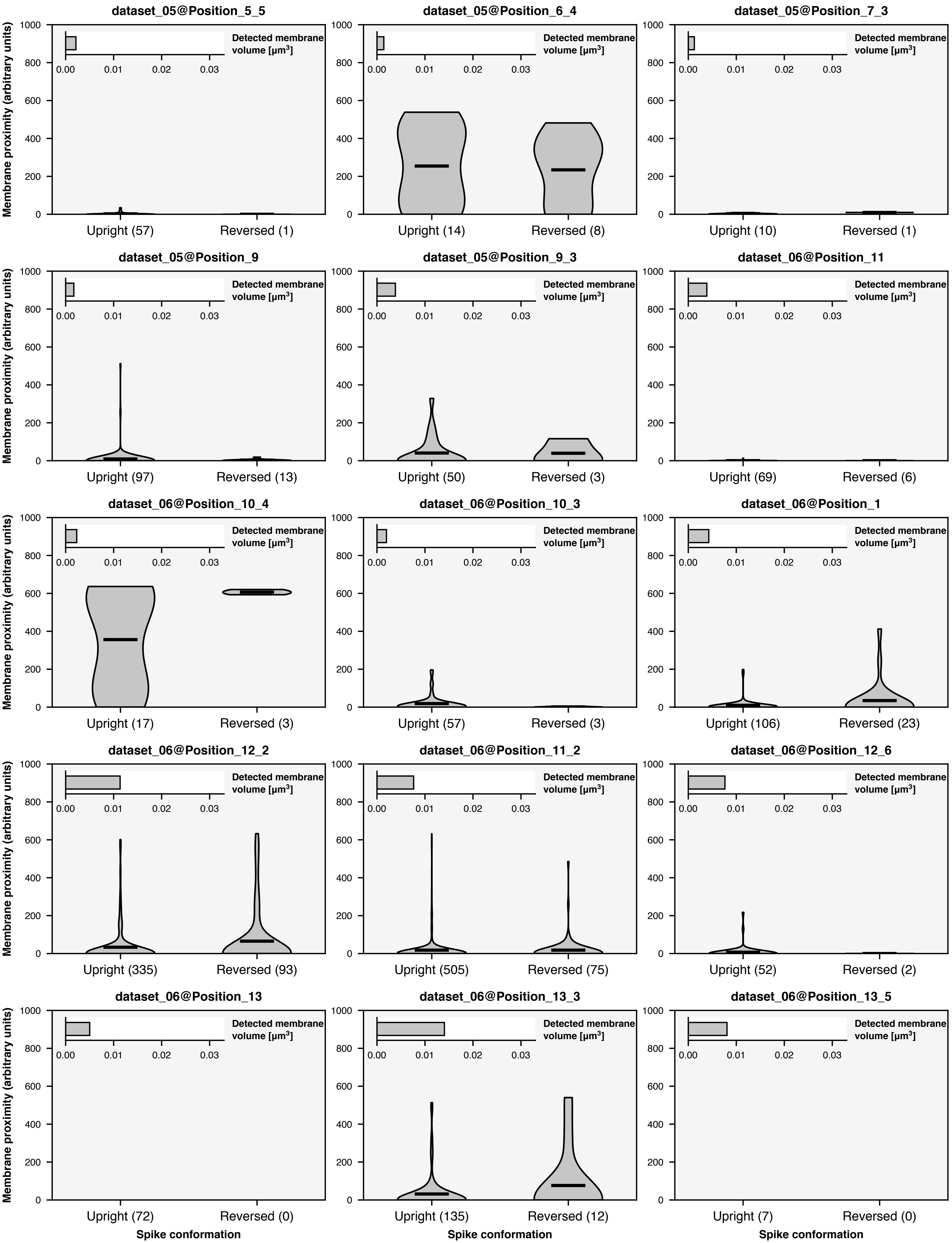
dataset_05@Position_5_5
Membrane proximity (arbitrary units)
Detected membrane volume [μm³]
Upright (57)
Reversed (1)
dataset_05@Position_6_4
Detected membrane volume [μm³]
Upright (14)
Reversed (8)
dataset_05@Position_7_3
Detected membrane volume [μm³]
Upright (10)
Reversed (1)
dataset_05@Position_9
Membrane proximity (arbitrary units)
Detected membrane volume [μm³]
Upright (97)
Reversed (13)
dataset_05@Position_9_3
Detected membrane volume [μm³]
Upright (50)
Reversed (3)
dataset_06@Position_11
Detected membrane volume [μm³]
Upright (69)
Reversed (6)
dataset_06@Position_10_4
Membrane proximity (arbitrary units)
Detected membrane volume [μm³]
Upright (17)
Reversed (3)
dataset_06@Position_10_3
Detected membrane volume [μm³]
Upright (57)
Reversed (3)
dataset_06@Position_1
Detected membrane volume [μm³]
Upright (106)
Reversed (23)
dataset_06@Position_12_2
Membrane proximity (arbitrary units)
Detected membrane volume [μm³]
Upright (335)
Reversed (93)
dataset_06@Position_11_2
Detected membrane volume [μm³]
Upright (505)
Reversed (75)
dataset_06@Position_12_6
Detected membrane volume [μm³]
Upright (52)
Reversed (2)
dataset_06@Position_13
Membrane proximity (arbitrary units)
Detected membrane volume [μm³]
Upright (72)
Reversed (0)
Spike conformation
dataset_06@Position_13_3
Detected membrane volume [μm³]
Upright (135)
Reversed (12)
Spike conformation
dataset_06@Position_13_5
Detected membrane volume [μm³]
Upright (7)
Reversed (0)
Spike conformation

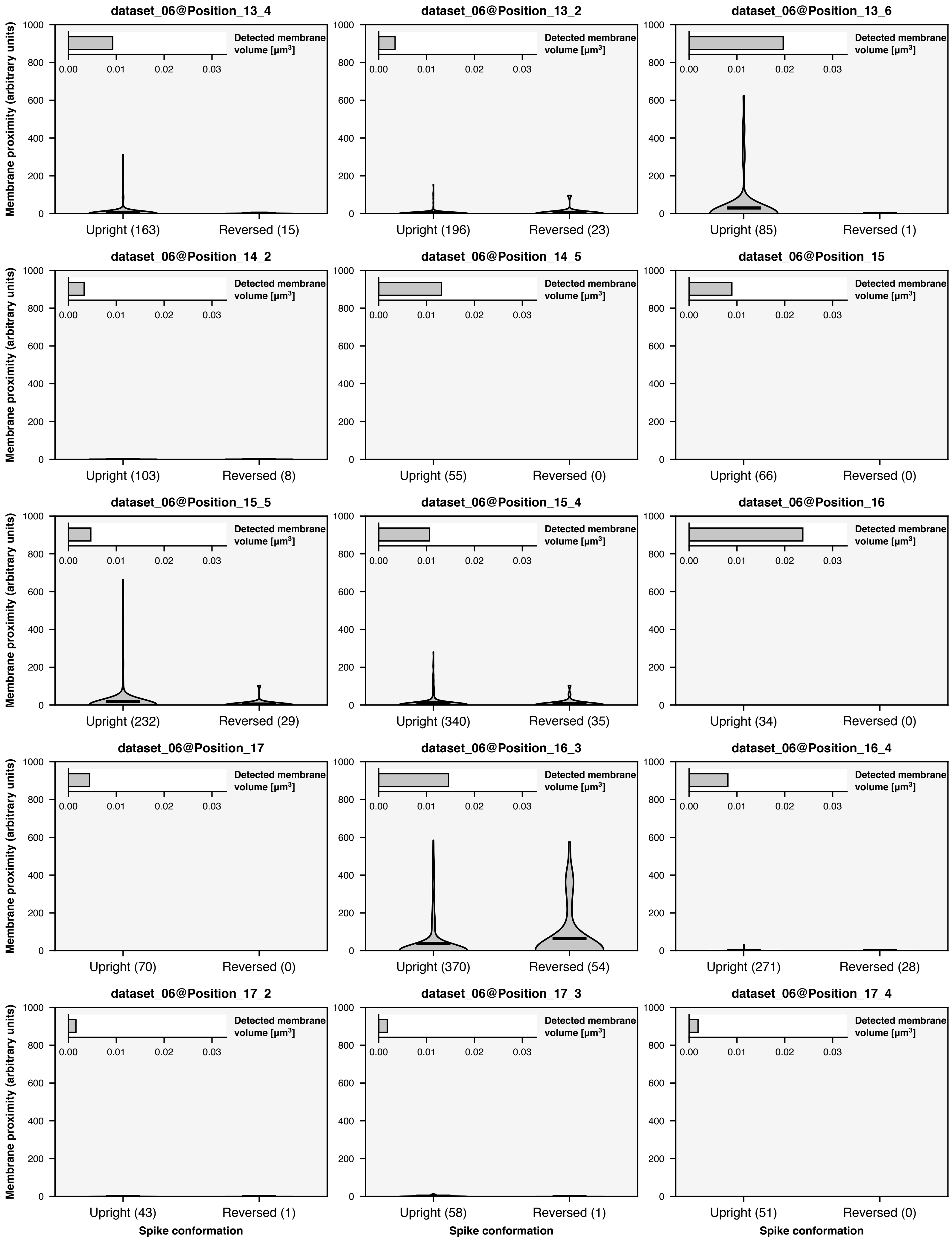
dataset_06@Position_13_4
dataset_06@Position_13_2
dataset_06@Position_13_6
dataset_06@Position_14_2
dataset_06@Position_14_5
dataset_06@Position_15
dataset_06@Position_15_5
dataset_06@Position_15_4
dataset_06@Position_16
dataset_06@Position_17
dataset_06@Position_16_3
dataset_06@Position_16_4
dataset_06@Position_17_2
dataset_06@Position_17_3
dataset_06@Position_17_4
Membrane proximity (arbitrary units)
Detected membrane volume [μm$^3$]
Upright (163)
Reversed (15)
Upright (196)
Reversed (23)
Upright (85)
Reversed (1)
Upright (103)
Reversed (8)
Upright (55)
Reversed (0)
Upright (66)
Reversed (0)
Upright (232)
Reversed (29)
Upright (340)
Reversed (35)
Upright (34)
Reversed (0)
Upright (70)
Reversed (0)
Upright (370)
Reversed (54)
Upright (271)
Reversed (28)
Upright (43)
Reversed (1)
Upright (58)
Reversed (1)
Upright (51)
Reversed (0)
Spike conformation

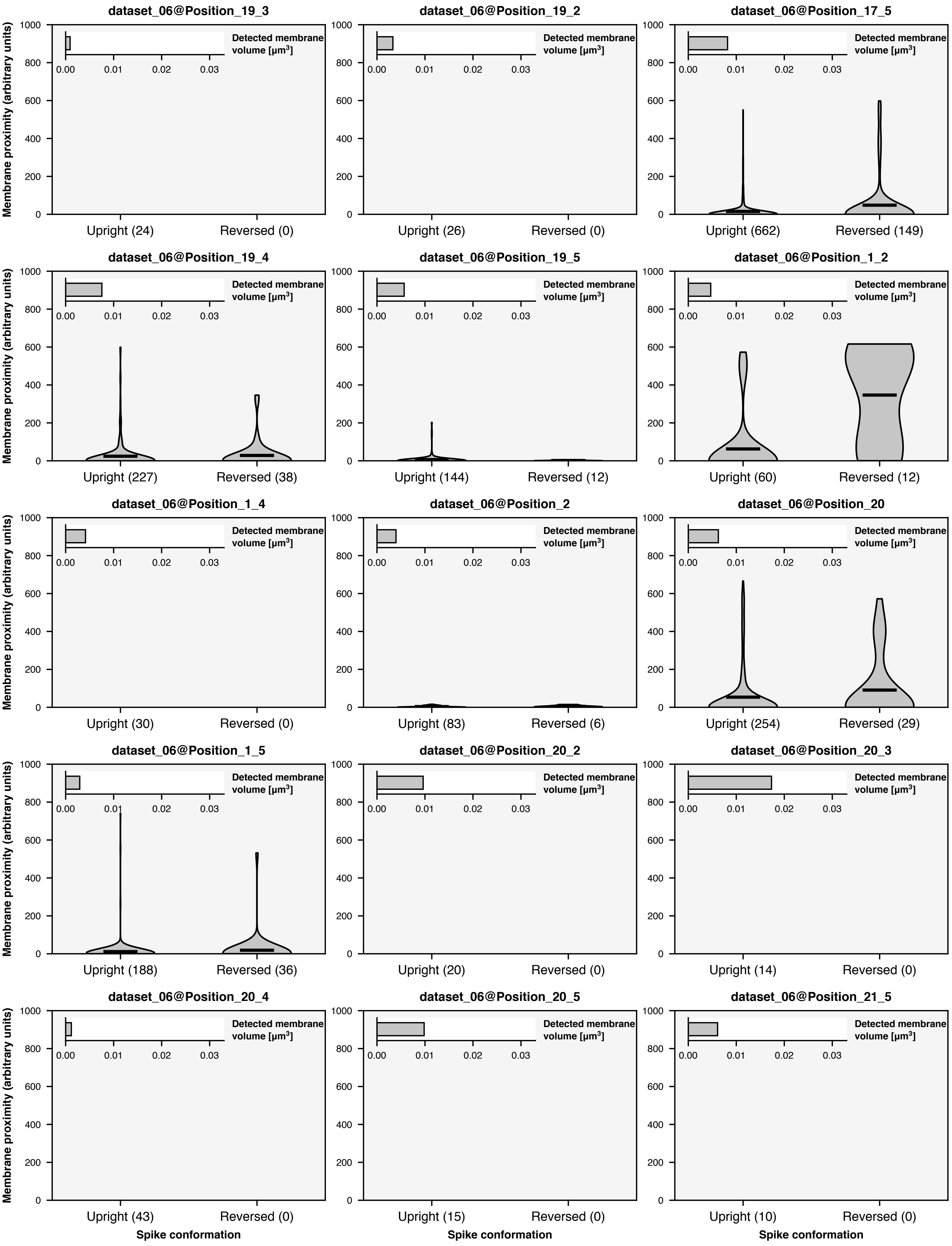
dataset_06@Position_19_3
dataset_06@Position_19_2
dataset_06@Position_17_5
Membrane proximity (arbitrary units)
Detected membrane volume [μm³]
0.00
0.01
0.02
0.03
0
200
400
600
800
1000
Upright (24)
Reversed (0)
Upright (26)
Reversed (0)
Upright (662)
Reversed (149)
dataset_06@Position_19_4
dataset_06@Position_19_5
dataset_06@Position_1_2
Upright (227)
Reversed (38)
Upright (144)
Reversed (12)
Upright (60)
Reversed (12)
dataset_06@Position_1_4
dataset_06@Position_2
dataset_06@Position_20
Upright (30)
Reversed (0)
Upright (83)
Reversed (6)
Upright (254)
Reversed (29)
dataset_06@Position_1_5
dataset_06@Position_20_2
dataset_06@Position_20_3
Upright (188)
Reversed (36)
Upright (20)
Reversed (0)
Upright (14)
Reversed (0)
dataset_06@Position_20_4
dataset_06@Position_20_5
dataset_06@Position_21_5
Upright (43)
Reversed (0)
Upright (15)
Reversed (0)
Upright (10)
Reversed (0)
Spike conformation

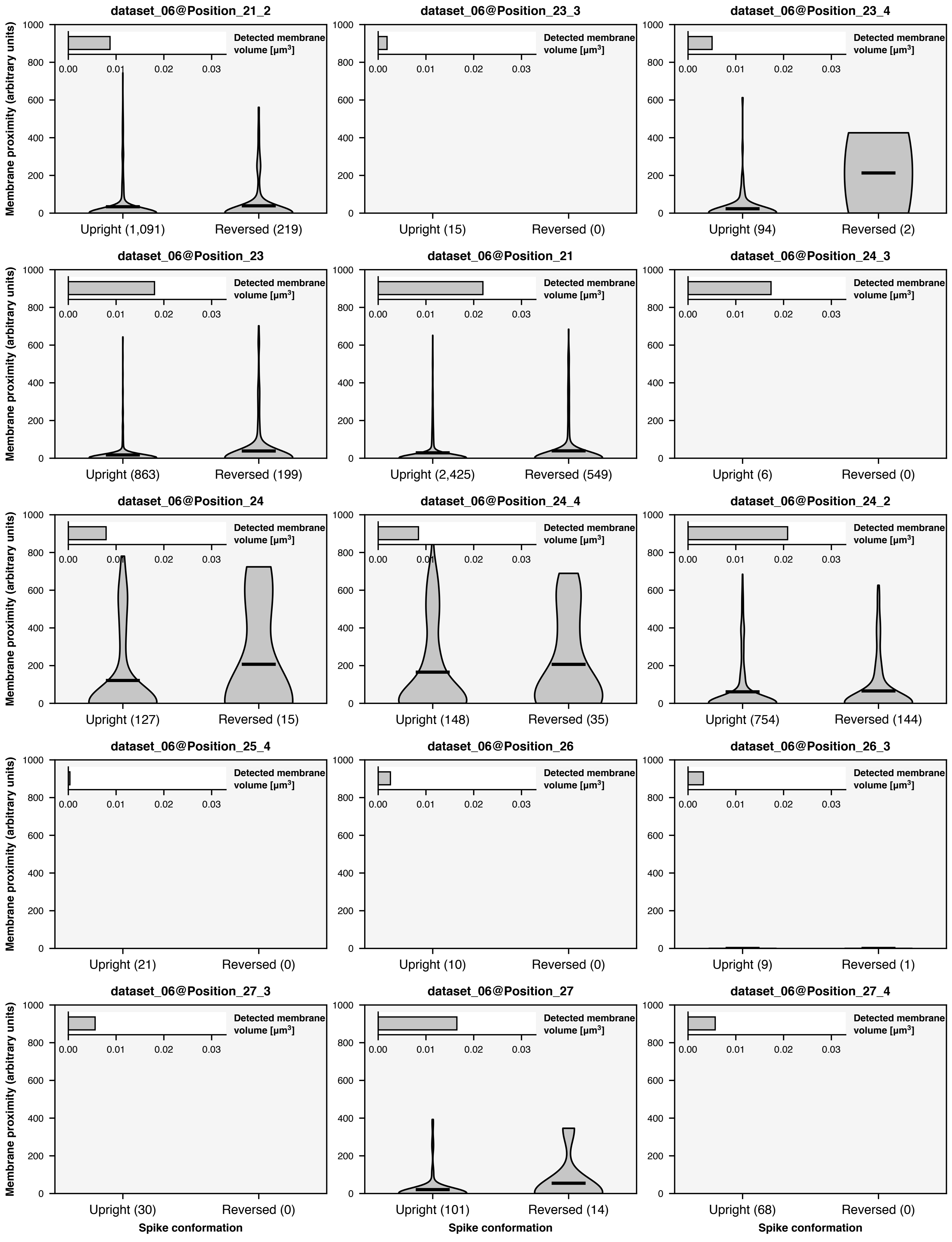
dataset_06@Position_21_2
Upright (1,091)
Reversed (219)
dataset_06@Position_23_3
Upright (15)
Reversed (0)
dataset_06@Position_23_4
Upright (94)
Reversed (2)
dataset_06@Position_23
Upright (863)
Reversed (199)
dataset_06@Position_21
Upright (2,425)
Reversed (549)
dataset_06@Position_24_3
Upright (6)
Reversed (0)
dataset_06@Position_24
Upright (127)
Reversed (15)
dataset_06@Position_24_4
Upright (148)
Reversed (35)
dataset_06@Position_24_2
Upright (754)
Reversed (144)
dataset_06@Position_25_4
Upright (21)
Reversed (0)
dataset_06@Position_26
Upright (10)
Reversed (0)
dataset_06@Position_26_3
Upright (9)
Reversed (1)
dataset_06@Position_27_3
Upright (30)
Reversed (0)
dataset_06@Position_27
Upright (101)
Reversed (14)
dataset_06@Position_27_4
Upright (68)
Reversed (0)
Membrane proximity (arbitrary units)
Detected membrane volume [μm³]
0.00
0.01
0.02
0.03
0
200
400
600
800
1000
Spike conformation

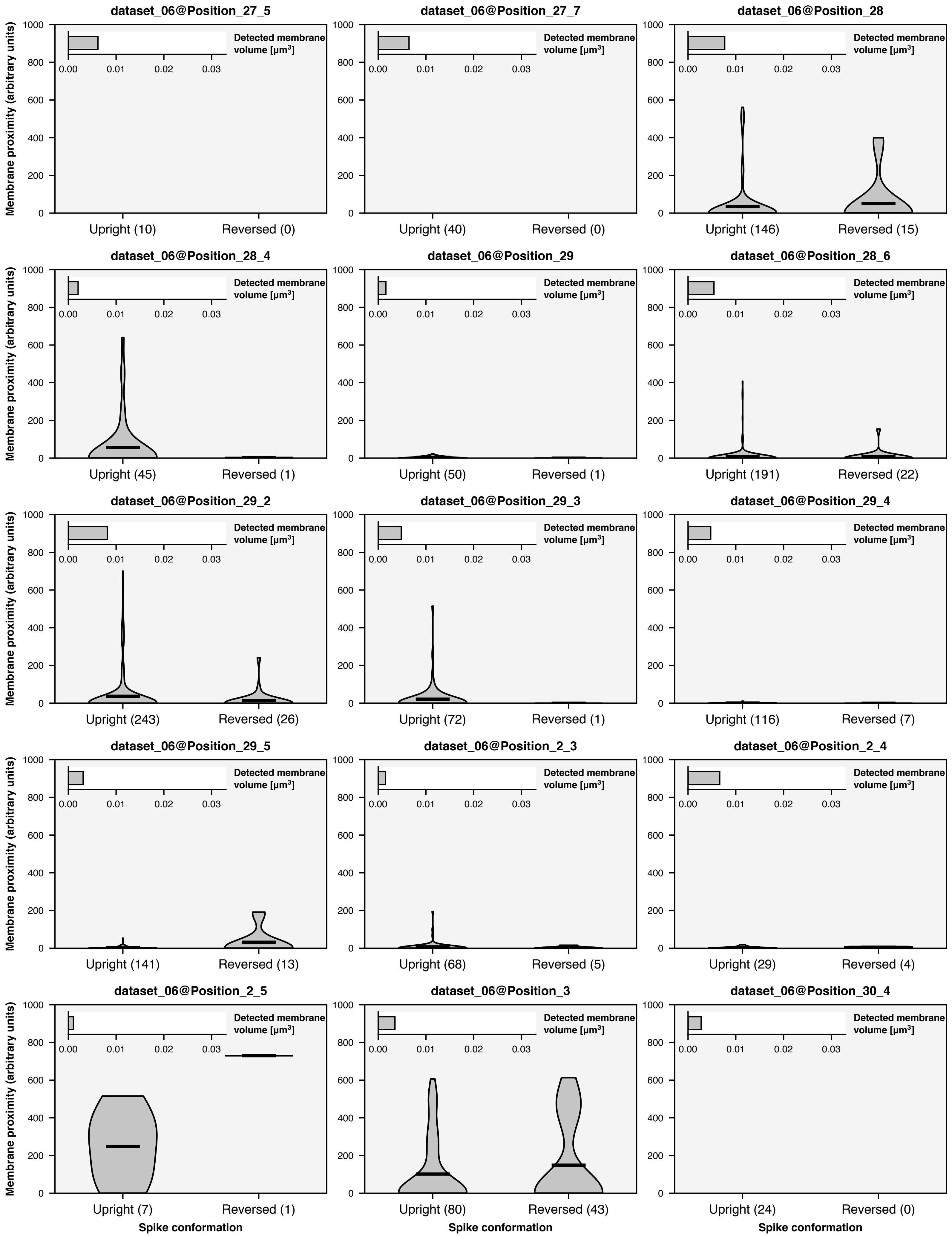

dataset_06@Position_27_5
dataset_06@Position_27_7
dataset_06@Position_28
dataset_06@Position_28_4
dataset_06@Position_29
dataset_06@Position_28_6
dataset_06@Position_29_2
dataset_06@Position_29_3
dataset_06@Position_29_4
dataset_06@Position_29_5
dataset_06@Position_2_3
dataset_06@Position_2_4
dataset_06@Position_2_5
dataset_06@Position_3
dataset_06@Position_30_4
Membrane proximity (arbitrary units)
Detected membrane volume [μm3]
Upright (10)
Reversed (0)
Upright (40)
Reversed (0)
Upright (146)
Reversed (15)
Upright (45)
Reversed (1)
Upright (50)
Reversed (1)
Upright (191)
Reversed (22)
Upright (243)
Reversed (26)
Upright (72)
Reversed (1)
Upright (116)
Reversed (7)
Upright (141)
Reversed (13)
Upright (68)
Reversed (5)
Upright (29)
Reversed (4)
Upright (7)
Reversed (1)
Upright (80)
Reversed (43)
Upright (24)
Reversed (0)
Spike conformation

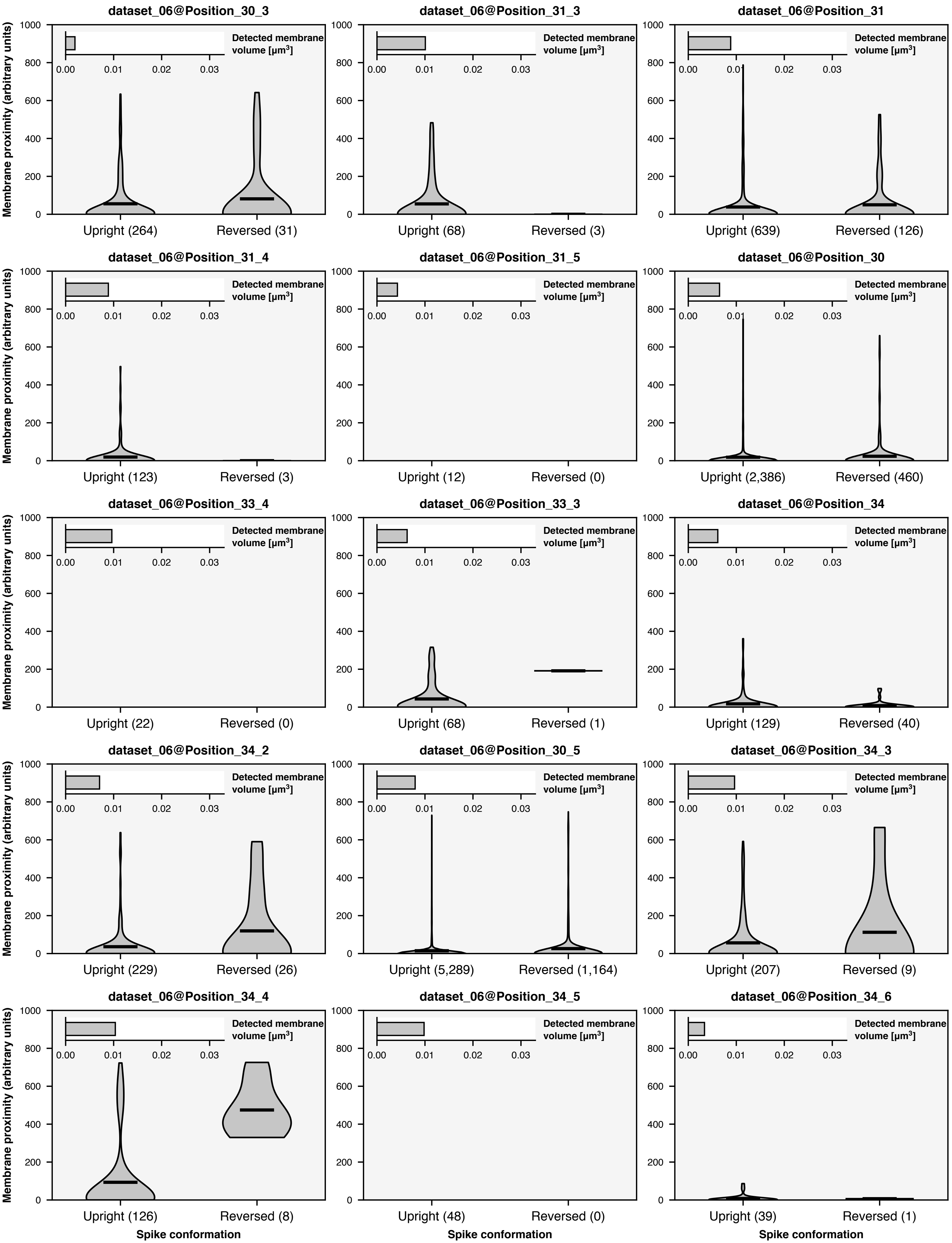
dataset_06@Position_30_3
dataset_06@Position_31_3
dataset_06@Position_31
dataset_06@Position_31_4
dataset_06@Position_31_5
dataset_06@Position_30
dataset_06@Position_33_4
dataset_06@Position_33_3
dataset_06@Position_34
dataset_06@Position_34_2
dataset_06@Position_30_5
dataset_06@Position_34_3
dataset_06@Position_34_4
dataset_06@Position_34_5
dataset_06@Position_34_6
Membrane proximity (arbitrary units)
Detected membrane volume [μm³]
0.00
0.01
0.02
0.03
0
200
400
600
800
1000
Upright (264)
Reversed (31)
Upright (68)
Reversed (3)
Upright (639)
Reversed (126)
Upright (123)
Reversed (3)
Upright (12)
Reversed (0)
Upright (2,386)
Reversed (460)
Upright (22)
Reversed (0)
Upright (68)
Reversed (1)
Upright (129)
Reversed (40)
Upright (229)
Reversed (26)
Upright (5,289)
Reversed (1,164)
Upright (207)
Reversed (9)
Upright (126)
Reversed (8)
Upright (48)
Reversed (0)
Upright (39)
Reversed (1)
Spike conformation

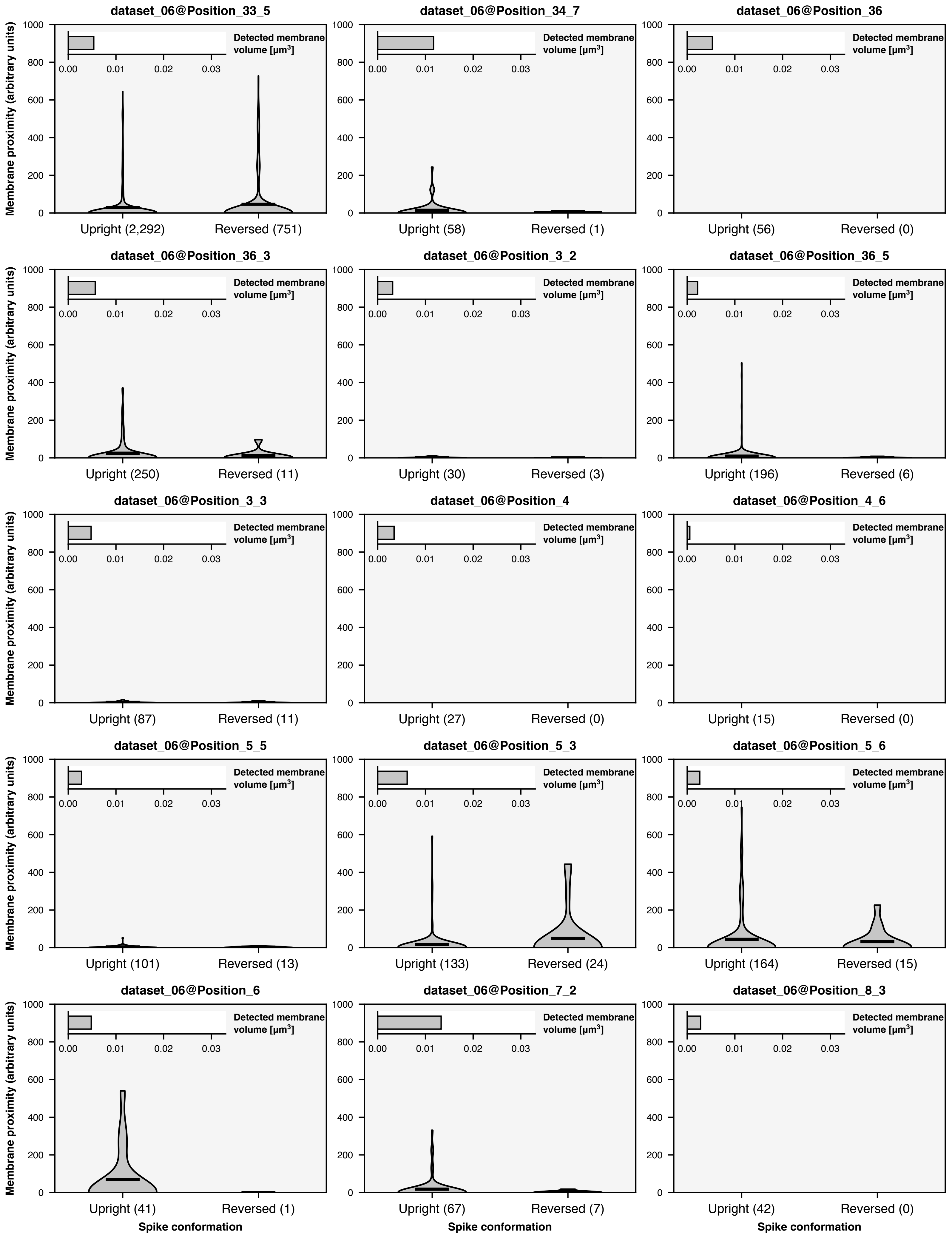

dataset_06@Position_33_5
dataset_06@Position_34_7
dataset_06@Position_36
dataset_06@Position_36_3
dataset_06@Position_3_2
dataset_06@Position_36_5
dataset_06@Position_3_3
dataset_06@Position_4
dataset_06@Position_4_6
dataset_06@Position_5_5
dataset_06@Position_5_3
dataset_06@Position_5_6
dataset_06@Position_6
dataset_06@Position_7_2
dataset_06@Position_8_3
Membrane proximity (arbitrary units)
Detected membrane volume [µm³]
0.00
0.01
0.02
0.03
0
200
400
600
800
1000
Upright (2,292)
Reversed (751)
Upright (58)
Reversed (1)
Upright (56)
Reversed (0)
Upright (250)
Reversed (11)
Upright (30)
Reversed (3)
Upright (196)
Reversed (6)
Upright (87)
Reversed (11)
Upright (27)
Reversed (0)
Upright (15)
Reversed (0)
Upright (101)
Reversed (13)
Upright (133)
Reversed (24)
Upright (164)
Reversed (15)
Upright (41)
Reversed (1)
Upright (67)
Reversed (7)
Upright (42)
Reversed (0)
Spike conformation

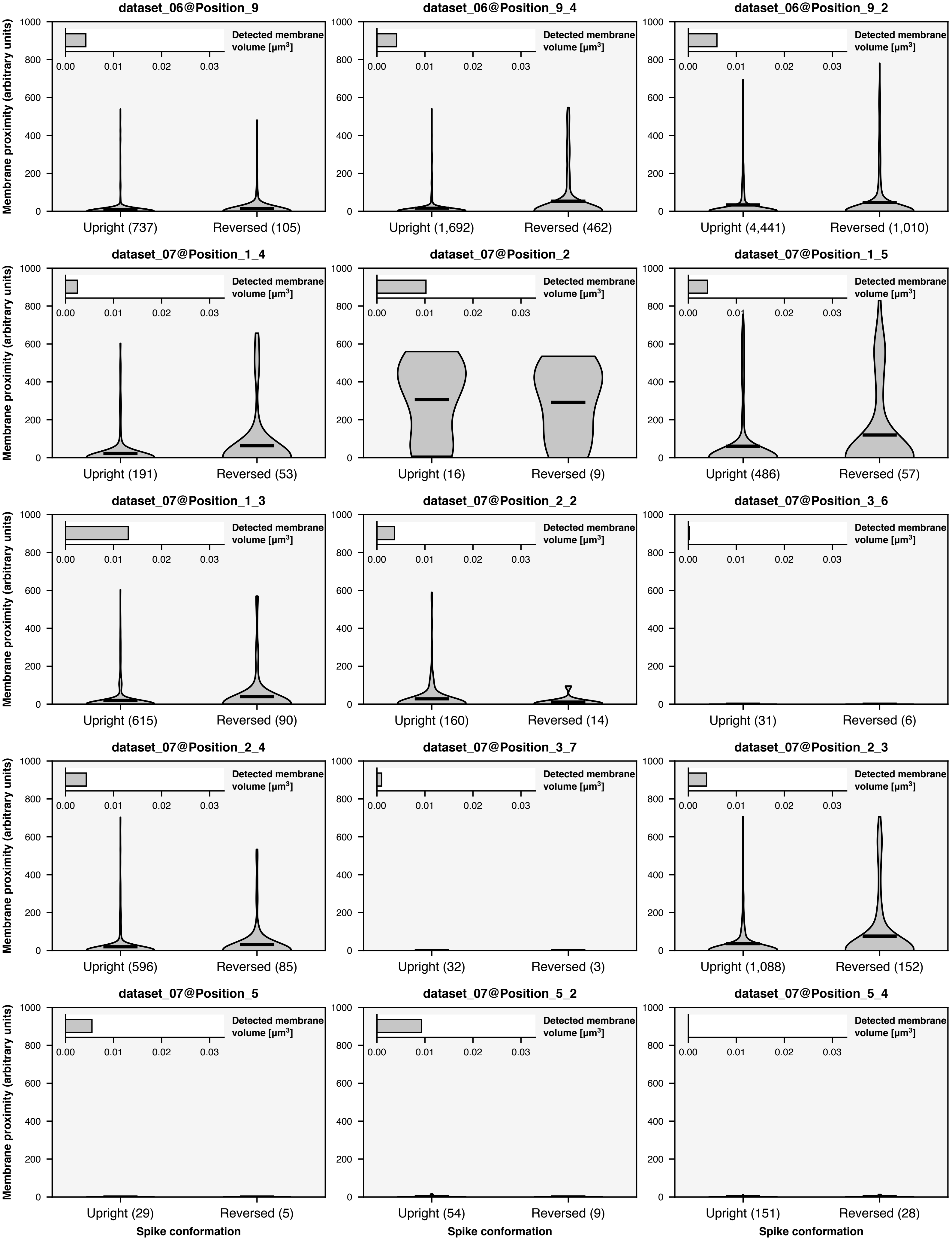

dataset_06@Position_9
Upright (737)
Reversed (105)
dataset_06@Position_9_4
Upright (1,692)
Reversed (462)
dataset_06@Position_9_2
Upright (4,441)
Reversed (1,010)
dataset_07@Position_1_4
Upright (191)
Reversed (53)
dataset_07@Position_2
Upright (16)
Reversed (9)
dataset_07@Position_1_5
Upright (486)
Reversed (57)
dataset_07@Position_1_3
Upright (615)
Reversed (90)
dataset_07@Position_2_2
Upright (160)
Reversed (14)
dataset_07@Position_3_6
Upright (31)
Reversed (6)
dataset_07@Position_2_4
Upright (596)
Reversed (85)
dataset_07@Position_3_7
Upright (32)
Reversed (3)
dataset_07@Position_2_3
Upright (1,088)
Reversed (152)
dataset_07@Position_5
Upright (29)
Reversed (5)
dataset_07@Position_5_2
Upright (54)
Reversed (9)
dataset_07@Position_5_4
Upright (151)
Reversed (28)
Detected membrane volume [μm3]
0.00 0.01 0.02 0.03
Membrane proximity (arbitrary units)
0 200 400 600 800 1000
Spike conformation

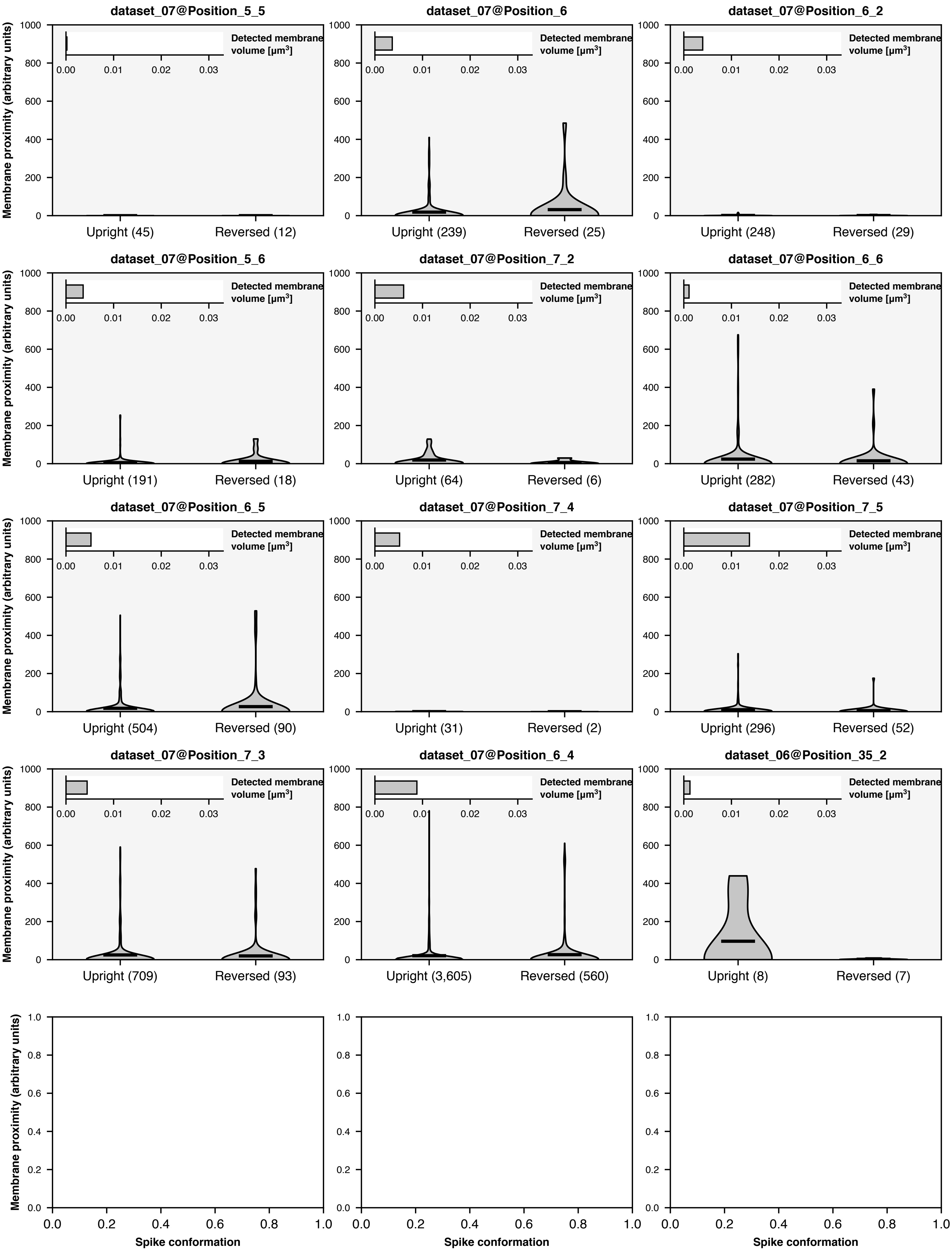

dataset_07@Position_5_5
dataset_07@Position_6
dataset_07@Position_6_2
dataset_07@Position_5_6
dataset_07@Position_7_2
dataset_07@Position_6_6
dataset_07@Position_6_5
dataset_07@Position_7_4
dataset_07@Position_7_5
dataset_07@Position_7_3
dataset_07@Position_6_4
dataset_06@Position_35_2
Membrane proximity (arbitrary units)
Detected membrane volume [μm³]
Upright (45)
Reversed (12)
Upright (239)
Reversed (25)
Upright (248)
Reversed (29)
Upright (191)
Reversed (18)
Upright (64)
Reversed (6)
Upright (282)
Reversed (43)
Upright (504)
Reversed (90)
Upright (31)
Reversed (2)
Upright (296)
Reversed (52)
Upright (709)
Reversed (93)
Upright (3,605)
Reversed (560)
Upright (8)
Reversed (7)
Spike conformation
